# Functional connectomics reveals general wiring rule in mouse visual cortex

**DOI:** 10.1101/2023.03.13.531369

**Authors:** Zhuokun Ding, Paul G. Fahey, Stelios Papadopoulos, Eric Y. Wang, Brendan Celii, Christos Papadopoulos, Andersen Chang, Alexander B. Kunin, Dat Tran, Jiakun Fu, Zhiwei Ding, Saumil Patel, Lydia Ntanavara, Rachel Froebe, Kayla Ponder, Taliah Muhammad, J. Alexander Bae, Agnes L. Bodor, Derrick Brittain, JoAnn Buchanan, Daniel J. Bumbarger, Manuel A. Castro, Erick Cobos, Sven Dorkenwald, Leila Elabbady, Akhilesh Halageri, Zhen Jia, Chris Jordan, Dan Kapner, Nico Kemnitz, Sam Kinn, Kisuk Lee, Kai Li, Ran Lu, Thomas Macrina, Gayathri Mahalingam, Eric Mitchell, Shanka Subhra Mondal, Shang Mu, Barak Nehoran, Sergiy Popovych, Casey M. Schneider-Mizell, William Silversmith, Marc Takeno, Russel Torres, Nicholas L. Turner, William Wong, Jingpeng Wu, Wenjing Yin, Szi-chieh Yu, Dimitri Yatsenko, Emmanouil Froudarakis, Fabian Sinz, Krešimir Josić, Robert Rosenbaum, H. Sebastian Seung, Forrest Collman, Nuno Maçarico da Costa, R. Clay Reid, Edgar Y. Walker, Xaq Pitkow, Jacob Reimer, Andreas S. Tolias

## Abstract

Understanding the relationship between circuit connectivity and function is crucial for uncovering how the brain implements computation. In the mouse primary visual cortex (V1), excitatory neurons with similar response properties are more likely to be synaptically connected, but previous studies have been limited to within V1, leaving much unknown about broader connectivity rules. In this study, we leverage the millimeter-scale MICrONS dataset to analyze synaptic connectivity and functional properties of individual neurons across cortical layers and areas. Our results reveal that neurons with similar responses are preferentially connected both within and across layers and areas — including feedback connections — suggesting the universality of the ‘like-to-like’ connectivity across the visual hierarchy. Using a validated digital twin model, we separated neuronal tuning into feature (what neurons respond to) and spatial (receptive field location) components. We found that only the feature component predicts fine-scale synaptic connections, beyond what could be explained by the physical proximity of axons and dendrites. We also found a higher-order rule where postsynaptic neuron cohorts downstream of individual presynaptic cells show greater functional similarity than predicted by a pairwise like-to-like rule. Notably, recurrent neural networks (RNNs) trained on a simple classification task develop connectivity patterns mirroring both pairwise and higher-order rules, with magnitude similar to those in the MICrONS data. Lesion studies in these RNNs reveal that disrupting ‘like-to-like’ connections has a significantly greater impact on performance compared to lesions of random connections. These findings suggest that these connectivity principles may play a functional role in sensory processing and learning, highlighting shared principles between biological and artificial systems.

## Introduction

In the late 1800’s, Santiago Ramón y Cajal — while poring over the structure of Golgi-stained neurons using only light microscopy — imagined the Neuron Doctrine, the idea that individual neurons are the fundamental units of the nervous system (Ramón y Cajal, 1911). Implicit in the Neuron Doctrine is the idea that the function of individual neurons — their role in what we would now call neural computation — is inextricably linked to their connectivity in neural circuits. A variety of influential proposals about the relationship between connectivity and function have been advanced in the past century. For example, Donald Hebb’s cell assembly hypothesis (Hebb, 1949) — colloquially stated as “neurons that fire together, wire together” — predicted that interconnected neuronal subnetworks “reverberate” to stabilize functionally relevant activity patterns. In the cortical visual system, Hubel and Wiesel proposed that the hierarchical organization of connected neurons might build more complex feature preferences from simpler ones; for example the position invariance of orientation-selective complex cells might be derived from convergent inputs of like-oriented simple cells with spatially scattered receptive fields (Hubel and Wiesel, 1962; Reid, 2012).

Testing these predictions has been difficult because of the challenges of measuring neural activity and synaptic-scale connectivity in the same population of neurons. In the mammalian visual cortex, evidence for several varieties of like-to-like connectivity (i.e. increased connectivity for cells with similar response preferences) has been found via spine imaging (Iacaruso et al., 2017), combined *in vivo* imaging and *in vitro* multipatching (Ko et al., 2011, 2013; Cossell et al., 2015; Znamenskiy et al., 2024), combined *in vivo* imaging and rabies monosynaptic retrograde tracing (Wertz et al., 2015; Rossi et al., 2020), and combined *in vivo* imaging with electron microscopy (EM) reconstruction (Lee et al., 2016; Scholl et al., 2021). However, a caveat of these important early studies is that they have mostly been limited to small volumes, usually single lamina of primary visual cortex (except see Wertz et al. 2015; Rossi et al. 2020), mostly due to the challenge of identifying synaptic connections between functionally-characterized neurons across distances larger than a few hundred microns. Thus, many questions remain unanswered about how these rules generalize across areas and layers.

The MICrONS dataset is the largest functionally-imaged EM dataset to date (MICrONS Consortium et al., 2021), with mesoscopic calcium imaging (Sofroniew et al., 2016) performed *in vivo* and subsequent EM imaging (Yin et al., 2020; Phelps et al., 2021) and dense reconstruction (Turner et al., 2020; Dorkenwald et al., 2022b; Mitchell et al., 2019; Lu et al., 2021; Wu et al., 2021; Dorkenwald et al., 2022a; Lee et al., 2017) for an approximately 1 mm^3^ volume spanning visual cortical areas V1, LM, AL, and RL in a single mouse. In contrast with previous studies that have selectively reconstructed presynaptic or postsynaptic partners of a small set of functionally-characterized target cells (Lee et al., 2016; Bock et al., 2011), the MICrONS volume is densely reconstructed, offering access to segmentation of all neurons in the volume, and enabling analyses that are not possible in targeted sparse reconstructions. Here, we take advantage of the dense reconstruction to compare the functional similarity of connected pairs with unconnected “bystanders” — pairs of neurons with closely-apposed axons and dendrites that had the opportunity to form synaptic connections, yet didn’t.

Our analysis of functional similarity builds on recent advances in using machine learning to characterize the response properties of neurons in visual cortex. By training a neural network to replicate the responses of recorded neurons across a rich stimulus set of natural and parametric movies (Wang et al., 2024), we produce a “digital twin” of the cortical population which can accurately predict the response of a neuron to any arbitrary visual stimulus. The digital twin makes it possible to explore a much larger stimulus space with *in silico* experiments than would be possible (due to time constraints) with *in vivo* measurements (Wang et al., 2024). We have extensively validated this approach by looping back *in vivo* and validating model predictions of the most-exciting natural images and synthetic stimuli for a neuron (Walker et al., 2019). As part of the current study, we have validated the correspondence between model predictions and empirically-observed visual response properties, including signal correlations, orientation tuning, and spatial receptive field location. These validation results are described below. Finally, the digital twin model allowed us to separate each neuron’s tuning into two components: a feature component (**what** the neuron responded to), and a spatial component (**where** the neuron’s receptive field is located), allowing us to dissociate these two aspects of function and their relationship to connectivity.

## Results

### MICrONS functional connectomic dataset

Data were collected and processed as described in the MICrONS data release publication (MICrONS Consortium et al. 2021, Fig. 1). Briefly, a single mouse expressing GCaMP6s in excitatory neurons underwent fourteen two-photon scans (awake and headfixed on treadmill) of a 1200 × 1100 × 500 µm^3^ volume (anteroposterior × mediolateral × radial depth) spanning layers 2 through 6 at the conjunction of lateral primary visual cortex (V1) and anterolateral (AL), lateromedial (LM) and rostrolateral (RL) higher visual areas (Fig. 1a). Mice rapidly acclimated to head fixation, and were able to walk, groom, and adjust their posture during imaging. We monitored treadmill velocity and collected video of the pupil to track behavioral state. Neuronal responses from 115,372 functional units representing an estimated 75,909 unique excitatory neurons were collected in response to visual stimuli composed of natural and rendered movies and parametric dynamic stimuli (Fig. 1b). A state-of-the-art deep recurrent neural network was trained to predict neural responses to arbitrary stimuli (Wang et al., 2024), and used to characterize the *in silico* functional properties of imaged neurons (Fig. 1c).

**Figure 1.**
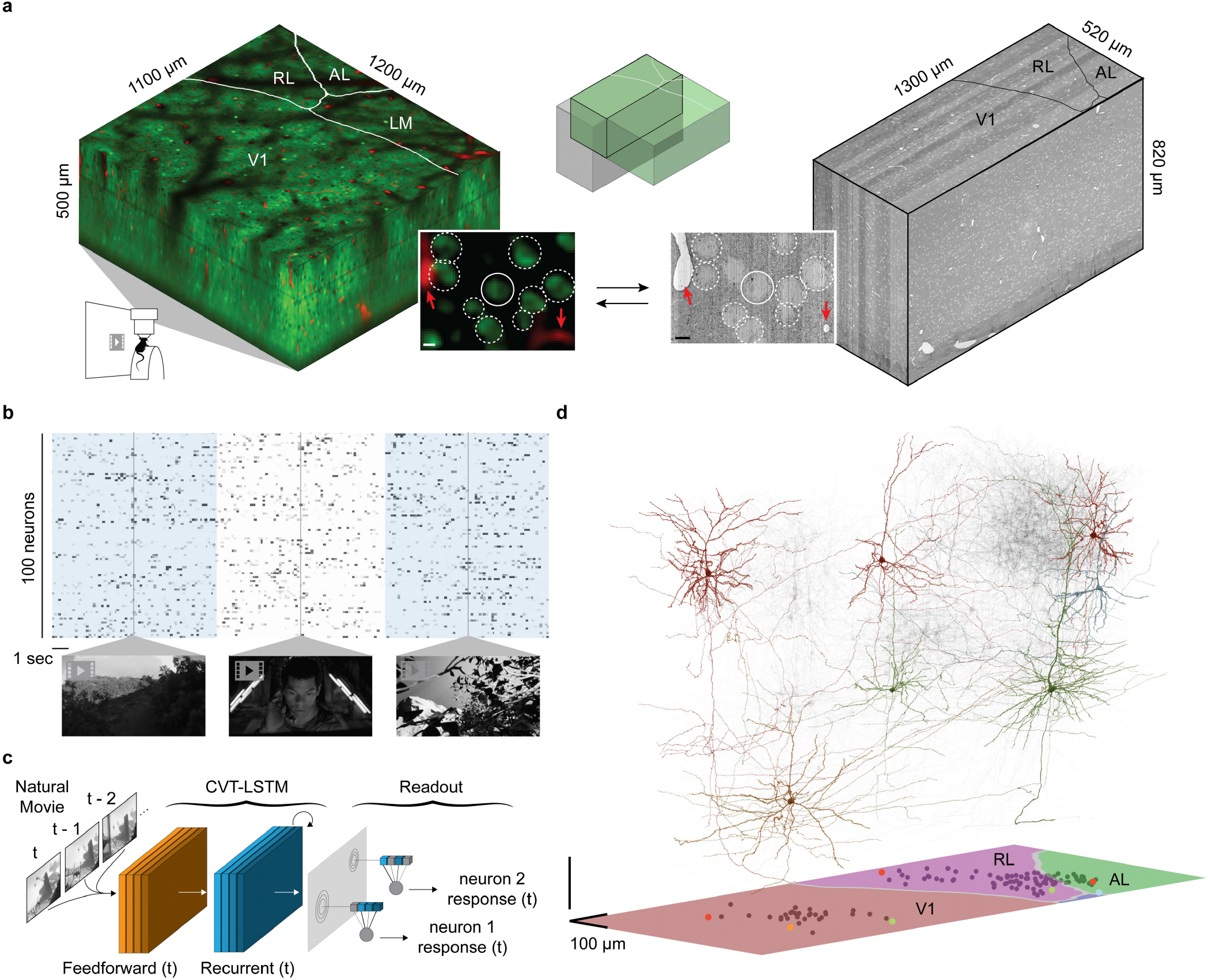
Overview of MICrONS Dataset. **a**, Depiction of functionally-characterized volumes (left; GCaMP6s in green, vascular label in red) and EM (right; gray). Visual areas: primary visual cortex (V1), anterolateral (AL), lateromedial (LM) and rostrolateral (RL).The overlap of the functional 2P (green) and structural EM (gray) volumes from which somas were recruited is depicted in the top inset. The bottom inset shows an example of matching structural features in the 2P and EM volumes, including a soma constellation (dotted white circles) and unique local vasculature (red arrowheads), used to build confidence in the manually assigned 2P-EM cell match (central white circle). All MICrONS data are from a single animal. Scale bars = 5 µm. **b**, Deconvolved calcium traces from 100 imaged neurons. Alternating blue/white column overlay represents the duration of serial video trials, with sample frames of natural videos depicted below. Parametric stimuli (not pictured) were also shown for a shorter duration than natural videos. **c**, Schematic of the digital twin deep recurrent architecture. During training, movie frames (left) are input into a shared convolutional deep recurrent core (orange and blue layers, CVT=convolutional vision transformer, LSTM=long short-term memory) resulting in a learned representation of local spatiotemporal stimulus features. Each neuron is associated with a location (*spatial component* in the visual field (gray layer) to read out feature activations (shaded blue vectors), and the dot product with the neuron-specific learned feature weights (shaded lines, *feature component*) results in the predicted mean neural activation for that time point. **d**, Depiction of 148 manually proofread mesh reconstructions (gray), including representative samples from Layer 2/3 (red), Layer 4 (blue), Layer 5 (green), and Layer 6 (gold). Bottom panel: presynaptic soma locations relative to visual area boundaries.

After functional imaging, the tissue was processed for electron microscopy and imaged (Yin et al., 2020) at 4 × 4 × 40 nm^3^ resolution (Fig. 1a). The EM images were aligned (Mitchell et al., 2019) and automatically segmented using 3D convolutional networks into “atomic” supervoxels, which were agglomerated to create objects (e.g. neurons) with corresponding 3D meshes (Lee et al., 2017; Dorkenwald et al., 2022b; Lu et al., 2021; Wu et al., 2021; Dorkenwald et al., 2022a), and synapses were automatically detected and assigned to presynaptic and postsynaptic partners (Dorkenwald et al., 2022b; Turner et al., 2020; Wu et al., 2021). The analysis presented here is restricted to the overlap of “sub-volume 65” (MICrONS Consortium et al., 2021) and the two-photon functional volume (Fig. 1a), an approximately 560 × 1100 × 500 µm^3^ volume (*in vivo* dimensions) that has been both densely functionally and structurally characterized. Of 82,247 automatically extracted neuronal nuclei in this subvolume, 45,334 were both classified as excitatory and located within the intersection of the EM reconstructed volume and functional volume.

The two-photon and EM volumes were approximately aligned (Fig. 1a, and 13,952 excitatory neurons were manually matched between the two volumes (Fig. 1a; MICrONS Consortium et al. 2021). Retinotopically-matched regions in V1 and higher visual areas AL and RL (together, HVA) were chosen to increase the likelihood of inter-area connections, and visually-responsive neurons within these regions were chosen for manual proofreading. Proofreading focused on extending axonal branches — with an emphasis on enriching projections across the V1/HVA boundary — and on removing false merges (instances where other somas, glia, axons, or dendrites were incorrectly merged into a neuron’s reconstruction) (MICrONS Consortium et al. 2021, Supplemental Table 1). Postsynaptic partners of the proofread neurons were automatically cleaned of false merges with NEURD (Celii et al., 2024). In total, this resulted in a connectivity graph consisting of 148 functionally-characterized presynaptic neurons and 4811 functionally-characterized postsynaptic partners (Fig. 1d).

### Multi-tiered anatomical controls

Connectivity between neurons may be affected by numerous mechanisms, ranging from developmental processes that broadly organize neural circuits, to fine-scale plasticity mechanisms that modulate the strength of individual synaptic connections. The MICrONS volume offers the opportunity to examine function-structure relationships at both of these scales. Because it is densely reconstructed, we not only know the distance between every pair of cell bodies in the volume, but also the relative geometry of their axons and dendrites. With this information, we can determine whether two neurons experience any fine-scale axon-dendrite proximities (ADP), with axon and dendrite coming within 5 µm of each other. Furthermore, for neurons pairs with one or more ADP, we can compute the axon-dendrite co-travel distance *L*_*d*_ (Lee et al., 2016), a pairwise measurement which captures the total extent of postsynaptic dendritic skeleton within 5 µm from any point on the presynaptic axonal skeleton.

With this metric in hand, we can define three cohorts of other neurons for functional comparisons with each presynaptic neuron (Fig. 2a-c, Supplemental Fig. 1). The first cohort are the connected postsynaptic targets of the presynaptic cell; these are neurons in the cortical region of interest that receive at least one synaptic input from the presynaptic neuron. The second group are “ADP controls”, these are neurons with dendrites that come within striking range (5 µm) of the presynaptic axon, but which don’t actually form a synaptic connection. Finally, there are “same region controls” which are non-ADP neurons in the same cortical region (V1 or higher visual area). All connected neurons, ADP controls, and same region controls are restricted to visually responsive neurons with high-quality predictions from the digital twin (see Methods).

**Figure 2.**
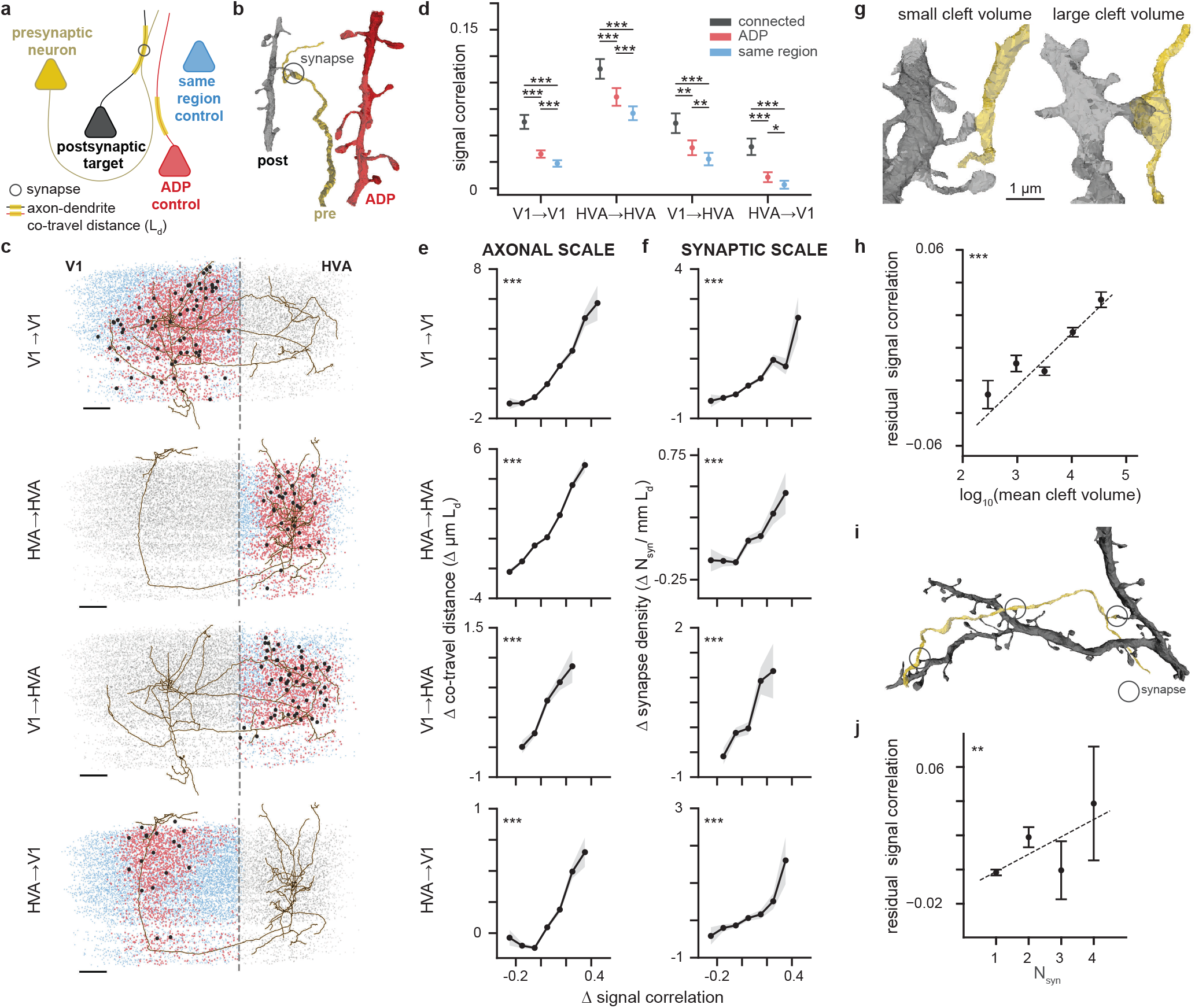
Neurons with higher signal correlation are more likely to form synapses. **a**, Schematic illustrating inclusion criteria for anatomical controls. For each proofread presynaptic neuron (yellow), control neurons for its true postsynaptic partners (black) are drawn either from unconnected neurons with non-zero axon-dendrite co-travel distance (Axonal-Dendritic Proximity (“ADP”), red), or unconnected neurons with zero axon-dendrite co-travel distance located in the same cortical region (blue). The axon-dendrite co-travel distance (*L*_*d*_, yellow highlight on dendrites) is quantified as the total skeletal length of dendrite within 5 *µm* from any point on the presynaptic axon. A synapse is indicated with a gray circle. **b**, Representative meshes demonstrating a true presynaptic (“pre”, yellow axon) to postsynaptic (“post”, black dendrite) pair and an axonal-dendritic proximity control (“ADP”, red dendrite). **c**, Presynaptic neuron axons plotted in EM cortical space for the four projection types (V1→V1, HVA→HVA, V1→HVA, HVA→V1) along with soma centroids of connected partners (black dots), ADP control neurons (red dots), same area control neurons (blue dots) and all other functionally matched neurons that are not used as controls (gray dots). The same presynaptic neuron is plotted for both the V1→V1 and V1→HVA group, and another neuron is used for both the HVA→HVA and HVA→V1 groups to demonstrate that a single presynaptic neuron can be represented in multiple projection types. Dashed line represents the boundary between V1 and HVA. Scale bar: 100*µm* **d**, Mean signal correlation is different (mean ± sem, paired t-test) between synaptically-connected partners (black), ADP controls (red), and same region controls (blue). This relationship was observed for within-area (V1→V1, HVA→HVA), feedforward (V1→HVA), and feedback (HVA→V1) connectivity. For details, see Supplemental Tab. 2 **e**, Axon-dendrite co-travel distance (*µmL*_*d*_) increases in a graded fashion with signal correlation. Δ*L*_*d*_ and Δ signal correlation are the deviations from the mean for each presynaptic neuron. For reference, the mean *L*_*d*_ for each projection type is: V1→V1, 9.03*µm*; HVA→HVA, 9.83*µm*; V1→HVA, 4.17*µm*; HVA→V1, 1.53*µm*. For details of the analysis, see Supplemental Tab. 3, 5 The shaded regions are bootstrap-based standard deviations. **f** As in **e**, but with synapse density (*N*_*syn*_*/mmL*_*d*_). Synapse density increases in a graded fashion with signal correlation, for within-area (V1→V1, HVA→HVA), feedforward (V1→ HVA), and feedback (HVA→V1) connectivity. For reference, the mean *N*_*syn*_*/mmL*_*d*_ for each projection type is: V1→V1, 1.12 synapses / *mmL*_*d*_; HVA→HVA, 0.83 synapses / *mmL*_*d*_; V1→HVA, 1.55 synapses / *mmL*_*d*_; HVA→V1, 1.26 synapses / *mmL*_*d*_. For details of the analysis, see Supplemental Tab. 4, 6 **g**, Representative meshes demonstrating synapses with small cleft volume (896 voxels, left) and large cleft volume (41716 voxels, right). **h**, Synapse size (*log*_10_ cleft volume in voxels) is positively correlated with signal correlation (p-values are from linear regression, residual signal correlation is obtained after regressing out the baseline effects on signal correlation due to differences in *L*_*d*_). **i**, Representative meshes demonstrating a multisynaptic presynaptic (yellow) to postsynaptic (black) pair. **j**, Signal correlations increase with number of synapses (p-values are from linear regression, residual signal correlation is obtained after regressing out the baseline effects on signal correlation due to differences in *L*_*d*_). (For all panels, * = p-value < 0.05, ** = p-value < 0.01, * * * = p-value < 0.001, multiple comparison correction by BH procedure)

At the “axonal scale”, we can ask how selective are axon trajectories within the volume, and whether neurons with axons and dendrites that meet and co-travel together have more similar tuning than nearby neurons that do not have any examples of axon-dendrite proximities. Selectivity at this scale could occur, for example, if a target cortical area has topographically-organized functional properties such as receptive field location (i.e. retinotopy) (Wang and Burkhalter, 2007; Garrett et al., 2014) or preferred orientation (Fahey et al., 2019; Ringach et al., 2016), and if axons preferentially target subregions with similar functional properties. In this case, we would expect functional properties between a presynaptic neuron and its ADP cohort to be more similar than random neurons selected from anywhere within the target region (same region control).

At the “synaptic scale”, we can ask whether there is a relationship between functional properties and connectivity beyond the axonal scale — i.e. beyond what can be explained by the axonal trajectory and the spatial organization of functional properties within the volume. For this analysis, we compare the functional similarity between synaptically-connected neurons on the one hand, and unconnected ADP controls on the other, asking how frequently a certain amount of axon-dendrite co-travel distance is converted to a synapse. One hypothesis is that converting proximities to synapses is independent of the functional similarity between pre- and postsynaptic neurons. In this case, axon trajectories and axon-dendrite proximities would be sufficient to explain all of the observed connectivity between neurons (“Peter’s rule”) (Peters and Feldman, 1976; Braitenberg and Schüz, 2013; Rees et al., 2017). A competing hypothesis is that synapse formation and/or stabilization depends on the functional similarity between pre- and postsynaptic neurons. In this case, we might expect to find an additional boost in synaptic connections in similarly-tuned neurons *above and beyond* whatever selectivity already exists due to axonal trajectories and functional inhomogeneities in the volume. The densely-reconstructed MICrONS volume offers the first opportunity to distinguish between these two hypotheses at a scale spanning layers and areas.

### Functional similarity is enhanced at both the axonal and synaptic scale

We tested the hypothesis of like-to-like connectivity in the context of signal correlations, a more general measure of functional similarity and a better predictor of connectivity in V1 L2/3 than orientation or direction tuning (Cossell et al., 2015). The digital twin was used to calculate the *in silico* signal correlation across a large battery of novel natural movies (250 ten-second clips). This approach was validated in a set of control experiments in a separate cohort of mice to ensure that the *in silico* signal correlation faithfully reproduced *in vivo* signal correlation measurements. In these control experiments, *in silico* signal correlations from the digital twin closely resembled the benchmark *in vivo* signal correlation matrix computed across a set of 30 movie clips each presented ten times, and in fact were more accurate than the *in vivo* signal correlation matrix computed with only six movie clips each presented ten times (which is the number of clips available in the MICrONS data, Supplemental Fig. 2). This excellent correspondence between *in vivo* and *in silico* signal correlation estimates was achieved even though none of the *in vivo* clips were used during training or testing of the digital twin.

For each proofread presynaptic neuron, we computed the mean signal correlation with postsynaptic neurons, ADP controls, and same region controls (Fig. 2d). We found that mean signal correlations were higher for connected neurons than both ADP and same region control groups, indicating that functional properties and connectivity are indeed related at the scale of individual synapses. Furthermore, signal correlations across pairs of neurons that experience at least one axon-dendrite proximity (ADP controls) were significantly higher than same region controls, indicating that there is also functional specificity at the axonal scale, with axons more likely to travel near dendrites of similarly-tuned neurons. These effects were independently observed when subsets of neuron pairs were considered within and across local V1 (V1 → V1), local HVA (HVA → HVA), feedforward (V1 → HVA) and feedback (HVA → V1) projection types (Fig. 2d, see Supp. Tab. 2 for details). In summary, we observed a functional “like-to-like” rule both at the level of axonal trajectories and for connectivity at the synaptic scale.

We explored this finding further, by asking whether there is a graded relationship between the amount of axon-dendrite co-travel distance and the corresponding boost in signal correlations (Fig. 2e).

For this analysis, to avoid confounding variability due to the size of each presynaptic neuron’s axonal arbor and their varying mean signal correlations, for each presynaptic neuron we first computed the mean *L*_*d*_ and mean signal correlations across all of its ADP targets and same region control neurons. Then for each of the pairwise comparisons, we subtracted the pre-computed mean and kept only the difference from the mean for each metric. This approach has the effect of centering both the x- and y-axes in Fig. 2e (and also 2f), in order to focus on the relative effect within each presynaptic neuron and its downstream partners, removing neuron-to-neuron variability in both metrics.

Binning these differences revealed that longer-than-average *L*_*d*_ between a presynaptic neuron and a downstream target was associated with higher-than-average signal correlation between the two neurons. This result was significant when repeated across all projection types, and indicates that the axons and dendrites of neurons with more similar functional properties are likely to meet more often and/or travel farther together in the volume, and there is a graded relationship in this effect that is observed both within and across cortical areas.

We next performed a similar analysis for synapses, looking at connected neuron pairs. For each presynaptic neuron we first computed the mean number of synapses per millimeter of co-travel distance (synapse density, *N*_*syn*_*/mmL*_*d*_), along with mean signal correlations across all pairs of synaptic and ADP targets. Then for each of these pairwise comparisons for a single presynaptic neuron, we subtracted the mean and kept only the difference from the mean. After centering on the means for each presynaptic cell in this way, the binned differences again revealed a strong graded relationship between synaptic connectivity and functional similarity (Fig. 2f). Specifically, higher-than-average rates of synaptic density (synapses per unit co-travel length) were associated with higher-than-average functional similarity, again in a graded fashion.

Given this relationship between synapse frequency and functional similarity, we wondered whether there might be a relationship between functional similarity and either synapse size (a proxy for synaptic strength; (Holler et al., 2021)) and/or the multiplicity of synaptic connections between two neurons. Indeed, previous studies have found that functionally-similar presynaptic-postsynaptic pairs have stronger synaptic connections (Cossell et al., 2015) and larger postsynaptic densities (PSDs) (Lee et al., 2016). In the MICrONS dataset, segmented synapses were automatically annotated with the cleft volume, which is positively correlated to spine head volume, PSD area, and synaptic strength (Celii et al., 2024; Holler et al., 2021; Dorkenwald et al., 2022b) (Fig. 2g).

We found that signal correlation positively correlates with cleft volume (Fig. 2h; r = 0.032, p < 0.001). Looking at the multiplicity of connections between neurons (the number of individual synapses connecting two cells), we also found that presynaptic-postsynaptic pairs with multiple synapses also had higher signal correlations (Fig. 2i, j) when compared to monosynaptic pairs. In both Fig. 2h, j, the synaptic scale effect is isolated by regressing out the contribution of *L*_*d*_ to signal correlation. In summary, both the strength (synaptic volume) and multiplicity of connections are higher when neurons are more functionally similar, consistent with an underlying Hebbian plasticity mechanism that might act to strengthen and stabilize connections between jointly-active neurons.

Lastly, to ensure the robustness of these findings, we ran the same analyses above with signal correlations measured directly from *in vivo* responses (rather than from the digital twin) and found that they replicated the like-to-like results achieved using the *in silico* signal correlations — including the graded relationships at the axonal and synaptic scale, and the relationships with synaptic cleft volume and synapse multiplicity (Supplemental Fig. 3).

**Figure 3.**
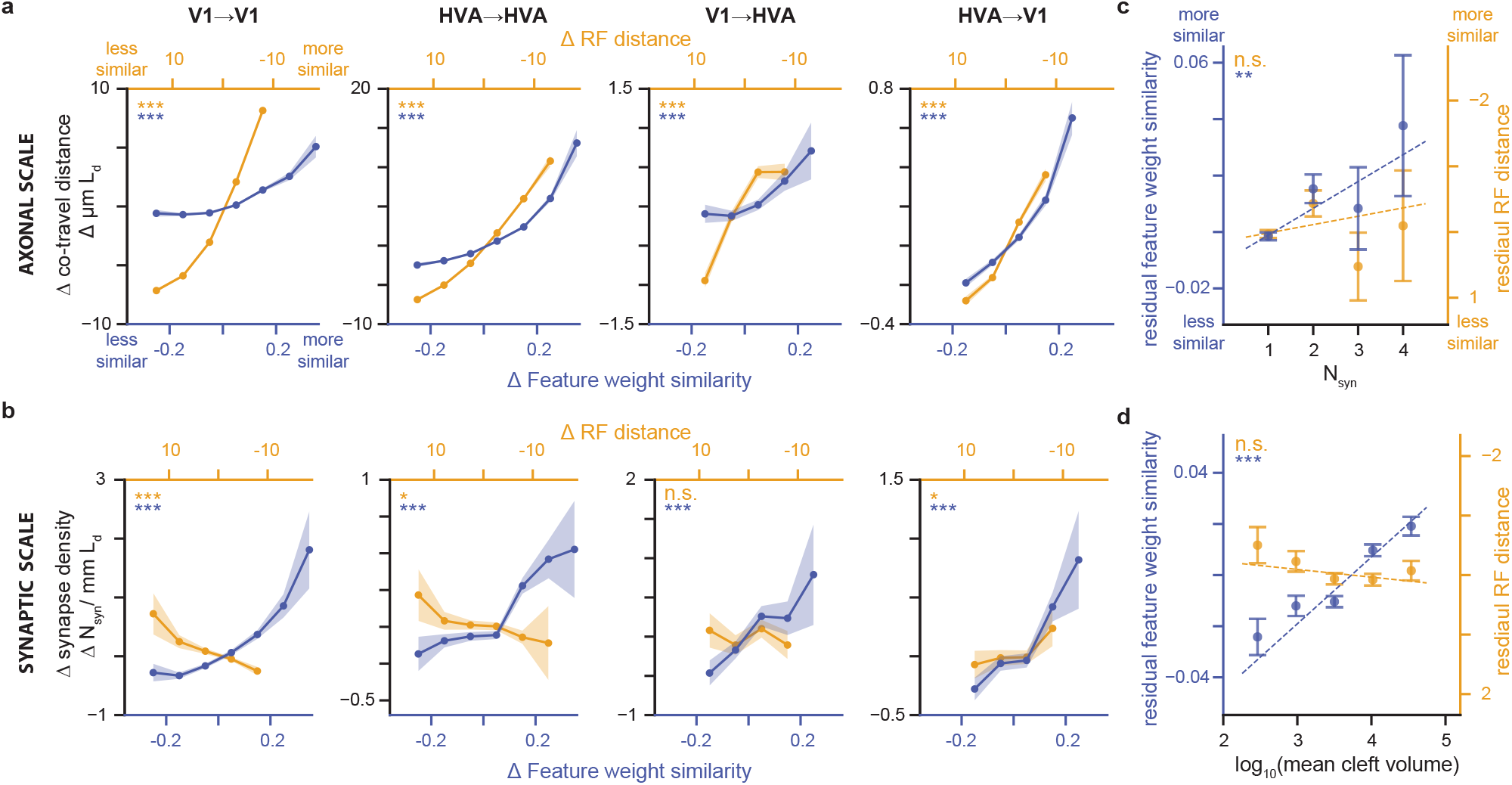
Feature weight similarity predicts synaptic selectivity better than receptive field center distance. **a**, Axon-dendrite co-travel distance increases with feature weight similarity and decreasing RF center distance for within-area (V1→V1, HVA→HVA), feedforward (V1→ HVA), and feedback (HVA→V1) connectivity. **b**,, Synapse density increases with feature weight similarity, but not with RF distance, except for HVA*to*V1 projections. **c**, Multiple synapses are associated with increasing feature similarity, but not receptive field center distance, after regressing out *L*_*d*_. **d**,, Only feature similarity (not receptive field center distance) is associated with an increase in cleft volume, after regressing out *L*_*d*_. (For all panels, * = p-value < 0.05, ** = p-value < 0.01, * * * = p-value < 0.001, p-values are corrected for multiple comparisons using BH procedure, for details, see Supplemental Tab. 11, 13, 15, 17, 12, 14, 16, 18,)

### Factorized *in silico* functional representation

A key advantage of the digital twin (Fig.1c, Wang et al. 2024) is the factorization of each modeled neuron’s predicted response into two factors: readout **location** in visual space—a pair of azimuth/altitude coordinates; and readout **feature weights**— the relative contribution of the core’s learned features in predicting the target neuron’s activity. Intuitively, these learned features can be thought of as the basis set of stimulus features that the network then weighs to predict the neural responses. For each neuron, the combination of feature weights (“what”) and receptive field location (“where”) together encode everything the model has learned about that neuron’s functional properties, and enable the model’s predictive capacity for that neuron. This factorized representation allowed us to examine the extent to which these two aspects of neural selectivity independently contribute to the relationship between signal correlation and connectivity we observed in Fig. 2. Feature weight similarity was measured as the cosine similarity between the vectors of presynaptic and postsynaptic feature weights. Receptive field (RF) location similarity was measured as the visual angle difference between the center of the model readout locations, with lesser distance between the centers (“center distance”) corresponding to greater location similarity. We conducted a separate series of experiments to validate the model’s readout location as an estimate of RF center. These experiments demonstrated that the read-out location correlates strongly with receptive field centers measured using classical sparse noise (dot-mapping) stimuli (Supplemental Fig.4a, b). Moreover, our approach outperformed classical linear *in vivo* measurements of the spatial receptive field for the significant fraction of neurons that are not responsive to the dot-mapping stimuli, even with one hour of dot-mapping data (Supplemental Fig.4c).

### Both RF location similarity and feature weight similarity increase with axon-dendrite co-travel distance

Among pairs of neurons with at least one axon-dendrite proximity (ADP neurons), axon/dendrite co-traveling for longer-than-average distances was associated with higher-than-average feature similarity (Fig. 3a). Similarly, neurons with higher-than-average receptive field similarity (i.e. receptive fields closer to each other), also co-traveled for longer-than-average distances. Thus, both feature tuning and receptive field location are positively correlated with the extent of axon-dendrite proximity between pairs of neurons, and these relationships held both within and across cortical areas. This result is consistent with a scenario where axonal projections are enriched in downstream regions with similar tuning properties, either via axon guidance cues during development or via selective stabilization of axons in areas with similar functional properties, or both.

### A like-to-like rule for feature similarity, but not spatial RF location, is observed at the scale of individual synapses

In contrast with the functional similarity in both features and RF locations associated with axon-dendrite proximity, synaptic connectivity between neurons was only positively correlated with similarity in feature preferences (Fig. 3b). Receptive field location similarity was either not correlated with synapse density or — in the case of V1 — was anti-correlated. Thus, at the synaptic scale, only like-to-like feature preference (not smaller spatial RF center distance) is associated with increased synaptic connectivity. This is a prominent difference between axonal-scale and synaptic-scale relationships with function, and suggests that Hebbian plasticity mechanisms operating at the level of individual synapses are driven by feature similarity rather than receptive field center distance. Consistent with this view, both synapse multiplicity (Fig. 3c) and synaptic cleft volume (Fig. 3d) strongly increase with feature similarity rather than RF location similarity (after regressing out *L*_*d*_ as for Fig. 2h, j).

### Like-to-like rule generalizes across joint layer and area membership of cells

To achieve a more detailed understanding of the organization of connections across layers and areas, for each functional similarity metric (signal correlation, feature weight similarity, and receptive field center distance), we also tested the relationship with connectivity across two areas (primary visual cortex, V1; higher visual areas AL and RL, HVA) and three layers (L2/3, L4, and L5, Fig. 4). For signal correlation (Fig. 4a, b, see Supplemental Tab. 25, 26 for details) and feature weight similarity (Fig. 4c, d, see Supplemental Tab. 27, 28 for details), like-to-like effects (red squares) were widespread across many area and layer combinations, at both the axonal and synaptic scale.

**Figure 4.**
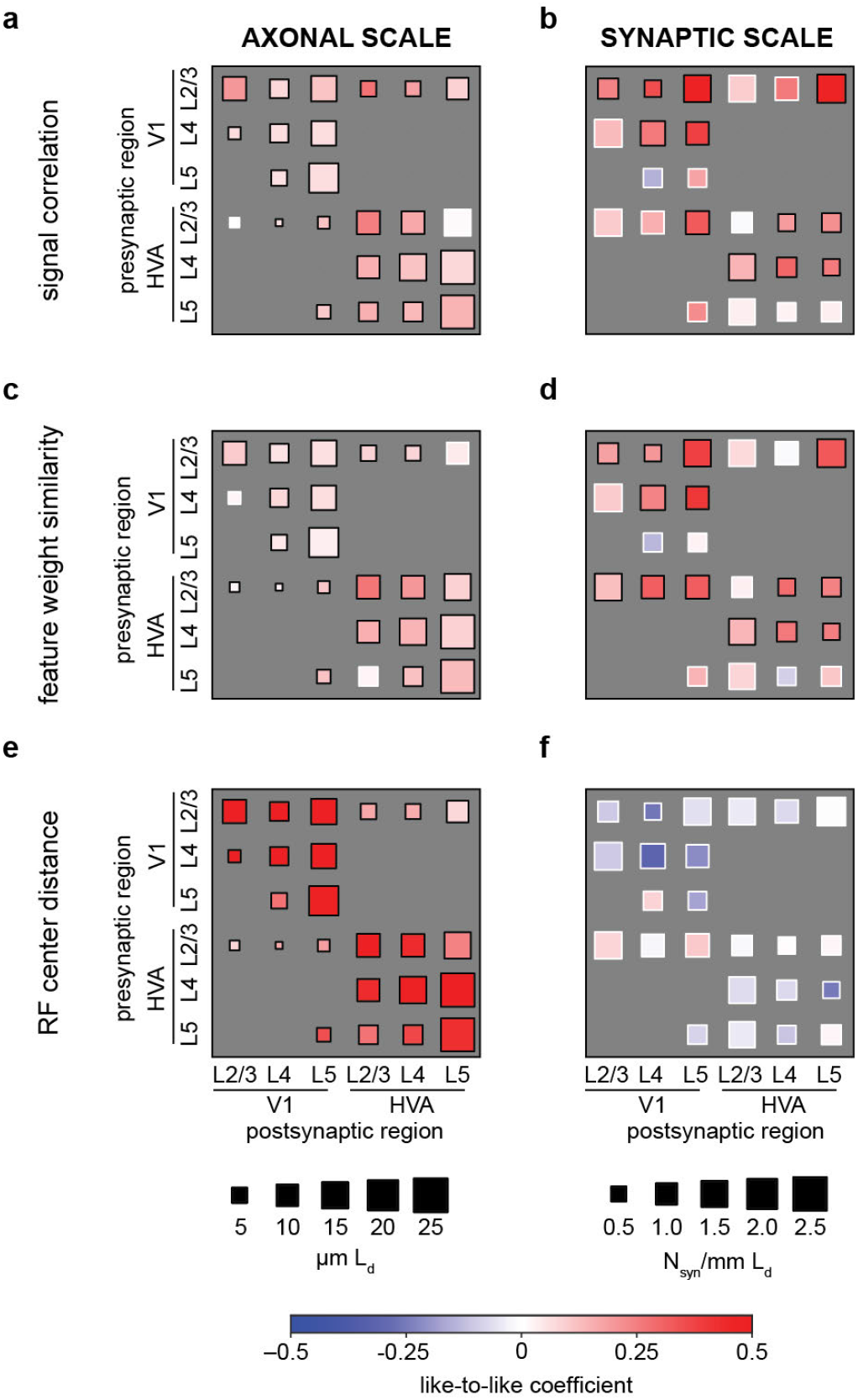
Like-to-like effects are widespread but vary across brain areas, cortical layers, and tuning similarity metrics. **a-f**, Degree of like-to-like broken down by area and layer membership measured at axonal (**a, c, e**) and synaptic scales (**b, d, f**). Colorbar: like-to-like coefficients, red is more like-to-like. For axonal scale, box size represents axon-dendrite co-travel distance (*µmL*_*d*_). For synaptic scale, box size represents synapse density (*N*_*syn*_*/mmL*_*d*_). Like-to-like coefficients are the coefficients of GLMMs fitted to predict axon-dendrite co-travel distance or synapse density with the corresponding functional similarity. (black border = significant at p-value < 0.05, white border = p-value > 0.05, by Wald test after BH correction for multiple comparisons, for details see Supplemental Tab. 26, 25, 28, 27, 30, 29).

In the case of RF center distance, while like-to-like effects (red squares) were widespread at the axonal scale, these effects disappeared when considering synaptic-scale specificity. This finding is consistent with the view that selectivity for retinotopic overlap exists at the scale of axon trajectories but not at the scale of individual synapse formation (Fig. 4e, f, see Supplemental Tab. 29, 30 for details). In this analysis, individual presynaptic baselines (e.g. variable *L*_*d*_, synapse rate, signal correlation), were accounted for with a generalized linear mixed model (GLMM) (see Methods for details). Distributions of all pairwise functional measurements, including *in vivo* signal correlation, *in silico* signal correlation, feature weight similarity, and receptive field distance are provided in Supplementary Fig. 10. Varying the inclusion thresholds of the above analyses across varying levels of digital twin model performance (quartiles of neurons ranked by prediction accuracy) did not substantially change the main results (Supplementary Fig. 11).

### Orientation tuning is like-to-like within V1 at both axonal and synaptic scales

Many neurons in mouse primary visual cortex and higher visual areas are strongly tuned for orientation, and a number of previous functional connectivity studies have used differences in preferred orientation as a metric for visual similarity within V1. In order to compare our findings more directly with this previous work, we repeated the central analysis in Fig. 2, but now using only the difference in preferred orientation — rather than signal correlations — to determine functional similarity.

We used the digital twin to estimate orientation tuning, and we validated this approach with *in vivo* validation experiments (Supplemental Fig. 7a, b), where we compared the *in silico* orientation tuning curve with the tuning curve estimated from the *in vivo* data. Orientation-selective responses were driven by lowpass filtered noise with coherent orientation and motion, a stimulus we have previously used to drive strong visual responses in orientation-tuned cells (Fahey et al., 2019; Wang et al., 2024). For orientation-tuned neurons (gOSI > 0.25, corresponding to more that 50% of co-registered neurons; please see methods for gOSI versus OSI comparison), the *in silico* orientation tuning curves align extraordinarily well with *in vivo* orientation tuning curves (Supplemental Fig. 7c-f).

We found that connected neurons in V1 have more similar orientation tuning than unconnected controls (Supplemental Fig. 8), as reported by previous studies (Rossi et al., 2020; Ko et al., 2011; Lee et al., 2016). However, in contrast with previous studies, we did not observe a similar significant like-to-like effect when restricting the analysis specifically to projections within V1 L2/3 excitatory neurons. To understand this deviation from previous literature, we first determined that connected neuron pairs within V1 L2/3 projections in the MICrONS dataset did indeed have similar orientation preferences (Supplemental Fig. 9), as expected. However, unconnected pairs showed the same level of similarity in orientation preference. We believe this is the result of a local orientation bias where the MICrONS volume is located in V1 (Fahey et al. 2019).

Overall, we found that the model feature weight similarity is a better predictor of connectivity than classical orientation preference, even for neurons tuned to oriented stimuli (Supplemental Fig. 6). Recent work by our group and others has emphasized that optimal stimuli for neurons in mouse V1 can exhibit complex spatial features that deviate strikingly from Gabor-like stimuli (Walker et al., 2019; Tong et al., 2023). These results highlight the advantages of studying more complete tuning functions, such as the model feature weights that we focus on here, rather than single tuning parameters such as orientation preference.

### Neurons with common input are functionally similar

If the pairwise “like-to-like” rule were the sole organizing principle of the visual cortex — implying that all postsynaptic neurons closely resemble their presynaptic partners — we would expect postsynaptic neurons to exhibit a certain degree of similarity to one another.

However, neural feature selectivity likely arises from more complex connectivity rules, so a cohort of neurons down-stream of a single presynaptic neuron might, on average, be less (Fig. 5a, left) or more functionally similar to each other (Fig. 5a, right). To evaluate whether the similarity among postsynaptic neurons differs from what the “like-to-like” rule predicts, we built a simple model network, and introduced the empirical relationships between presynaptic/postsynaptic functional similarity and connectivity that we observed in our data. Specifically, we replicated the empirical distribution of signal correlations, feature weight similarities, and receptive field location distances over all model neuron pairs, and then predicted the expected number of synapses between neuron pairs — based on their functional similarity — with a Poisson linear mixed-effects model (Fig. 5b). We confirmed that this model replicated the expected functional similarity between connected neurons, indicating that it accurately captured the same pairwise “like-to-like” rule that we observed in the data (Supplemental Fig. 5). Then, we measured the similarity among all postsynaptic neurons downstream of a single presynaptic neuron, by calculating the mean pairwise signal correlations. As expected, on average, postsynaptic neurons were more functionally-similar to other postsynaptic neurons than random pairs (Supplemental Fig. 5). However, we also found that postsynaptic neurons receiving common synaptic inputs in the MICrONS dataset were even more similar than the “like-to-like” model predicted (Fig. 5c). These relationships held when tested at both axonal and synaptic scales for three out of the four projection types (Supplemental Fig. 5). This suggests the existence of higher-order functional organization beyond the simple pairwise relationships that we focused on up to this point.

**Figure 5.**
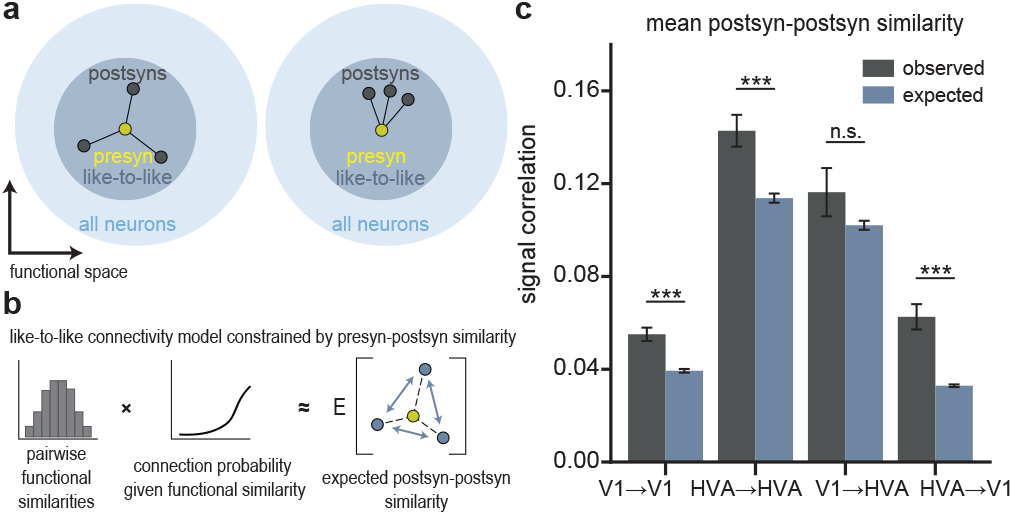
Postsynaptic neurons with a common input are more functionally similar to each other than expected from a pairwise like-to-like rule. **a**, Left: Schematic illustrating the null hypothesis that postsynaptic neurons (gray circles, “postsyns”) of a common presynaptic neuron (yellow circle, “presyn”) have no additional feature similarity with each other beyond their like-to-like similarity with their common presyn. In this scenario, postsyns are distributed uniformly around the presyn in the “like-to-like” region of functional space (dark blue region). Right: Schematic illustrating the alternative hypothesis that the postsynaptic neurons are closer in functional space than predicted from a pairwise like-to-like rule, equivalent to being clustered non-uniformly within the “like-to-like” region. **b**, Schematic illustrating the functional connectivity model used to simulate the null hypothesis in **a**. Pairwise functional measurements (left) — including signal correlations, feature weight similarity and receptive field location distance — were passed through a function relating functional similarity to connection probability. Then, within this modeled network, we computed the pairwise similarity of all postsyns downstream of a common presyn (right). In **c**, we compare the actual postsynaptic functional similarity we observed in the data (black) to the expected postsyn similarity as determined from the model (blue). In three out of four area comparisons, we find that postsyns are significantly more similar to each other than expected from a pairwise like-to-like rule.

### Like-to-like connectivity and its functional role in artificial neural networks

A possible functional role for the like-to-like connectivity observed in our data is suggested by theoretical work on recurrent neural network (RNN) models, starting with early work on attractor-based models like Hopfield networks (Hopfield, 1982; Khona and Fiete, 2022a). In these models, like-to-like connectivity increases similarity in neural responses to similar stimuli, which aids both supervised and unsupervised learning. Artificial neural networks produce stimulus representations similar to those in primary visual cortex (Yamins et al., 2014; Cadena et al., 2019), but there have been no comparisons to date between like-to-like connectivity in biological and artificial neural networks.

To make such a comparison, we trained an RNN model on a simple image classification task (Fig. 6a; see Methods for details). The trained RNN showed increased like-to-like connectivity compared to the same model before training (Fig. 6b,c), and a small shift in the distributions of signal correlations, similar to those in our data (Supplementary Figs. 10 and 12). Intriguingly, we found that ablating like-to-like connections in the trained model decreased performance more than ablating random connections with the same connection strengths (Fig. 6d), suggesting that like-to-like connectivity plays a functional role in the model. Finally, we found that the trained model exhibits an increase in signal correlations within cohorts of postsynaptic cells defined by a shared presynaptic neuron, similar to the higher order connectivity rule that we observed in our data. (Fig. 6e). These results suggest that like-to-like connectivity — similar to the magnitude we observed in the MICrONS data — could be sufficient to subserve a functional role in sensory processing and learning.

**Figure 6.**
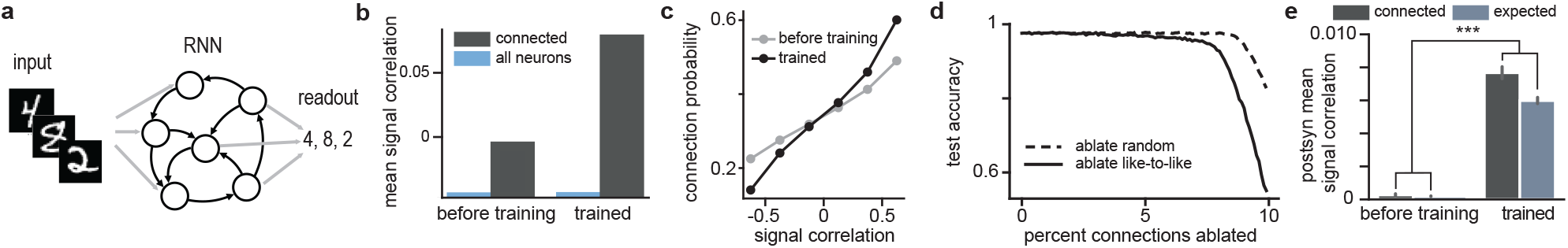
Like-to-like connectivity in an RNN. **a**, A vanilla RNN was provided images as inputs and weights were trained so that a readout of the final state identifies the input’s label. **b**, Mean signal correlations among all (blue) and connected (black) neuron pairs for the same RNN before (left) and after (right) training. Neurons were classified as connected when their weights exceeded a fixed threshold. **c**, Connection probability as a function of signal correlation for the same network before (gray) and after (black) training. **d**, Test accuracy of the network as a function of the number of connections ablated when ablating random (dashed) or like-to-like (solid) connections. Connections were classified as like-to-like whenever the weight and signal correlation both exceeded a fixed threshold. **e**, Mean post-post signal correlations and the expected post-post signal correlation given a pairwise model similar to Fig. 5c before and after training.

## Discussion

Discovering the principles that relate structure to function is central in the pursuit of a circuit-level mechanistic understanding of brain computations. Here, we used the MICrONS multi-area dataset — the largest of its kind — to study the relationship between the connections and functional responses of excitatory neurons in mouse visual cortex across cortical layers and visual areas. Our findings revealed that neurons with highly correlated responses to natural videos (i.e. high signal correlations) tended to be connected with each other, not only within the same cortical areas but also across multiple layers and visual areas, including feedforward and feedback connections. While the overall principle of “like-to-like” connectivity that we describe here is consistent with a number of previous studies (Ko et al., 2011, 2013; Wertz et al., 2015; Lee et al., 2016; Rossi et al., 2020), our work leverages three unique strengths of the MICrONS dataset to extend and refine these previous findings.

First, the **scale of the volume** enabled us to look at connection principles across layers 2-5 of cortex, not just within V1, but also in projections between V1 and higher visual areas. In agreement with previous findings from V1 L2/3, we found that pairs of cells with higher signal correlations were more likely to be connected (Ko et al., 2011, 2013; Cossell et al., 2015). This general principle held not just in V1 L2/3, but also in higher visual areas and for inter-area feedforward and feedback projections.

Second, we were able to take advantage of the **dense reconstruction** to ask questions about functional specificity at the axonal scale that would be difficult to address with any other data. We found that axons are more likely to co-travel with dendrites of similarly-tuned neurons, even for long-range axons spanning areas. The dense reconstruction also allowed us to compute a set of null distributions for the expected synaptic connectivity between neurons based on axon-dendrite proximities. These controls enable us to distinguish whether the relationships we observed between connectivity and function are due to the overall geometry of axonal and dendritic arbors in the volume, or whether they reflect a more precise connectivity rule at the level of individual synapses. For example, it is only with the inclusion of both same region and ADP controls that we are able to observe the diverging findings of axon trajectory level selectivity for receptive field center distance (Fig. 3 d, e, f) and synaptic level selectivity for feature weight similarity (Fig. 3 a, b, c). These different controls can be mapped onto potential developmental or adult plasticity mechanisms that may shape the coarse axon trajectory and fine-scale synaptic connectivity across the brain.

Finally, our **deep learning neural predictive modeling approach** enabled us to comprehensively characterize the tuning function of a neuron, factorize it into spatial and feature tuning components, and facilitate *in silico* exploration with neural responses to novel visual stimuli. The digital twin model allowed us to measure signal correlations over a much larger set of naturalistic videos, resulting in better connectivity predictions compared to *in vivo* measurements from a smaller stimulus set (Supplemental Fig. 6). Moreover, the model’s factorized architecture provided a unique opportunity to discover distinct synaptic organizing principles for two interpretable components of neuronal tuning: what the neuron is tuned to and where its receptive fields are located. Notably, the digital twin model demonstrated excellent out-of-training-set performance (Supplemental Fig. 2) even for novel stimulus domains (Supplemental Fig. 4). This generalization ability opens exciting possibilities for future *in silico* visual experiments, although validation experiments remain essential when studying the digital twin model with new stimulus domains. Currently we treat this model as a black box, but future models could constrain the architecture in order to make internal model parameters more interpretable. Additionally, recent studies have shown that behavioral states and task variables explain a substantial portion of neural responses, even in sensory cortices (Stringer et al., 2019; Musall et al., 2019). Future digital twins could incorporate additional behavioral measurements that enable us to study more general relationships between structure and function, beyond visual processing.

Many of the analyses described in this paper evaluated pair-wise relationships between one presynaptic neuron and one postsynaptic or control neuron. While our experiment in an RNN toy model shows that a pairwise like-to-like rule can have important functional consequences for task performance (Fig. 6), there is still a question of whether there exist higher-order functional motifs beyond simple, pairwise relationships. We explored one such higher-order pattern in our analysis of functional similarity among postsynaptic neurons sharing at least one common input (Fig. 5). This investigation revealed functionally similar postsynaptic cohorts, suggesting the presence of more complex organizational principles. Other studies have looked at functional similarity in presynaptic cells converging on a single common postsynaptic neuron (Bock et al., 2011; Wertz et al., 2015; Rossi et al., 2020). As proofreading in the MICrONS volume continues, it will become possible to test motifs of much higher order and complexity and their relationship to more complex functional properties. In addition to a more complete connectivity graph, another route to discovery may be to study more richly-colored graphs that include additional modalities about each neuron, including features such as morphological or transcriptomic information, or local ultrastructure. Alternatively, it may be important to investigate functional connectivity rules operating at the scale of sub-cellular compartments, for example looking at synapse clustering on dendrites.

Just as pairwise relationships might only partially reveal rules at play in higher-order motifs, the principle of “like-to-like” may only partially capture more complicated principles relating structure and function. For example, Wertz et al. found that for some networks (multiple presynaptic neurons converging onto a single postsynaptic output), the similarity of inputs differed depending on layer origin, a phenomenon they termed “feature-variant” networks (Wertz et al., 2015). Several studies have also found that there is an interplay between the geometric relationship of receptive field positions and feature preferences (Rossi et al., 2020; Marques et al., 2018; Oldenburg et al., 2024). For example, Rossi et al. found that the spatial offset between the receptive fields of excitatory and inhibitory inputs matched the postsynaptic cell’s direction selectivity (Rossi et al., 2020). Future work could further discriminate along the relevant feature dimensions to find more precise rules.

Like-to-like connectivity is a recurring theme in theoretical models of neural circuit function, including Hebb’s theory of neural assemblies (Hebb, 1949), Hubel and Wiesel’s theory of receptive field formation (Hubel and Wiesel, 1962), and later work by Hopfield (Hopfield, 1982) and others (Khona and Fiete, 2022b) on attractor based models. Like-to-like connectivity is often assumed *a priori* or emerges due to Hebbian plasticity in these models, but our analysis of a vanilla RNN trained by gradient descent shows that like-to-like connectivity — in addition to higher order connectivity motifs observed in our data — can arise naturally from optimizing a recurrent system for a simple visual task (Fig. 6). Theory-driven experiments will allow us to move beyond correlational to causal understanding of biological systems. Future works could study these connectivity rules further in artificial systems with greater complexity or biological realism.

Our work provides a first glimpse of principles of cortical organization that can be discovered with large datasets combining detailed functional characterization with synaptic-scale connectivity. While the incredible accuracy of machine learning-based reconstruction methods has rightly increased optimism about the potential discoveries that can be made from large EM volumes — especially when combined with functional characterization — we should also not forget the magnitude of the challenge contained in even a 1 mm^3^ volume of cortex in a single mouse. The analyses in this paper are based on only a small number of manually proofread neurons, but even this limited view of the dataset represents an impressive volume of axonal and dendritic reconstruction. Ongoing investments in proofreading, matching, and extension efforts within this volume will have exponential returns for future analyses as they yield a more complete functional connectomic graph, and reduce or eliminate potential biases in the connections. As more large-scale datasets like MICrONS are publicly released, there will be much more to discover about the organizing principles relating structure and function in other brain areas (Kuan et al., 2024) and even other model organisms (Wanner et al., 2016). Our hope is that this dataset, including both the structural anatomy and the immortalized digital twin for ongoing *in silico* experiments, will be a community resource that will yield concrete insights as well as inspiration about the scale of investigation that is now possible in neuroscience.

## ACKNOWLEDGEMENTS

The authors thank David Markowitz, the IARPA MICrONS Program Manager, who coordinated this work during all three phases of the MICrONS program. We thank IARPA program managers Jacob Vogelstein and David Markowitz for co-developing the MICrONS program. We thank Jennifer Wang, IARPA SETA for her assistance. The work was supported by the Intelligence Advanced Research Projects Activity (IARPA) via Department of Interior/ Interior Business Center (DoI/IBC) contract numbers D16PC00003, D16PC00004, and D16PC0005. The U.S. Government is authorized to reproduce and distribute reprints for Governmental purposes notwithstanding any copyright annotation thereon. XP acknowledges support from NSF CAREER grant IOS-1552868. ZD, SP, RR, KJ, XP, and AST acknowledge support from NSF NeuroNex grant 1707400. AST, XP, KJ, and JR are supported by RF1 MH130416. AST acknowledges support from National Institute of Mental Health and National Institute of Neurological Disorders And Stroke under Award Number U19MH114830 and National Eye Institute award numbers R01 EY026927 and Core Grant for Vision Research T32-EY-002520-37. ABK acknowledges support from a training fellowship from the Gulf Coast Consortia, on the NLM Training Program in Biomedical Informatics & Data Science (T15LM007093). RR acknowledges support from Air Force Office of Scientific Research (AFOSR) award number FA9550-21-1-0223. Disclaimer: The views and conclusions contained herein are those of the authors and should not be interpreted as necessarily representing the official policies or endorsements, either expressed or implied, of IARPA, DoI/IBC, or the U.S. Government.

## AUTHOR CONTRIBUTIONS

We adopted the following contribution categories from CRediT (Contributor Roles Taxonomy). Authors within each category are sorted in the same order as in the author list.

**Conceptualization**: ZhuD, PGF, StP, XP, JR, AST

**Methodology**: ZhuD, PGF, StP, ErYW, BC, CP, AC, DY, EF, KJ, RR, EdYW, AST

**Software**: ZhuD, PGF, StP, ErYW, BC, CP, DY, EF, RR

**Validation**: ZhuD, PGF, StP, ErYW

**Formal analysis**: ZhuD, PGF, StP, AC, ABK, RR

**Investigation**: ZhuD, PGF, StP, ErYW, SaP, JF, ZhiD, KP, TM, RR

**Resources**: PGF, StP, SaP, JAB, ALB, DB, JB, DJB, MAC, EC, SD, LE, AH, ZJ, CJ, DK, NK, SK, KiL, KaL, RL, TM, GM, EM, SSM, SM, BN, SeP, CMSM, WS, MT, RT, NLT, WW, JW, WY, SY, DY, EF, FS, RR, HSS, FC, NMdC, RCR, XP, JR, AST

**Data Curation**: ZhuD, PGF, StP, BC, CP, ZhiD

**Writing - Original Draft**: ZhuD, PGF, StP, ErYW, RR, JR

**Writing - Review & Editing**: ZhuD, PGF, StP, AC, JF, SaP, CMSM, KJ, RR, FC, NMdC, RCR, EdYW, XP, JR, AST

**Visualization**: ZhuD, PGF, StP, RR

**Supervision**: PGF, XP, JR, AST

**Project administration**: ZhuD, PGF, StP, JR, AST

**Funding acquisition**: HSS, FC, NMdC, RCR, XP, JR, AST

## Methods

### MICrONS Dataset

MICrONS dataset was collected in a single animal as described in MICrONS Consortium et al. (2021), including neurophysiological data collection, visual stimulation, stimulus composition, EM data collection, automatic EM segmentation and reconstruction, manual EM proofreading, volume coregistration, and manual soma-soma matching between the functional and EM volumes. Neuro-physiological experiments, Visual Stimulution,and Stimulus Composition sections below are specific to additional experiments described in Supplemental Fig. 7.

### Neurophysiological experiments

All procedures were approved by the Institutional Animal Care and Use Committee of Baylor College of Medicine. Ten mice (Mus musculus, 3 female, 7 males, 78-190 days old at first experimental scan) expressing GCaMP6s in excitatory neurons via Slc17a7-Cre and Ai162 transgenic lines (recommended and generously shared by Hongkui Zeng at Allen Institute for Brain Science; JAX stock 023527 and 031562, respectively) were anesthetized and a 4 mm craniotomy was made over the visual cortex of the right hemisphere as described previously (Reimer et al., 2014; Froudarakis et al., 2014).

Mice were head-mounted above a cylindrical treadmill and calcium imaging was performed with a mesoscope (Sofroniew et al., 2016) as described in release (MICrONS Consortium et al., 2021), with surface power not exceeding 20 mW, depth constant of 220 µm, and greatest laser power of ∼ 86 mW was used at approximately 400 µm from the surface.

The cranial window was leveled with regard to the objective with six degrees of freedom. Pixel-wise responses from an ROI spanning the cortical window (3600 × 4000 µm, 0.2 px/µm, 200 µm from surface, 2.5 Hz) to drifting bar stimuli were used to generate a sign map for delineating visual areas (Garrett et al., 2014).

For the orientation tuning validation data in Supplemental Fig. 7, our target imaging site was a 1200 × 1100 µm^2^ area spanning L2-L5 at the conjunction of lateral primary visual cortex (V1) and three lateral higher visual areas: anterolateral (AL), lateromedial (LM), and rostrolateral (RL). This resulted in an imaging volume that was roughly 50% V1 and 50% higher visual area. This target was chosen in order to mimic the area membership and functional property distribution in the MICrONS animal. Each scan was performed at 6.3 Hz, collecting eight 620 × 1100 µm^2^ fields per frame at 0.4 px*/*µm xy resolution to tile a 1190 − 1200 × 1100 µm^2^ FOV at four depths (two planes per depth, 40 − 50 µm overlap between coplanar fields). The four imaging planes were distributed across layers with at least 50 µm spacing, with two planes in L2/3 (depths: 180 µm, 230 µm), one in L4 (325 µm), and one in L5 (400 µm).

Movie of the animal’s eye and face was captured throughout the experiment. A hot mirror (Thorlabs FM02) positioned between the animal’s left eye and the stimulus monitor was used to reflect an IR image onto a camera (Genie Nano C1920M, Teledyne Dalsa) without obscuring the visual stimulus. The position of the mirror and camera were manually calibrated per session and focused on the pupil. Field of view was manually cropped for each session. The field of view contained the left eye in its entirety, 212-330 pixels height x 262-424 pixels width at 20 Hz. Frame times were time stamped in the behavioral clock for alignment to the stimulus and scan frame times. Video was compressed using Labview’s MJPEG codec with quality constant of 600 and stored the frames in AVI file.

Light diffusing from the laser during scanning through the pupil was used to capture pupil diameter and eye movements. A DeepLabCut model (Mathis et al., 2018) was trained on 17 manually labeled samples from 11 animals to label each frame of the compressed eye video (intraframe only H.264 compression, CRF:17) with 8 eyelid points and 8 pupil points at cardinal and intercardinal positions. Pupil points with likelihood *>*0.9 (all 8 in 69.8-99.2% of frames per scan) were fit with the smallest enclosing circle, and the radius and center of this circle was extracted. Frames with *<* 3 pupil points with likelihood *>*0.9 (*<*1.1% frames per scan), or producing a circle fit with outlier *>* 5.5 standard deviations from the mean in any of the three parameters (center x, center y, radius, *<*0.1% frames per scan) were discarded (total <1.2% frames per scan). Gaps were filled with linear interpolation. The mouse was head-restrained during imaging but could walk on a treadmill. Rostro-caudal treadmill movement was measured using a rotary optical encoder (Accu-Coder 15T-01SF-2000NV1ROC-F03-S1) with a resolution of 8000 pulses per revolution, and was recorded at ∼ 100 Hz in order to extract locomotion velocity.

### Visual stimulation

For the validation data in Supplemental Fig. 2, 4 and 7, monitor size and positioning relative to the mouse were as described in MICrONS Consortium et al. (2021), with the exception of replacing the dot stimulus for monitor positioning with 10 × 10 grid tiling a central square (approx 90 degrees width and height) with 10 repetitions of 200 ms presentation at each location.

A photodiode (TAOS TSL253) was sealed to the top left corner of the monitor, and the voltage was recorded at 10 KHz and timestamped with a 10 MHz behavior clock. Simultaneous measurement with a luminance meter (LS-100 Konica Minolta) perpendicular to and targeting the center of the monitor was used to generate a lookup table for linear interpolation between photodiode voltage and monitor luminance in cd/m^2^ for 16 equidistant values from 0-255, and one baseline value with the monitor unpowered.

At the beginning of each experimental session, we collected photodiode voltage for 52 full-screen pixel values from 0 to 255 for one second trials. The mean photodiode voltage for each trial was collected with an 800 ms boxcar window with 200 ms offset. The voltage was converted to luminance using previously measured relationship between photodiode voltage and luminance and the resulting luminance vs voltage curve was fit with the function *L* = *B* + *A* · *P* ^*γ*^ where L is the measured luminance for pixel value P, and the *γ* of the monitor was fit as 1.73. All stimuli were shown without linearizing the monitor (i.e. with monitor in normal gamma mode).

During the stimulus presentation, display frame sequence information was encoded in a 3 level signal, derived from the photodiode, according to the binary encoding of the display frame (flip) number assigned in-order. This signal underwent a sine convolution, allowing for local peak detection to recover the binary signal together with its behavioral time stamps. The encoded binary signal was reconstructed for *>*93% of the flips. Each flip was time stamped by a stimulus clock (MasterClock PCIe-OSC-HSO-2 card). A linear fit was applied to the flip timestamps in the behavioral and stimulus clocks, and the parameters of that fit were used to align stimulus display frames with scanner and camera frames. The mean photodiode voltage of the sequence encoding signal at pixel values 0 and 255 was used to estimate the luminance range of the monitor during the stimulus, with minimum values of approximately 0.003-0.60 cd/m^2^ and maximum values of approximately 8.68-10.28 cd/m^2^.

### Preprocessing of neural responses and behavioral data

Fluorescence traces from the MICrONS dataset and the additional data for Supplemental Fig. 2, 4 and 7 were detrended, deconvolved, and aligned to stimulus and behavior as described in Wang et al. (2024), and all traces were resampled at 29.967 Hz. Possible redundant traces, where a single neuron produced segmented masks in multiple imaging fields, were all kept for downstream model training. We elected to remove one of the 14 released scans from the analysis (session 7, scan_idx 4) due to compromised optics (water ran out from under the objective for ∼ 20 minutes), leaving 13 scans.

### Model architecture and training of the digital twin model

The model architecture and training for the digital twin model used for assessing in silico signal correlation, feature weight similarity, and receptive field center distance is the same as the cvt-lstm model described in Wang et al. (2024).

Briefly, the core network of the cvt-lstm models was trained on 8 scans collected from 8 mice with natural movie stimuli to capture cortical representations of visual stimuli shared across mice. The parameters of the core network are then frozen, and the rest of the network parameters are trained for each scan with trials where natural movies are shown in the MICrONS dataset. Trials were excluded from model training if more than 25% of their pupil frames were untrackable. This issue most commonly arose when the animal closed its eye, rendering the functional relationship between neural activity and the visible stimulus ambiguous. The number of excluded trials varied across scans, ranging from 2 to 123 per scan, representing 0.6-38.0% of total trials.

To assess orientation tuning similarity, we used a slightly different digital twin model with a conv-lstm architecture as described in Wang et al. (2024). The core network of the convlstm models was trained with the same 8 scans as the cvt-lstm model. The rest of the network parameters are fine-tuned with both natural movies and oriented noise stimuli available from the MICrONS dataset to reach maximum alignment between in vivo and in silico orientation tuning.

### Functional unit inclusion criteria

In order to focus our analyses on neurons that are visually responsive and well modeled by the digital twin, we applied a dual functional threshold over two metrics (*in vivo* reliability and model prediction performance) prior to all analyses related to signal correlation, receptive field center distance, and feature weight similarity.

#### In vivo reliability threshold

In order to estimate the reliability of neuronal responses to visual stimuli, we computed the upper bound of correlation coefficients for each neuron(*CC*_*max*_, Schoppe et al. 2016) across 60 seconds of natural movie stimuli repeated 10 times across the stimulus period (10 min total). *CC*_*max*_ was computed as:

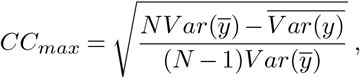

where *y* is the *in vivo* responses, and *N* is the number of trials. A threshold of *CC*_*max*_ > 0.4 was applied. Where more than one two-photon functional unit was matched to a given EM unit, the functional trace with the higher oracle score was used for analysis.

#### Model prediction performance threshold

In order to focus our analyses on neurons for which adequate model performance indicated sufficiently accurate representation of the neuronal tuning features, we computed the test correlation coefficient on the withheld oracle test dataset, which was not part of the training set. Test correlation coefficients (*CC*_*abs*_) were computed as:

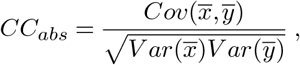

where *x* is the *in silico* response and *y* is the *in vivo* response. A threshold of *CC*_*abs*_ > 0.2 was applied.

144 out of 148 presynaptic neurons and 3920 out of 4811 postsynaptic neurons passed the dual functional unit inclusion criteria.

#### Oracle score

The oracle score was computed for all units as described in MICrONS Consortium et al. (2021). Oracle score is later used to select presynaptic neurons for morphological proofreading (see below).

### Two-photon/ Electron Microscopy Matching

The matching between two-photon functional units and EM cells aligns closely with table `coregistration_manual_v4` (MICrONS Consortium et al., 2021) with some additional restrictions applied. First, the matches to the excluded scan described in Section **Preprocessing of neural responses and behavioral data** were removed. Then, two thresholds were applied directly to the table (residual *<* 20 and score *>* −10).

### Morphological Proofreading

While automation of the EM segmentation has progressed to where dense reconstruction is possible at the millimeter scale, even state-of-the-art methods still leave imperfections in the graph relative to human expert performance. The two categories of reconstruction error are false merges (the incorrect grouping of segmented objects, such as including an axon or dendrite that does not belong to a specific soma) and false splits (the incorrect separation of objects, such as excluding an axon or dendrite that does belong to a specific soma). These errors lead to incorrect associations between pre- and post-synaptic partners and ultimately an incorrect connectivity graph. Proof-reading corrects false merges by “cleaning” the reconstruction, i.e. removing incorrectly associated segments, and corrects false splits by “extending” the reconstruction, i.e. adding back missing segments. We used two proofreading approaches in this study: manual and automatic. “Manual proofreaders” were trained to both clean and extend reconstructions to a high degree of accuracy, as validated by expert neuroanatomists. All of the presynaptic cells in this study were manually proofread. The manual proofreading protocol can be found in the primary dataset paper, (MICrONS Consortium et al., 2021). For the rest of the cells (postsynaptic and control neurons), we used the NEURD package (Celii et al., 2024) to perform automated proofreading. Automated proofreading cleans reconstructions to a high degree of accuracy relative to manual proofreaders, but it does not extend reconstructions.

#### Dendritic Proofreading

At baseline, reconstructed dendrites were generally complete and required little extension (Elabbady et al., 2024). However, they often contained false merges that required cleaning (Elabbady et al., 2024). The dendrites of all of the presynaptic neurons were manually cleaned and extended. The dendrites of other neurons were cleaned with NEURD (Celii et al., 2024).

#### Axonal Proofreading

At baseline, reconstructed axons require both cleaning and extension (Elabbady et al., 2024). Only the axons of presynaptic neurons were manually cleaned and extended. In order to balance morphological completeness (per neuron) and coverage (across projection types), we extended axons to varying degrees of completion. Specifically, we performed full manual proofreading on a subset of neurons (n=84), which involved thoroughly cleaning and extending all axonal branches throughout the dataset. For the remaining neurons (n=64), we applied partial proof-reading, focusing exclusively on extending axonal branches that were pre-screened to feedback from HVA to V1. The full list of proofread presynaptic neurons, their area and layer membership, and whether they were fully or partially proof-read is included in Supplemental Table 1, and a subset of proofread axons are shown in Supplemental Fig. 1.

### Presynaptic Neuron Selection

Our approach for selecting presynaptic neurons for manual proofreading was designed to enrich for higher-order connectivity motifs within and (especially) across visual areas. Because connection probability drops off with distance (Holmgren et al., 2003), we elected to initially focus proofreading efforts on spatially clustered cells in two cylindrical columns spanning cortical layers 2-5, with the first column located in V1 and the second located in RL. Column centers were chosen according to retinotopic maps, as it has been shown that inter-areal projections are retinotopically matched (Wang and Burkhalter, 2007; Marques et al., 2018). During the proofreading process we added an additional column in V1 and another spanning the RL and AL border, to increase coverage of the volume. Lastly, a few HVA cells that were postsynaptic to proofread V1 cells were chosen to enrich for higher order motifs (n=9). All neurons selected for proofreading had an oracle score greater than 0.25 and model test correlation (model predictive performance from an intermediate version of the digital twin) greater than 0.15. The first 40 neurons were selected by experienced neuroscientists unblinded to functional properties for an emphasis on functional diversity. All remaining neurons were chosen blind to functional properties.

### Anatomical controls

In order to control for anatomy at the axonal scale, we recruited all visually responsive, well predicted, functionally matched excitatory neurons (*CC*_*max*_ > 0.4, *CC*_*abs*_ > 0.2) that are located in the same region as the postsynaptic target, but are not observed to form a synapse with the presynaptic neuron (same region control). Area membership labels per neuron were used from the MICrONS release (MICrONS Consortium et al., 2021). Additionally, control candidates that meet criteria for both the same region control and the ADP control (described below) will only be included in ADP control.

In order to control for anatomy at the synaptic scale, we recruited all visually responsive, well predicted, functionally matched excitatory neurons (*CC*_*max*_ > 0.4, *CC*_*abs*_ > 0.2) with a dendritic skeleton passing within 5 µm of the presynaptic neuron axonal synapse in the presynaptic axonal arbor (3D euclidean distance), but which are not observed to form a synapse with the presynaptic neuron (ADP control). Presynaptic axonal skeletons were computed using the `pcg_skel` package developed by collaborators at the Allen Institute for Brain Science (Schneider-Mizell et al., 2024; Schneider-Mizell and Collman, 2023). For postsynaptic dendritic skeletons, we used the automatically proofread and skeletonized dendritic arbors as described in Celii et al. 2024.

To compute the axon-dendrite co-travel distance (*L*_*d*_) between a pair of neurons, we first discretized both the axonal skeleton of one neuron and the dendritic skeleton of the other neuron so that no edge exceeded a length of 1 *µm*. Next, we identified all pairs of vertices from the two skeletons that were within 5 *µm* of each other by performing spatial queries using KDTree’s `query_ball_tree` method from the `scipy.spatial` module in SciPy (Virtanen et al., 2020). From these proximal vertices (“proximity”), we identified the associated dendritic edges. The lengths of these dendritic edges were summed to obtain *L*_*d*_.

Synapses were obtained from Table `synapses_pni_2` (MICrONS Consortium et al., 2021) and were assigned to an axon-dendrite proximity if they were within 3 *µm* of any vertex in the proximity.

In the case of the joint area and layer analysis (Fig. 4), candidates in both the “same region” and “ADP” controls must additionally match the same layer classification as the post-synaptic target in order to be included. Layer assignment was performed as in (Weis et al., 2024).

### Measuring functional similarities

#### In silico response correlations

To characterize the pair-wise tuning similarity between two modeled neurons, we computed the Pearson correlation of their responses to 2500 seconds of natural movies. The natural movies were fed in to the model as trials of 10 sec. Model responses were generated at 29.967 Hz and Pearson correlations were computed after binning the responses into 500 msec non-overlapping bins and concatenating across trials.

#### In silico feature weight similarity and receptive field center distance

The digital twin model architecture includes a shared core which is trained to represent spatiotemporal features in the stimulus input, and a final layer where the spatiotemporal features at a specific readout location are linearly weighted in order to produce the predicted activity of a specific neuron at the current time point (Wang et al., 2024). The readout location and linear feature weight are independently learned for each neuron. In order to measure the feature weight similarity between two units, we extract the linear feature weights from this final step as vector of length 512, and take the cosine similarity between the two vectors. In order to measure the receptive field center distance between two units, we extract the readout location as 2D coordinates on the monitor, and take the angle between them with respect to the mouse’s eye, assuming the monitor is centered on, 15 cm away from, and normal to the surface of the mouse’s eye at the closest point.

#### In silico *difference in preferred orientation*

240 blocks of parametric directional visual stimuli (“Monet”) are shown to the model, with each fifteen-second block consisting of 16 trials of equally distributed and randomly ordered unique directions of motion between 0-360 degrees. A modeled neuron’s direction tuning curve is computed as its mean responses to 16 directions averaged across blocks. We calculated the global orientation selectivity index (gOSI) and the orientation selectivity index (OSI) from the modeled neuron’s tuning curve as follows:

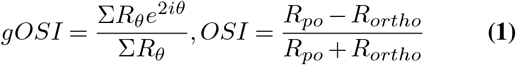

where *θ* is the direction of the stimulus, *R*_*θ*_ is the mean modeled response to the stimulus at direction *θ*, and *R*_*po*_ and *R*_*ortho*_ are the mean modeled responses at the preferred and orthogonal orientation, respectively. The gOSI metric is based on the 1 − *CircV ar* metric in (Mazurek et al., 2014), which is a vector-based method designed to reduce the uncertainty in quantifying orientation selectivity of responses, especially in cases where high throughput, unbiased recording methods return many cells with low orientation selectivity, as is the case with calcium imaging. Only neurons with *gOSI >* 0.25 were included in the analyses in this paper. For neurons selected with our gOSI threshold *>* 0.25, the computed OSI ranges from 0.43 to 0.99, with mean of 0.56. For both thresholds, the fraction of cells considered orientation tuned (57.4% of coregistered V1 neurons has gOSI *>* 0.25, 62.7% of coregistered V1 neurons has OSI *>* 0.4) is similar to those reported in other studies (72% in V1 layer 2/3 (Ko et al., 2011), 62.9% in V1 layer 2/3 and 58.0% in V1 layer 4 (Kondo and Ohki, 2016).Unit-wise direction tuning curves are then modeled by a bivariate von Mises function with an offset:

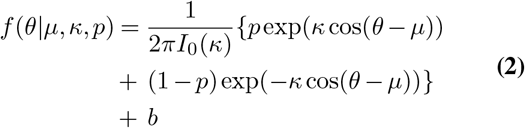

where *I*_0_ is the modified Bessel function, *µ* is the preferred direction, *κ* measures the concentration of the two peaks (larger *κ* means higher peaks thus higher orientation selectivity), *p* measures the relative height of the two peaks (*p* = 0.5 means two peaks of the same height, when *p* approaches 0 or 1, the bi-modal distribution reduces to a uni-model von Mises distribution), *b* is the offset. *µ, κ, p*, and *b* are fit by minimizing least squared error. The preferred orientation of a neuron is taken as the modulus of *µ* to 180 degrees.

### Validation of the digital twin model

#### *Validation of* In silico *signal correlations*

To validate the *in silico* signal correlations generated by our digital twin model, we first established a benchmark for *in vivo* signal correlations. We began by determining the optimal number of stimulus repetitions for measuring *in vivo* signal correlations. Two mice were presented with 6 unique 10-second natural movie clips, each repeated 60 times over a 60-minute period. Based on the results shown in Supplemental Fig. 2a, we determined that 10 repetitions per clip provided a reliable estimate of *in vivo* signal correlation while maintaining a reasonable experimental time for presenting a large number of clips in subsequent experiments.

With this optimal repetition count established, we conducted experiments with three mice using an expanded set of visual stimuli. These stimuli contained those presented in the MICrONS dataset as described in MICrONS Consortium et al. (2021), including natural movies, global directional parametric stimuli (“monet”), and local directional parametric stimuli (“trippy”). Additionally, we presented 36 unique 10-second natural movie clips, each repeated 10 times, totaling 60 minutes of stimulation. To facilitate comparison with the MICrONS dataset and establish a robust ground truth, we divided these 36 clips into two sets: a benchmark set of 30 clips repeated 10 times, serving as our “ground truth” for signal correlation, and a MICrONS-equivalent set of 6 clips repeated 10 times, mimicking the amount of repeated natural clip data available in the MICrONS dataset.

For each mouse, we trained a digital twin model using the same architecture and training data as the MICrONS digital twin. This allowed us to generate three signal correlation matrices for comparison: an *in vivo* matrix computed from the MICrONS-equivalent set, an *in silico* matrix generated by the digital twin model using 250 novel natural movie clips, and a benchmark matrix computed from the 30-clip set. To compare these matrices, we randomly sampled submatrices of signal correlations between 1000 neurons. We then performed hierarchical clustering using Ward’s method on the benchmark matrix and used the resulting dendrogram to sort neurons. This sorting was applied to the MICrONS-equivalent and *in silico* matrices for visual comparison, as shown in Supplemental Fig. 2b. Following this initial comparison, we calculated the Pearson correlation coefficient between the corresponding entries in the lower triangles of the three matrices. To assess statistical significance, we employed a resampling approach, performing 1000 random splits of the benchmark and MICrONS-equivalent sets, from which we estimated the standard deviation and resampling-based p-value of the Pearson correlations. This comprehensive approach enabled us to evaluate how well our digital twin model’s *in silico* signal correlations matched the ground truth compared to *in vivo* measurements with limited data, thus validating the model’s performance in replicating neural response correlations.

#### Validation of receptive field center

To validate the receptive field estimates of our digital twin model, we conducted additional experiments and analyses comparing *in vivo* and *in silico* silico receptive field measurements. We collected three additional functional scans using an expanded set of visual stimuli. These stimuli contained those presented in the MICrONS dataset as described in MICrONS Consortium et al. (2021), including natural movies, global directional parametric stimuli (“monet”), and local directional parametric stimuli (“trippy”). Additionally, we presented 57.6 minutes of sparse noise stimuli. The sparse noise stimuli consisted of bright (pixel value 255) and dark (pixel value 0) square dots, each approximately 6° in visual angle, presented on a grey background (pixel value 127) in a randomized order. These dots were presented at 12 positions covering 70° of visual angle along both the horizontal and vertical axes of the screen. Each presentation lasted 200 ms, and each condition was repeated 60 times.

We computed the *in vivo* spike-triggered average (STA) receptive fields by cross-correlating the visual stimuli with deconvolved calcium traces. STAs for bright dots (on-STAs) and dark dots (off-STAs) were estimated independently and then combined by taking the pixel-wise maximum of the on- and off-STAs. We then presented the same sparse noise stimuli to the digital twin model and computed *in silico* silico STA receptive fields using the model responses. To assess STA quality, we generated response predictions by multiplying each neuron’s STA with the stimulus frames and compared these predictions to either the *in vivo* trial-averaged responses or model responses using Pearson correlation coefficients. Neurons with correlations greater than 0.2 were considered well-characterized. We then extracted the STA receptive field centers by fitting a 2D Gaussian to the STAs, with fits yielding an r-squared value over 0.5 considered well-fit. Our analysis revealed that 40% of all imaged neurons had well-characterized, well-fit *in vivo* STAs.

Finally, we visualized the retinotopic maps measured with either *in vivo* STA or *in silico* STA by converting the STA receptive field centers to azimuth and elevation angles, assuming the mouse was looking at the center of the monitor. To exclude partially measured STAs, we included only neurons with fitted STA centers located in the central 8×8 square of the entire 12×12 stimulus grid (27% of all imaged neurons) for the analysis presented in Supplemental Fig. 4 a, b, and c left. For the analysis in Supplemental Fig. 4 c right, we included neurons with the bottom 25% of response correlations.

#### Validation of orientation tuning

To validate *in silico* orientation tuning with *in vivo* orientation tuning, we collected three additional functional scans with an expanded set of stimuli. These stimuli contained those presented in the MICrONS dataset as described in MICrONS Consortium et al. (2021), including natural movies, global directional parametric stimuli (“monet”), and local directional parametric stimuli (“trippy”). In addition, each stimulus contained an additional 40 minutes of trials, randomly intermixed, as follows:

- **Unique Global Directional Parametric Stimulus (“Monet”):** 120 seeds, 15 seconds each, 1 repeat per scan, 30 minutes total. Seeds conserved across all scans.
- **Oracle Global Directional Parametric Stimulus (“Monet”):** 4 seeds, 15 seconds each, 10 repeats, 10 minutes total. Seeds conserved across all scans.

We characterized both the *in vivo* orientation tuning in response to 30 minutes of global directional parametric stimulus (“Monet”, Supplemental Fig. 7a), as well as the *in silico* orientation tuning as described above for digital twin models with shared cores and readouts trained on neurons from the same scans, in response to stimuli matching the composition and duration of the MICrONS release scans (Supplemental Fig. 7b). When we applied a threshold of *gOSI >* 0.25, we found that 95% of cells had an absolute difference between their *in silico* and *in vivo* preferred orientations less than 9.77°.

### Statistical analysis of mean signal correlations

We employed paired t-tests to compare signal correlations between presynaptic neurons and three groups of potential target neurons: connected postsynaptic neurons, axon-dendrite proximity (ADP) neurons, and same-region control neurons. Our analysis focused on presynaptic neurons with more than 10 postsynaptic targets for each projection type to ensure robust comparisons. For each presynaptic neuron, we computed mean signal correlations with its synaptically connected postsynaptic targets, ADP neurons (neurons with dendrites in proximity to the presynaptic axon but not synaptically connected), and same-region control neurons (neurons in the same brain region but without proximal axon-dendrite contacts). We then performed paired t-tests to compare these mean correlations. For example, to compare connected and ADP neuron pairs, we conducted a paired t-test between each presynaptic neuron’s mean signal correlation with its postsynaptic targets versus its mean signal correlation with ADP neurons. This approach allowed us to control for variability across presynaptic neurons while directly comparing their correlations with different target groups. All statistical analyses were performed using the `scipy` package in Python. We set the significance level (α) at 0.05 for all tests. To account for multiple comparisons, we adjusted p-values using the Benjamini-Hochberg (BH) procedure as implemented in the `statsmodels` package.

### Visualization of the relationship between *L*_*d*_, *N*_*syn*_*/L*_*d*_ and the functional similarities

#### Visualization of L_d_

To quantify the changes in *L*_*d*_ as a function of functional similarities, we restrict our analysis to neuron pairs with no synaptic connections observed between them. We then follow these steps:

1. We compute the mean *L*_*d*_ and mean functional similarities for each presynaptic neuron across all other neurons that no synaptic connections with the presynaptic neuron were observed.
2. We subtract the presynaptic mean from each of the pairwise *L*_*d*_ and functional similarities between every neuron pair to compute Δ*L*_*d*_ and Δ*similarity*.
3. The neurons pairs are then binned by Δ*similarity* and the average Δ*L*_*d*_ is computed for each bin.
4. The standard deviation of average Δ*L*_*d*_ is estimated by bootstrapping. Specifically, we resampled the neuron pairs 1000 times with replacement and repeated steps 1-3.

Only bins with more than 10 connected neuron pairs and more than 10 presynaptic neurons are included in the visualization.

#### Visualization of N_syn_/L_d_

To quantify the changes in *N*_*syn*_*/L*_*d*_ as a function of functional similarities, we restrict our analysis to neuron pairs with positive *L*_*d*_ observed between them. We then follow these steps:

1. We compute the mean *N*_*syn*_*/L*_*d*_ and mean functional similarities for each presynaptic neuron across all other neurons that no synaptic connections with the presynaptic neuron were observed.
2. We subtract the presynaptic mean from each of the pairwise *N*_*syn*_*/L*_*d*_ and functional similarities between every neuron pair to compute Δ*N*_*syn*_*/L*_*d*_ and Δ*similarity*.
3. The neurons pairs are then binned by Δ*similarity* and the average Δ*N*_*syn*_*/L*_*d*_ is computed for each bin.
4. The standard deviation of average Δ*L*_*d*_ is estimated through bootstrapping. Specifically, we resampled the neuron pairs 1000 times with replacement and repeated steps 1-3.

Only bins with more than 10 connected neuron pairs and more than 10 presynaptic neurons are included in the visualization.

### Statistical modeling of “like-to-like” rules for different anatomical measurements

#### Axon-dendrite co-travel distance (L_d_)

*L*_*d*_ measures the distance dendrites of one neuron travel within 5 *µm* from another neuron’s axon. Most pairs of neurons’ axons and dendrites never come into close proximity with each other, and their *L*_*d*_ is zero. Thus, the *L*_*d*_ distribution is a non-negative continuous distribution with a substantial non-zero probability measure at zero *L*_*d*_. Thus, we modeled *L*_*d*_ as a random variable following the Tweedie exponential dispersion family (with Tweedie index parameter *ξ* ∈ (1, 2)). Tweedie distributions with such index parameters are Poisson mixtures of gamma distributions, commonly used to model continuous data with exact zeros. We assume two neurons’ axons and dendrites travel within 5 *µm* at *N* proximity points, where *N* ∼ Pois(*λ*\*), *λ*\* is the mean number of axonal dendritic proximal contacts of the Poisson distribution. When *N >* 0, we assume the distance dendrites travel within five *µm* at each proximal point *z*_*i*_ (*i* = 1,…, *N*) follows a Gamma distribution Gam(*µ,ϕ*). Under these assumptions, the total potential synapsing distance

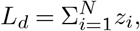

where *L*_*d*_ = 0 when *N* = 0, follows a Tweedie distribution with 1 *< ξ <* 2. We then model the relationship between *L*_*d*_, functional similarities *Sim* (e.g., signal correlation, feature weight similarity, receptive field location distance between two neurons), and projection types *Proj* using a Tweedie-distributed generalized linear mixed model (GLMM) with a log link function. For analysis at the brain area level, *Proj* is a nominal variable with 4 categories: V1 intra-area, HVA intra-area projections, feedforward projections, and feedback projections. For analysis at the brain area and layer level, we apply GLMMs for modeling as they have been recommended for accounting for multi-level data dependencies in datasets (Yu et al., 2022), such as the projection types and presynaptic neuron proofreading progress in our study. We specify the model as follows:

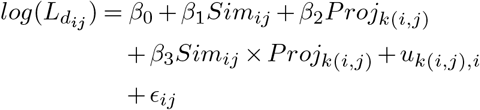

where:

- 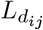 is the axon-dendrite co-travel distance between presynaptic neuron *i* and postsynaptic neuron *j*
- *Sim*_*ij*_ is the functional similarity between the neuron pair
- *Proj*_*k*(*i,j*)_ is the projection type of the neuron pair (*i, j*)
- *β*_1_, *β*_2_, and *β*_3_ are the fixed effect coefficients of the functional similarity, projection type, and their interaction term, respectively
- *β*_0_ is the intercept
- *u*_*k*(*i,j*),*i*_ is the random effect accounting for the projection type *k* and the proofread status associated with presynaptic neuron *i*
- *ϵ*_*ij*_ is the error term, following a Tweedie distribution

The coefficients *β*_1_, *β*_2_, and *β*_3_ represent how functional similarities and projection types affect connectivity at the axonal scale. We fit the models for each functional similarity independently using the `glmmTMB` R package. The goodness-of-fit of the estimated models is reported as Nakagawa’s R squared, computed with the `performance` R package. We define the axonal-scale like-to-like coefficients for each functional similarity and projection type as the estimated linear association between each category of functional similarity conditioned on the projection type. The coefficient estimates and the corresponding significance tests are computed for the fitted GLMM using the `emtrends` function from the `emmeans` R package.

#### Number of synapses (N_syn_)

*N*_*syn*_ measures the number of synapses between two neurons. We model it as a *P oisson*-distributed random variable and its relationship to functional similarities as a GLMM model with the following specifications:

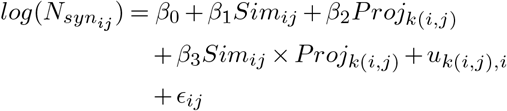

where:

- 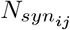 is the number of synapses between presynaptic neuron *i* and postsynaptic neuron *j*
- *Sim*_*ij*_ is the functional similarity between the neuron pair
- *Proj*_*k*(*i,j*)_ is the projection type of the neuron pair (*i, j*)
- *β*_1_, *β*_2_, and *β*_3_ are the fixed effect coefficients of the functional similarity, projection type, and their interaction term, respectively
- *β*_0_ is the intercept
- *u*_*k*(*i,j*),*i*_ is the random effect accounting for the projection type *k* and the proofread status associated with presynaptic neuron *i*
- *ϵ*_*ij*_ is the error term, following a Poisson distribution

The coefficients *β*_1_, *β*_2_, and *β*_3_ estimate how the functional similarities and projection types affect connectivity regardless of the spatial scales (i.e., axonal or synaptic). We fit the models for each functional similarity independently using the `glmmTMB` R package. The goodness-of-fit of the estimated models is reported as Nakagawa’s R squared, computed with the `performance` R package. We define the axonal-scale like-to-like coefficients for each functional similarity and projection type as the estimated linear association between each category of functional similarity conditioned on the projection type. The coefficient estimates and the corresponding significance tests are computed for the fitted GLMM using the `emtrends` function from the `emmeans` R package.

#### Synapse conversion rate (N_syn_/L_d_)

*N*_*syn*_*/L*_*d*_ measures the number of synapses per millimeter axon-dendrite co-travel distance for each neuron pair. To quantify its relationship to functional similarities, we adopted the following GLMM model:

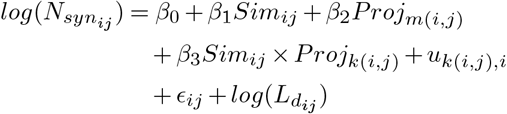

where:

- 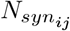 is the number of synapses between presynaptic neuron *i* and postsynaptic neuron *j*
- 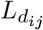 is the axon-dendrite co-travel distance between the neuron pair
- *Sim*_*ij*_ is the functional similarity between the neuron pair
- *Proj*_*m*(*i,j*)_ is the projection type of the neuron pair (*i, j*)
- *β*_1_, *β*_2_, and *β*_3_ are the fixed effect coefficients of the functional similarity, projection type, and their interaction term, respectively
- *β*_0_ is the intercept
- *u*_*k*(*i,j*),*i*_ is the random effect accounting for the projection type *k* and the proofread status associated with presynaptic neuron *i*
- *ϵ*^*ij*^ is the error term, following a Poisson distribution

The above equation can be re-arranged to:

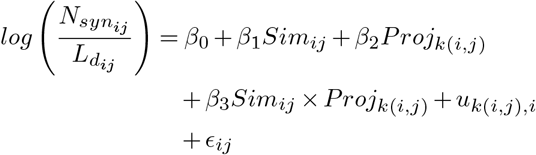

Thus, *β*_1_, *β*_2_, and *β*_3_ model how the functional similarities affect synapse conversion rate (*N*_*syn*_*/L*_*d*_) at the synaptic scale. We fit the models for each functional similarity independently using the `glmmTMB` R package. The goodness-of-fit of the estimated models is reported as Nakagawa’s R squared, computed with the `performance` R package. We define the like-to-like coefficients of each functional similarity for each projection type as the estimated linear association between each category of functional similarity conditioned on the projection type. The coefficient estimates and the corresponding significance tests are computed for the fitted GLMM using the `emtrends` function from the `emmeans` R package. To avoid fitting models to projection types with little data or dominated by few presynaptic neurons, for all the models described above, we only include and report like-to-like coefficients to projection types with more than 30 synapses observed, more than 5 presynaptic neurons, and with none of the presynaptic neurons contributing more than half of all synapses observed.

### Statistical analysis of functional similarities and synaptic anatomy

We investigated the relationship between functional similarities of neurons and the anatomical features of their synaptic connections. Our analysis accounted for the confounding effect of axon-dendrite co-travel distance (*L*_*d*_), which correlates with both functional similarities and synaptic measurements. To isolate the effect of synaptic anatomy on functional similarity, we employed a two-step regression approach:

First, we condition our analysis on the effect of *L*_*d*_ from the functional similarity measure. This process involves:

1. Fitting a linear regression model with functional similarity as the dependent variable and *L*_*d*_ as the independent variable.
2. Calculating the residuals from this model, which represent the variation in functional similarity that cannot be explained by *L*_*d*_ alone.

These residuals become our new measure of functional similarity, adjusted for the influence of *L*_*d*_. Next, we constructed a linear regression model using these residuals as the dependent variable. The independent variables in this model included anatomical measurements of synaptic connections, the total number of synapses between neuron pairs and the mean synaptic cleft volume.

This approach allows us to test whether synaptic measurements significantly predict functional similarities between neurons, beyond what can be explained by their physical proximity (as measured by *L*_*d*_).

### Common input analysis

#### Functional similarity among all postsynaptic neurons sharing one common input

For a connectivity graph *G*, we define

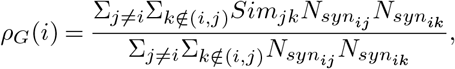

where *i* is a presynaptic neuron and *j, k* are any two neurons in the volume. *ρ* measures the average similarity of all post-synaptic neurons of the presynaptic neuron *i*.

#### Estimation of ρ expected by pairwise “like-to-like” connectivity rules

With the observed connectivity graph *G*, we estimated the relationship between *N*_*syn*_ and the functional similarities (in silico signal correlation, feature weight similarity, and receptive field center distance) with GLMM similar to the specifications for modeling the number of synapses described above. Instead of modeling each functional similarity independently, we included all functional similarities and their interaction with projection types in a single model to account for as much pairwise connectivity rule as possible. We then estimated the expected functional similarity among all postsynaptic neurons sharing one common input *i* as:

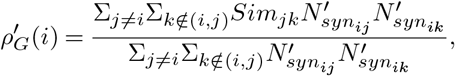

where 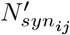 is the predicted number of synapses between neurons *i* and *j* given their functional similarities by the GLMM.

### RNN model

The RNN model used to produce the results in Fig. 6 consisted of a vanilla RNN layer with 1000 hidden units and a hyperbolic tangent activation function simulated over 20 time steps. Static inputs were obtained by passing MNIST images through a linear layer. Outputs were obtained by passing the hidden activations at the last time step through another linear layer. All three layers were trained for 10 epochs, a batch size of 512, the categorical cross entropy loss function, and the Adam optimizer in PyTorch. A pre- and post-synaptic neuron pair was classified as connected if the associated weight was in the top 35^th^ percentile of all weights, specifically if the weight larger than 0.01. In Fig. 6d, weights were chosen as candidates for ablation if the weight was above 0.01 and the neurons’ signal correlation was above 0.2. About 10.5% of the weights met these criteria, and ablated weights were selected randomly from this set. Changing the thresholds for weights and signal correlations did not change our conclusions.

### Software

Experiments and analysis are carried out with custom built data pipelines. The data pipeline is developed in Matlab, Python, and R with the following tools: Psychtoolbox, ScanImage, DeepLabCut, CAIMAN, and Lab-view were used for data collection. DataJoint, MySQL, and CAVE were used for storing and managing data. Meshparty, NEURD, and pcg_skel were used for morphology analysis. Numpy, pandas, SciPy, statsmodels, scikit-learn, PyTorch, tidyverse, glmmTMB, performance, and emmeans were used for model training and statistical analysis. Matplotlib, seaborn, HoloViews, Ipyvolume, and Neuroglancer were used for graphical visualization. Jupyter, Docker, and Kubernetes were used for code development and deployment.

## Data availability

All MICrONS data have already been released on BossDB (https://bossdb.org/project/microns-minnie, please also see https://www.microns-explorer.org/cortical-mm3 for details). Additional data including learned weights of the digital twin model and *in silico* similarity metrics will be made publicly available in an online repository latest upon journal publication. Please contact the corresponding authors for advance access.

## Code availability

Custom developed code used in the analysis including digital twin architecture will be made publicly available in an online repository latest upon journal publication. Please contact the corresponding authors for advance access.

---

**Supplemental Figure 1.**
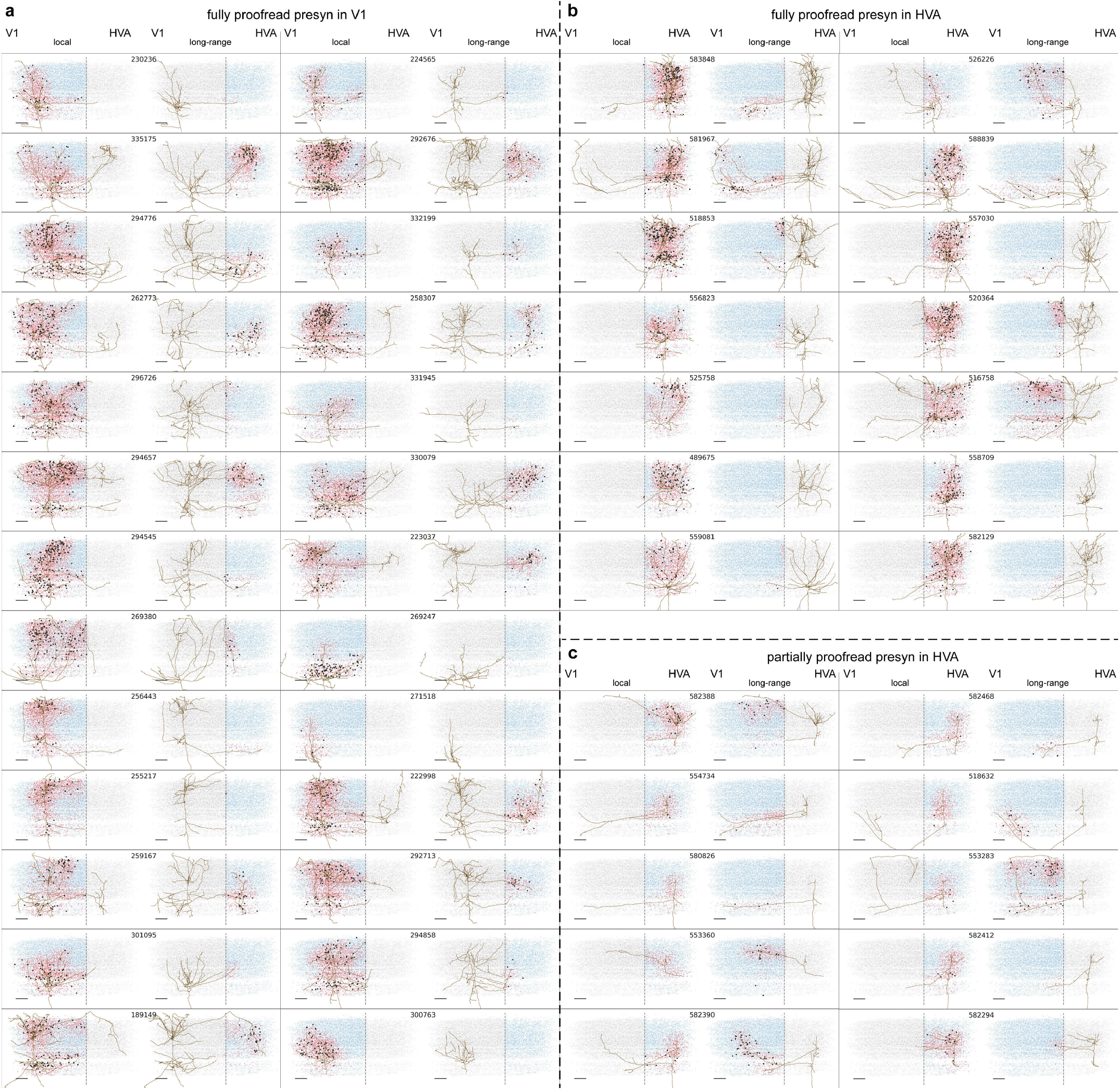
Example proofread presynaptic axons in EM cortical space and their connected, ADP, and same region controls. The axon for every presynaptic (presyn) neuron is shown twice, once as a “local” projection type and again as a “long-range” type (even if the neuron has no local or long-range projections). The six digit ID from Table “nucleus_detection_v0” (MICrONS Consortium et al., 2021) is displayed above both plots. For each plot, the soma centroids of connected neurons, ADP controls, and same region controls are plotted in black, red, and blue, respectively. Gray dots are soma centroids of all other functionally matched neurons not used as controls for that presyn. The dashed gray line represents the V1-HVA boundary. Scale bar = 100*µm*. **a**, Example fully proofread presynaptic axons with somas in V1. “Fully proofread” neurons are those where a proofreader attempted to extend every axonal branch to completion. **b**, Example fully proofread presynaptic axons with somas in HVA **c**, Example partially proofread presynaptic axons with somas in HVA. “Partially proofread” neurons are those where a proofreader only extended axonal branches that were pre-screened for whether they projected inter-areally (specifically to enrich for feedback connections).

**Supplemental Figure 2.**
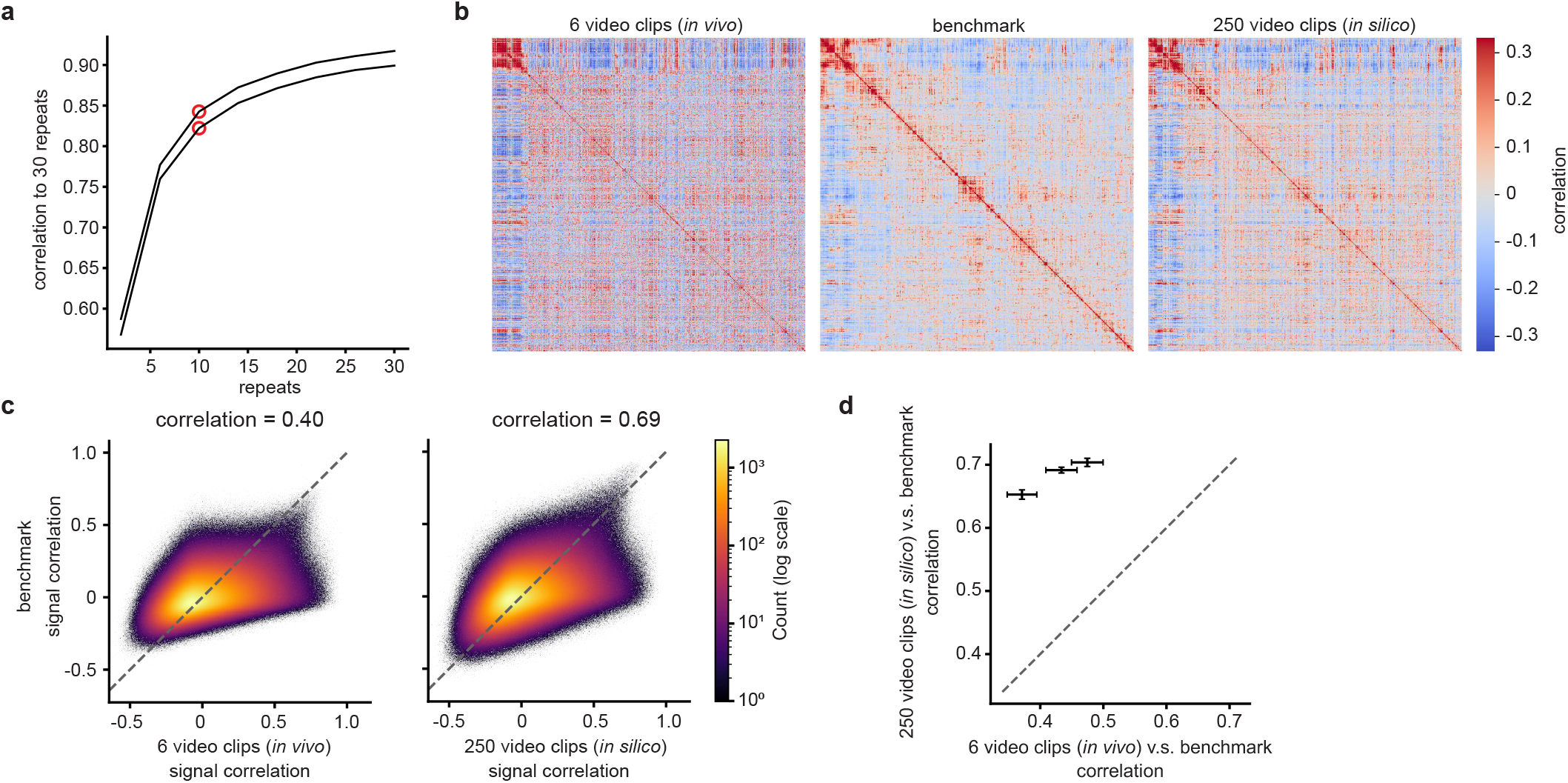
The digital twin signal correlations align better with the *in vivo* benchmark than *in vivo* signal correlations generated with less data. **a**, Correlation of *in vivo* signal correlations generated with 6 video clips and varying numbers of repeats to *in vivo* signal correlations generated with 6 clips and 30 repeats, for two animals. 10 repeats (red marker) reasonably approximates the saturation point and is the number used for all other analyses. **b**, Signal correlation matrices of 1000 neurons generated from *in vivo* responses to 6 video clips (left), *in vivo* responses to 30 video clips (benchmark, middle) and digital twin responses to 250 video clips (*in silico*, right). The benchmark matrix is ordered by ward’s hierarchical clustering. The *in vivo* and *in silico* signal correlation matrices are ordered in the same order as the benchmark matrix. The fine structure of the *in silico* matrix is qualitatively more similar to the benchmark than the *in vivo* matrix generated with 6 video clips is to the benchmark. **c**, 2D heatmaps of signal correlations from the benchmark (same benchmark as in **b**) vs *in vivo* responses to 6 video clips (left) and *in silico* responses to 250 clips (right). The correlation of *in silico* signal correlations to the benchmark is higher than the correlation of *in vivo* signal correlations generated with 6 video clips to the benchmark (0.69 vs 0.40). Colorbar: 2D bin counts in log scale. **d**, The correlation of *in silico* signal correlations to the benchmark vs the correlation of *in vivo* signal correlations generated with 6 video clips to the benchmark for three animals. Error bars are standard deviations estimated through resampling. All data points are in the upper left corner indicating that *in silico* signal correlations outperform *in vivo* signal correlations generated with 6 video clips. (p-value < 0.001 for all three animals)

**Supplemental Figure 3.**
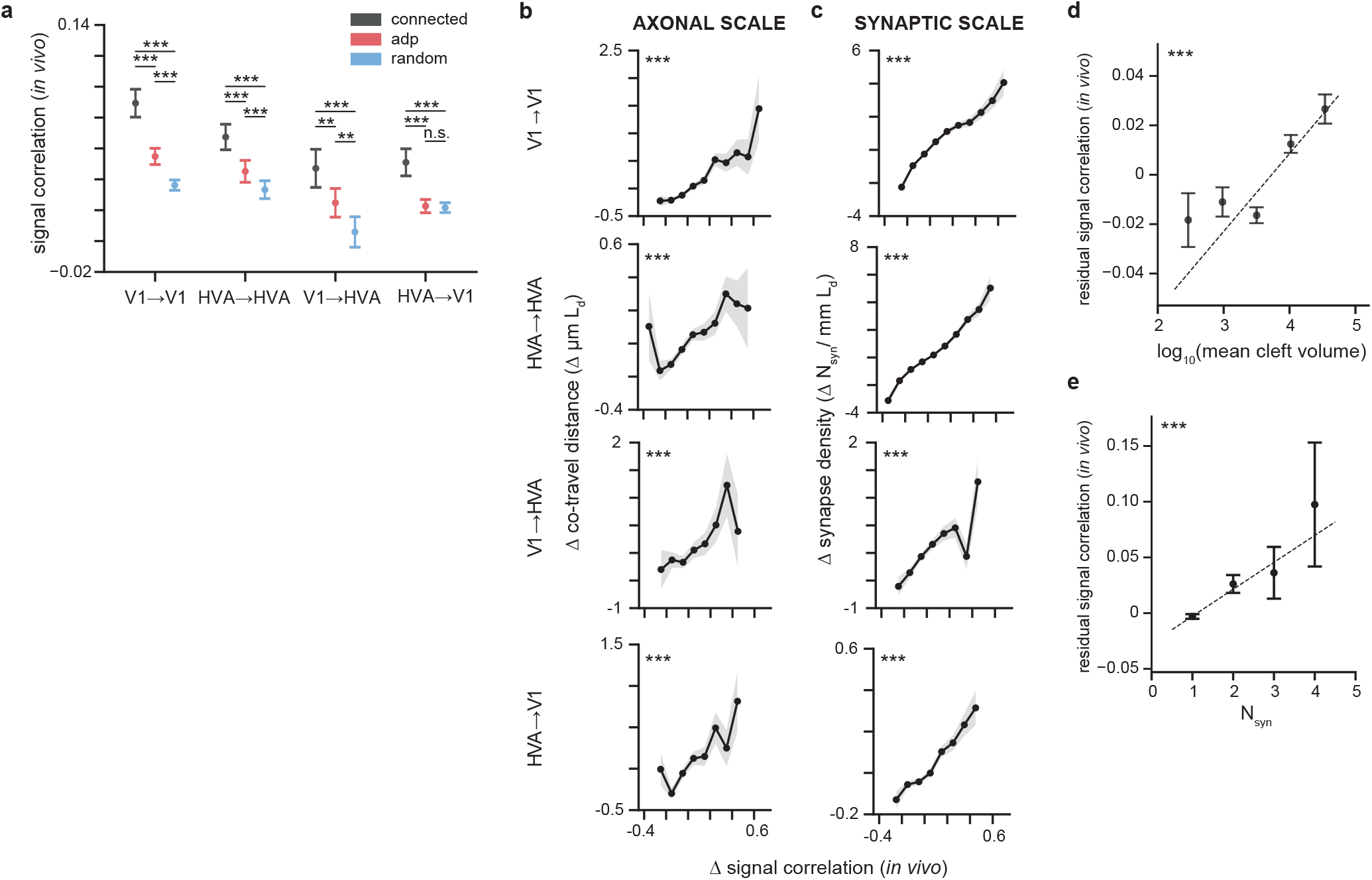
Synaptic connectivity increases with empirical signal correlations measured directly *in vivo* rather than via the digital twin. **a**, Mean *in vivo* signal correlation is different (mean *±* sem, paired t-test) for connected pairs, ADP controls, and same area controls for all projection types, as in Fig 2d. **b**, Axon-dendrite co-travel distance (*µmL*_*d*_) increases in a graded fashion with *in vivo* signal correlation for all projection types, as in Fig 2e. **c** Synapse density (*N*_*syn*_*/mmL*_*d*_) increases in a graded fashion with signal correlation, for all projection types, as in Fig 2f. The shaded regions in **b** and **c** are bootstrap-based standard deviation. **d**, Synapse size (*log*_10_ cleft volume in voxels) is positively correlated with *in vivo* signal correlation after regressing out *L*_*d*_ (p-value by linear regression), as in Fig 2h. **e**, *In vivo* signal correlations increases with number of synapses after regressing out *L*_*d*_ (p-values by linear regression), as in Fig 2j. (For all panels, *\** = p-value < 0.05, *\*\** = p-value < 0.01, *\* * \** = p-value < 0.001, multiple comparison correction by BH procedure)

**Supplemental Figure 4.**
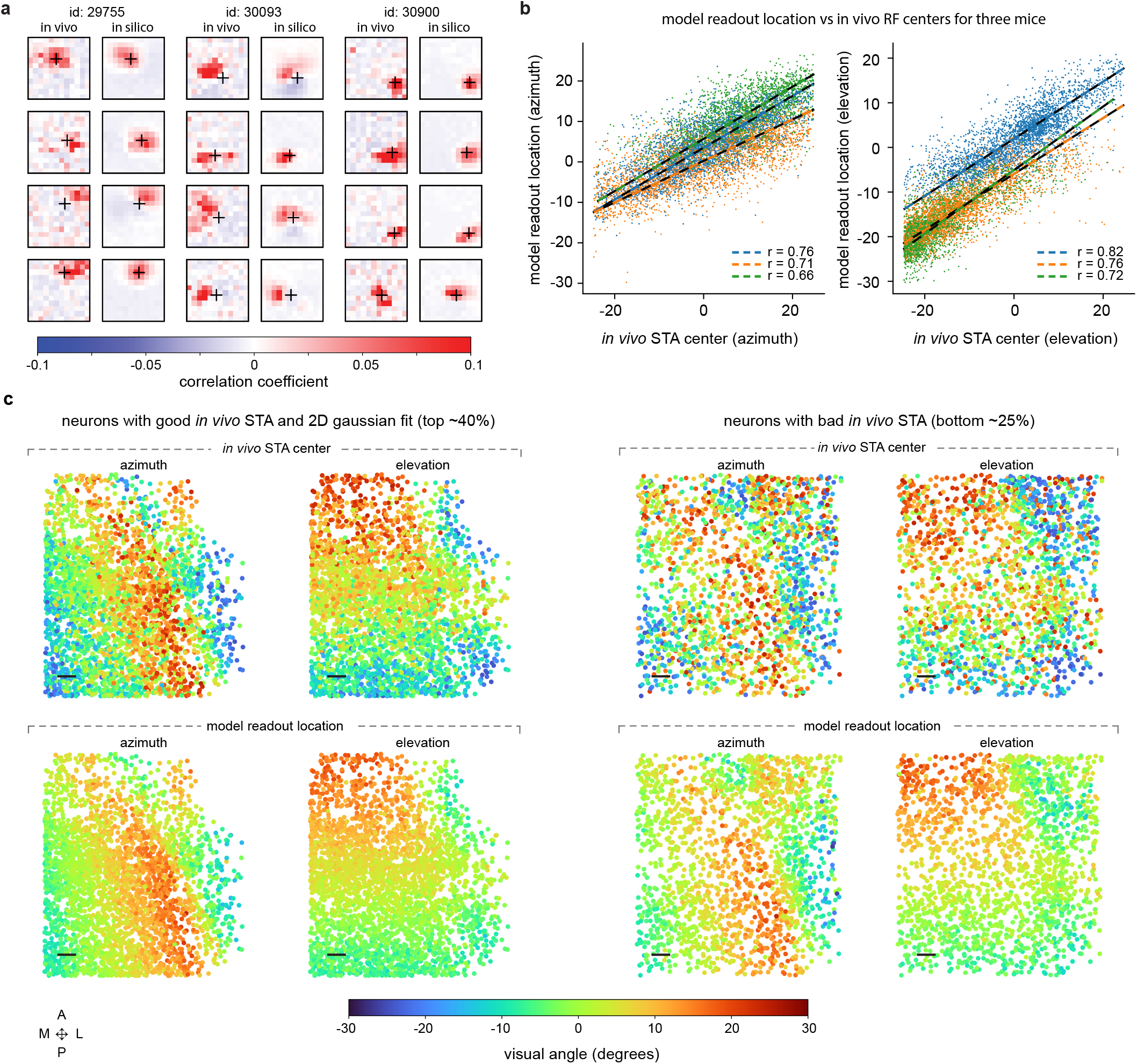
Model readout center aligns with receptive field center measured *in vivo* with sparse noise stimuli. **a**, Visual comparison of STAs generated from *in vivo* responses to a sparse noise stimulus (left) vs STAs generated from *in silico* responses to the same stimulus (right) for three animals (blue, orange, and green). The black cross represents the model readout location. Examples are randomly chosen from the top ≈ 40% of neurons remaining after a threshold on *in vivo* STA quality is applied. **b**, Model readout location vs *in vivo* STA center for azimuth coordinate (left) and elevation coordinate (right). **c**, Retinotopic maps for animal id: 29755. Left: Maps generated with top ≈ 40% of neurons after an *in vivo* STA quality threshold is applied. Right: Maps for the bottom ≈ 25% of neurons. Top row: maps generated from *in vivo* STA’s centers. Bottom row: maps generated from the digital twin model readout location. The maps generated from the model are qualitatively less noisy, even for maps generated from neurons with poor STA quality. Colorbar: degree of visual angle for both azimuth and elevation coordinates. Anatomical axes: A = anterior, P = posterior, M = medial, L = lateral. Scale bar: 100 *µm*.

**Supplemental Figure 5.**
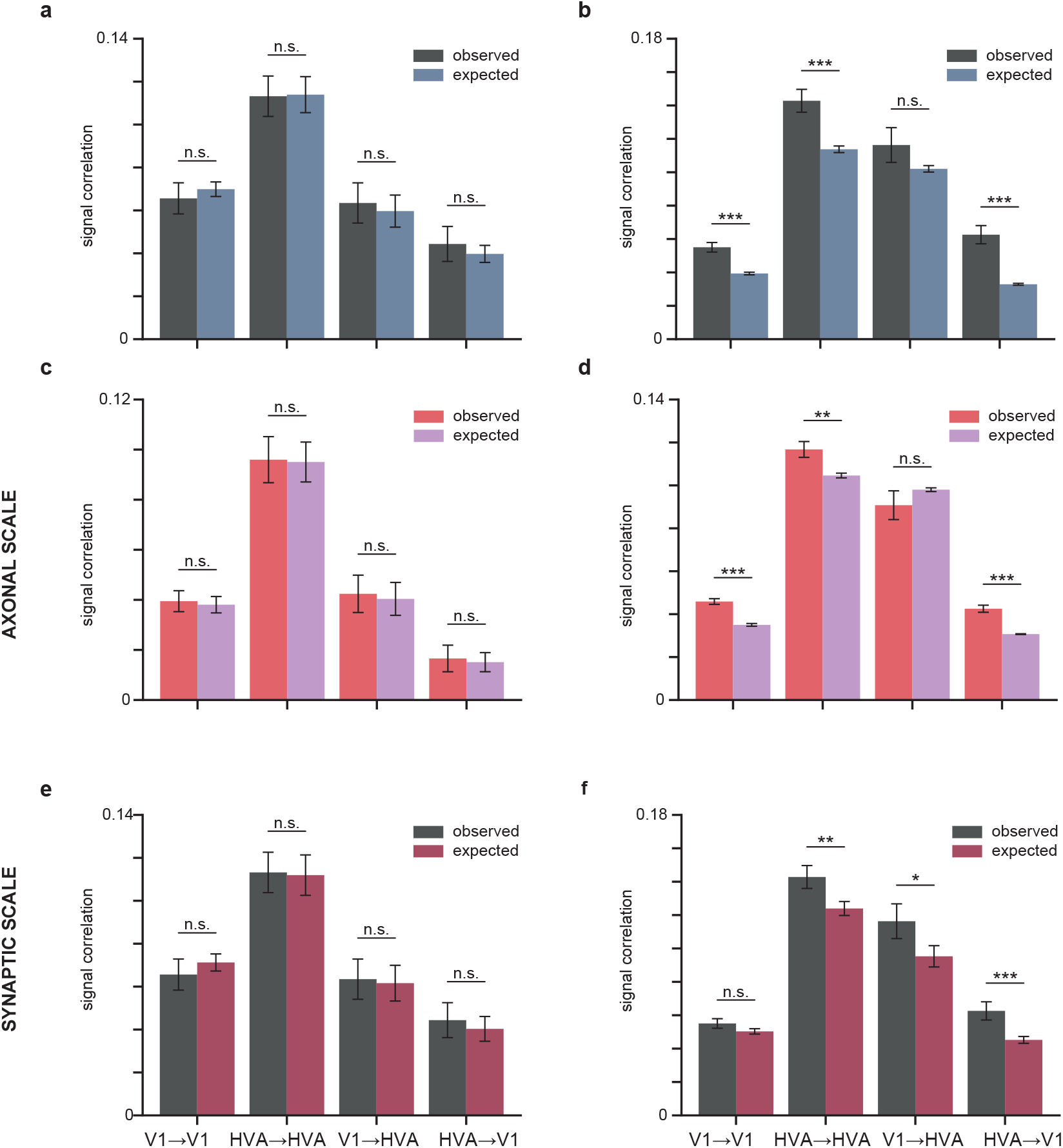
Postsyns with a common input are more similar to each other than expected by a pairwise like-to-like rule at both axonal and synaptic scale. **a**, Mean pre-post signal correlations in the data (dark gray, “observed”) and the model (blue, “expected”) are not significantly different, indicating that the model reproduces the expected pairwise like-to-like rule **b**, Mean pairwise *in silico* signal correlation of postsyns, reproduced from Fig 5c. The observed data shows significantly higher postsyn to postsyn similarity than predicted by the model fit with only a pairwise rule, for three out of four projection types. **c**, As in **a**, but at “Axonal” scale. **d**, As in **b**, but at “Axonal” scale. **e**, As in **c**, but at “Synaptic” scale. **f**, As in **d**, but at “Synaptic” scale.

**Supplemental Figure 6.**
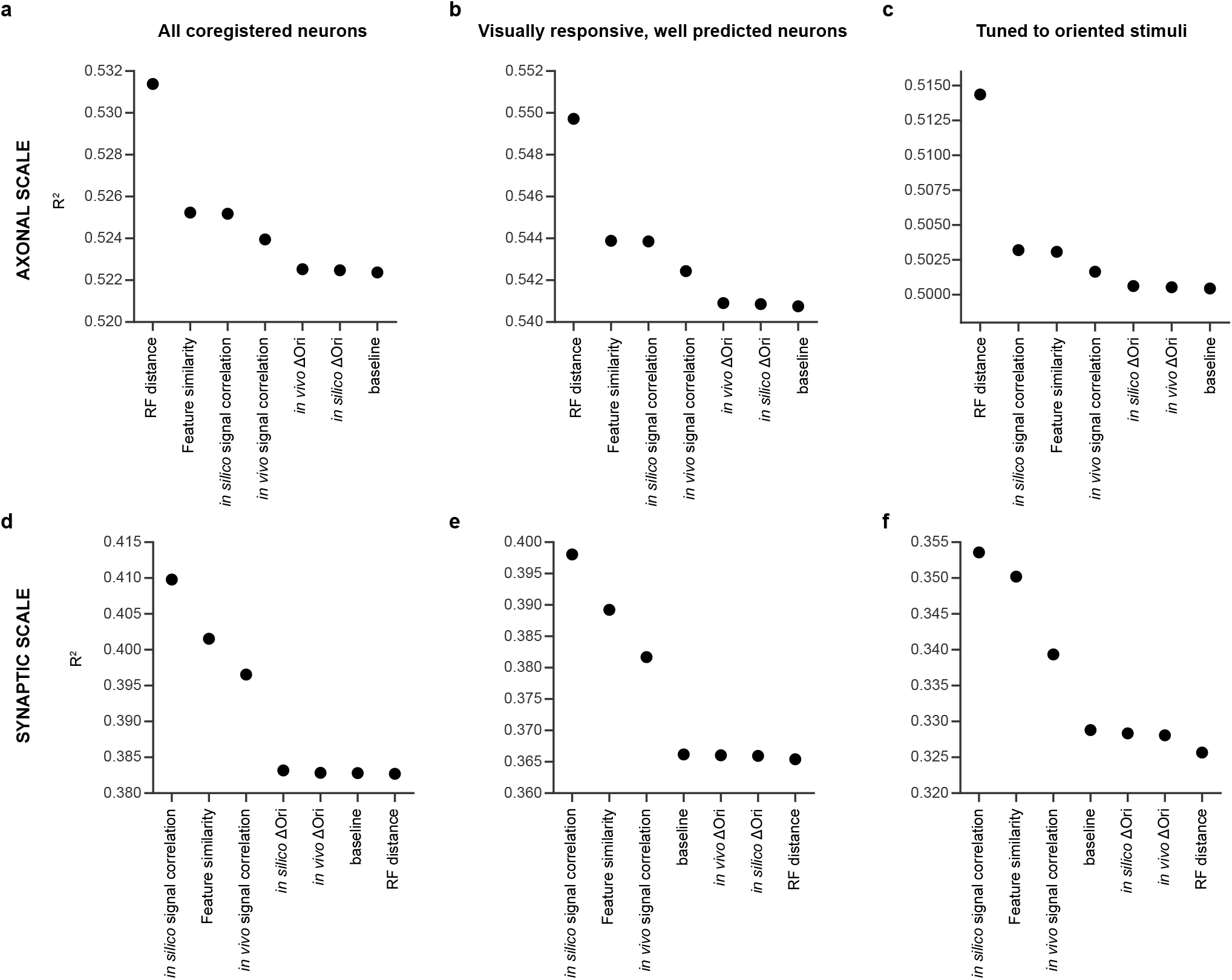
Performance of various functional metrics in predicting axon-dendrite co-travel distance (*L*_*d*_, Axonal scale) or synapse density (*N*_*syn*_*/mmL*_*d*_, Synaptic scale). Model performance of GLMMs (Nakagawa’s conditional *R*^2^) for predicting axon-dendrite co-travel distance (*L*_*d*_): **a, b, c** and synapse density (*N*_*syn*_*/mmL*_*d*_): **d, e, f**, for all coregistered neurons: **a, d**, all visually responsive, well predicted neurons: **b, e**, and neurons tuned to oriented stimuli: **c, f**. The GLMMs are fit to predict axon-dendrite co-travel distance or synapse density independently with each functional metric, the projection type, and the interaction between the two while considering the interaction term of projection type and presynaptic neuron identity as random effects. The baseline models were not fitted with information about functional metrics. They predict axon-dendrite co-travel distance or synapse density with the projection type alone while considering the interaction term of projection type and presynaptic neuron identity as random effects.

**Supplemental Figure 7.**
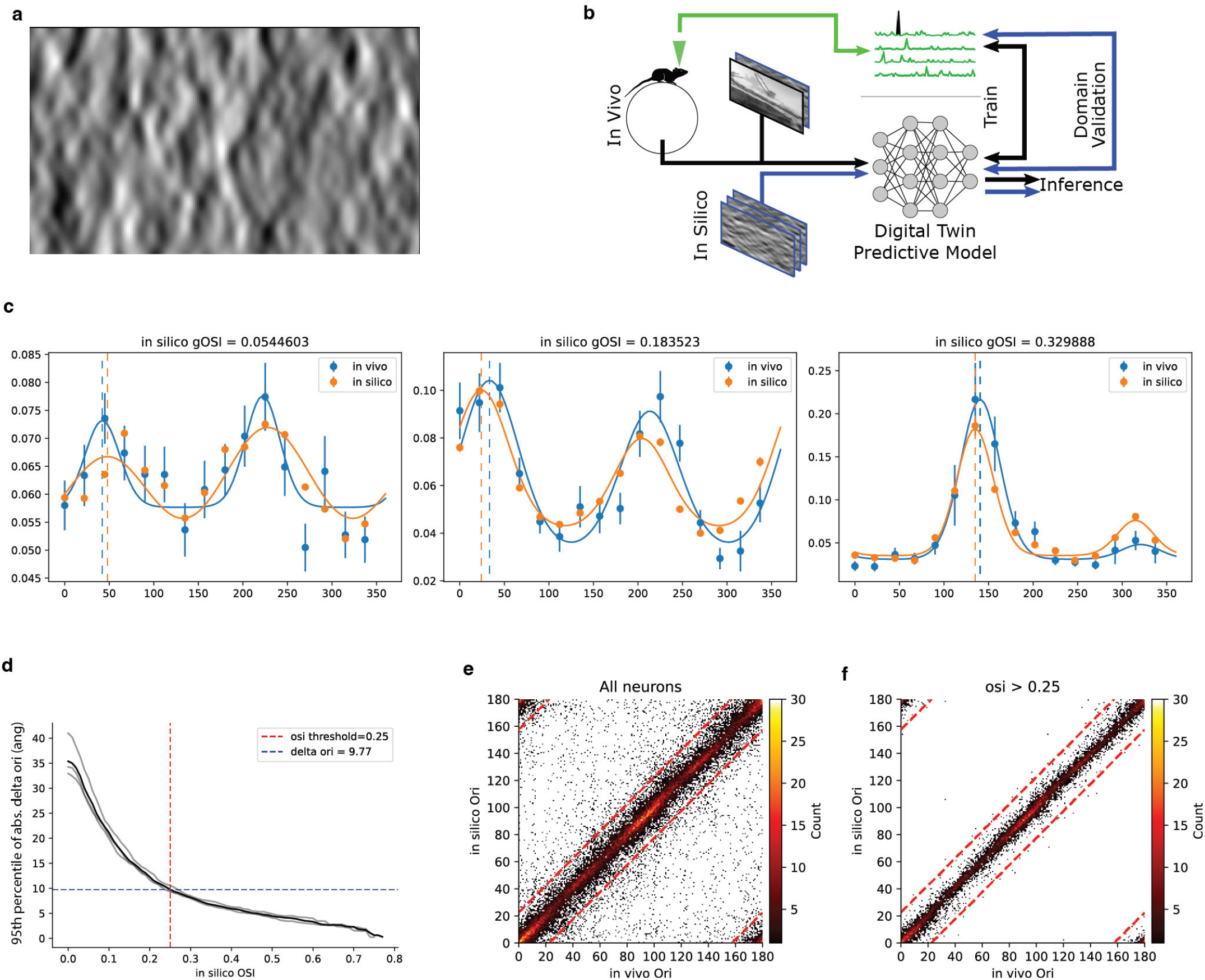
In silico orientation tuning is consistent with in vivo orientation tuning. **a**, Sample frame from global directional parametric stimulus (“Monet”) used to characterize orientation and direction selectivity. Directional motion was orthogonal to orientation, and was tested at 22.5°intervals. **b**, Schematic of domain validation experimental design. In a single scan in a new animal, neuronal responses are collected in response to sufficient stimuli to both train the digital twin model (natural stimuli) and characterize orientation tuning (Monet) from *in vivo* responses. Later, *in silico* orientation tuning is extracted from model responses to parametric stimuli, and compared against *in vivo* orientation tuning for the same neurons. **c**, Comparison of *in silico* and *in vivo* mean responses per stimulus direction (mean *±* SEM), fitted tuning curves (lines), and extracted preferred orientation (dotted lines) for three neurons. **d**, 95th percentile difference in preferred orientation between *in silico* and *in vivo* fitted responses as a function of gOSI threshold. Dotted lines correspond to gOSI > 0.25 threshold applied for all analyses and resulting 95th percentile difference in preferred orientation ≈ 9.77° across all three animals imaged. Lines correspond to individual animals (gray) or cumulative across all animals (black). **e, f**, Two-dimensional histogram of *in silico* versus *in vivo* preferred orientation for all neurons across three animals (**e**) and only neurons with gOSI > 0.25 (**f**).

**Supplemental Figure 8.**
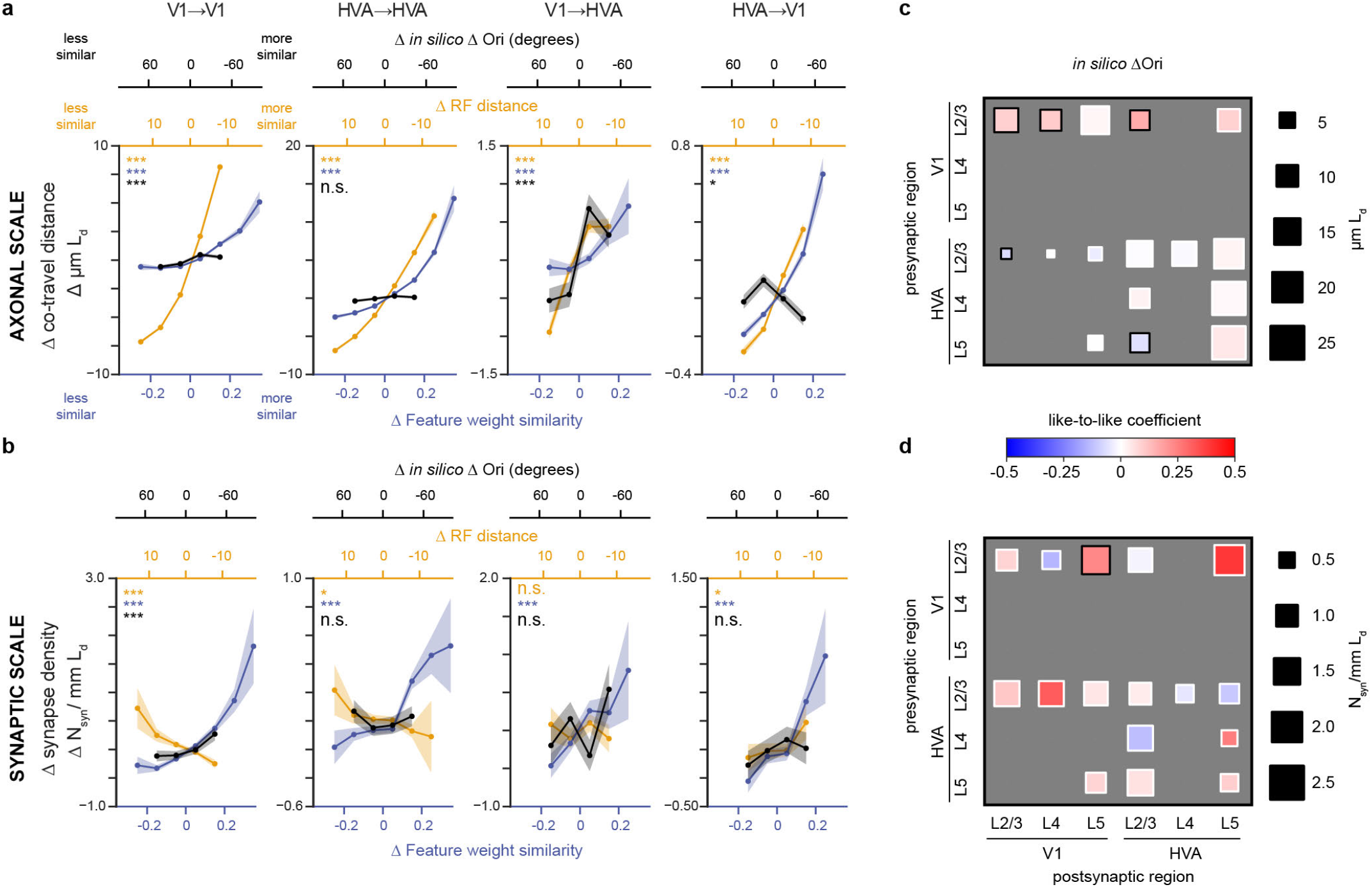
Analysis repeated with *in silico* orientation preference. **a**, Difference in preferred orientation (Δ Ori) derived from *in silico* responses to parametric stimuli for tuned (gOSI > 0.25) neurons along with both feature weight similarity and receptive field center distance (reproduced from Fig 3) at axonal scale. **b**, same as in **a**, at synaptic scale. **c**, Area/ layer joint membership breakout as in Fig 4 for *in silico* Δ ori at axonal scale. **d**, As in **c** but at synaptic scale. All analyses are centered per presyn by accounting for the presyn mean (e.g. Δ feature weight similarity). For details, see Supplemental Tab. 13, 14, 17, 18, 21, 22, 31, 32,

**Supplemental Figure 9.**
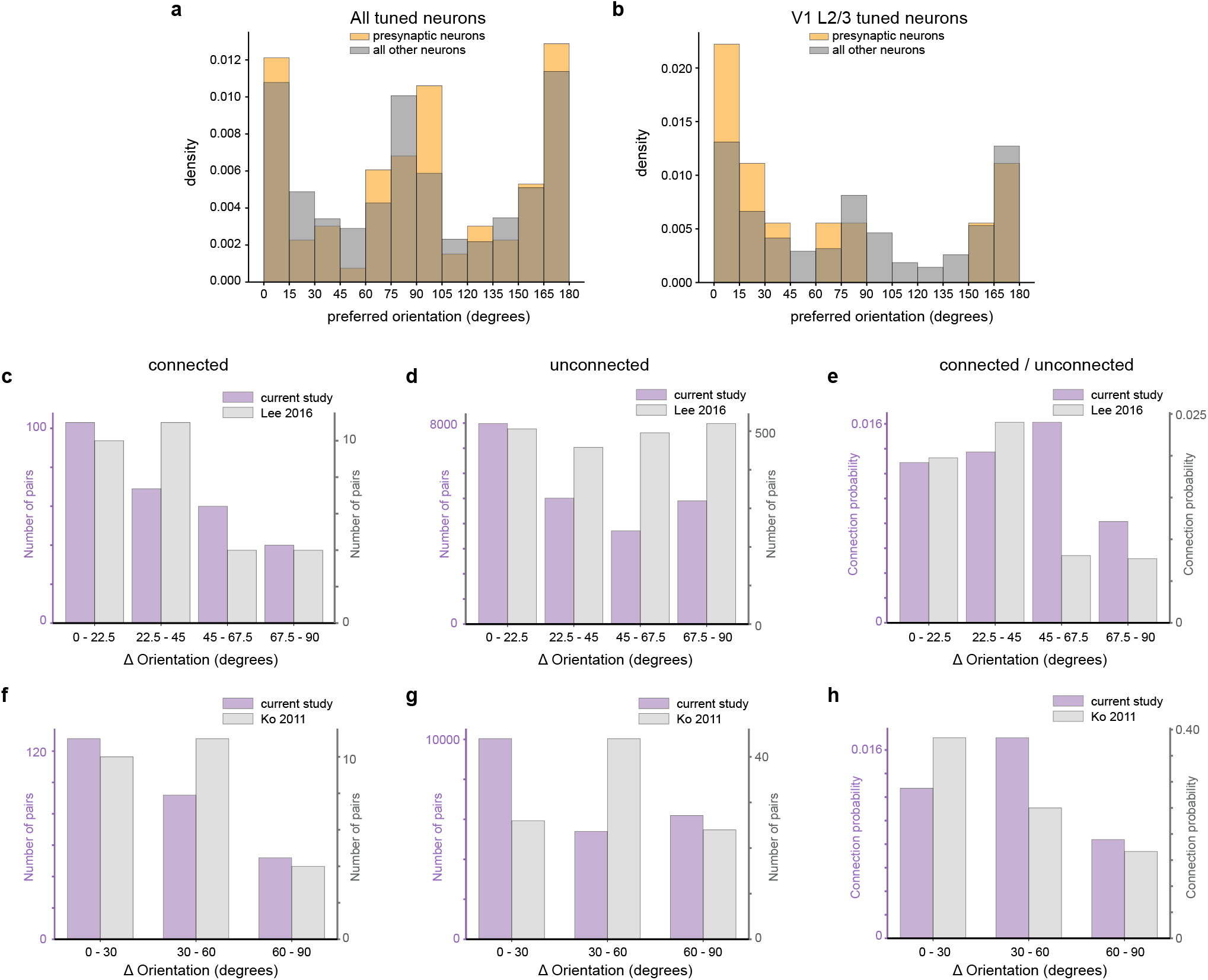
Distribution of *in silico* orientation preference and comparison to previous literature. **a**, Distribution of orientation preference of tuned neurons (gOSI > 0.25) derived from *in silico* responses to parametric stimuli (see Methods). Note the cardinal bias in orientation preference distribution, in which orientation preference for 0 and 90 degree angles is overrepresented. Gold: presynaptic neurons, Gray: all other neurons. **b**, As in **a** but for tuned neurons in V1 L2/3. Difference in preferred orientation (Δ Orientation) for neurons in V1 L2/3 for connected pairs (**c, f**), unconnected pairs (**d, g**), and the ratio of connected / unconnected (“connection probability”, **e, h**) for our study vs Lee et al. 2016 (**c-e**) and vs Ko et al. 2011 (**f-h**). The connected V1 L2/3 neurons in our study show a strong like-to-like effect, consistent with both Lee et al. 2016 and Ko et al. 2011 (**c, f**), however unlike Lee et al. 2016 and Ko et al. 2011, the unconnected neurons in our study also show a strong like-to-like effect (**d, g**) indicating that the like-to-like effect seen in connected pairs results from an orientation preference bias. This bias likely explains why we do not observe significant a like-to-like effect between V1 L2/3 neurons at axonal scale or synaptic scale in Supplemental. Fig 8, (i.e. when pairs are tested against region-matched controls).

**Supplemental Figure 10.**
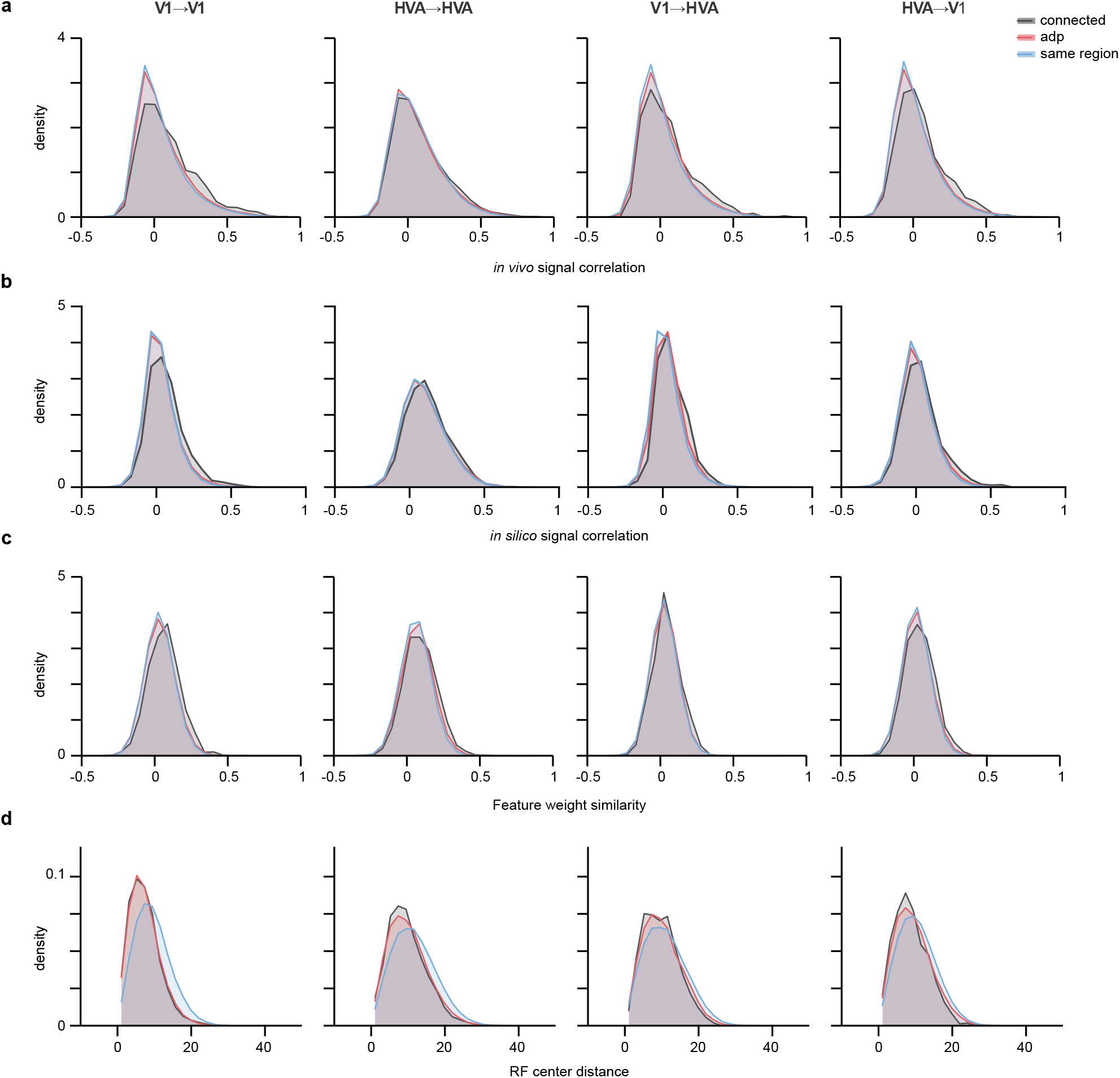
Distribution of pairwise functional measurements. Density distribution of connected pairs (black), ADP control pairs (red) and same region control pairs (blue) for *in vivo* signal correlations (**a**), *in silico* signal correlations (**b**), feature weight similarity (**c**), and RF center distance (**d**) for all projection types.

**Supplemental Figure 11.**
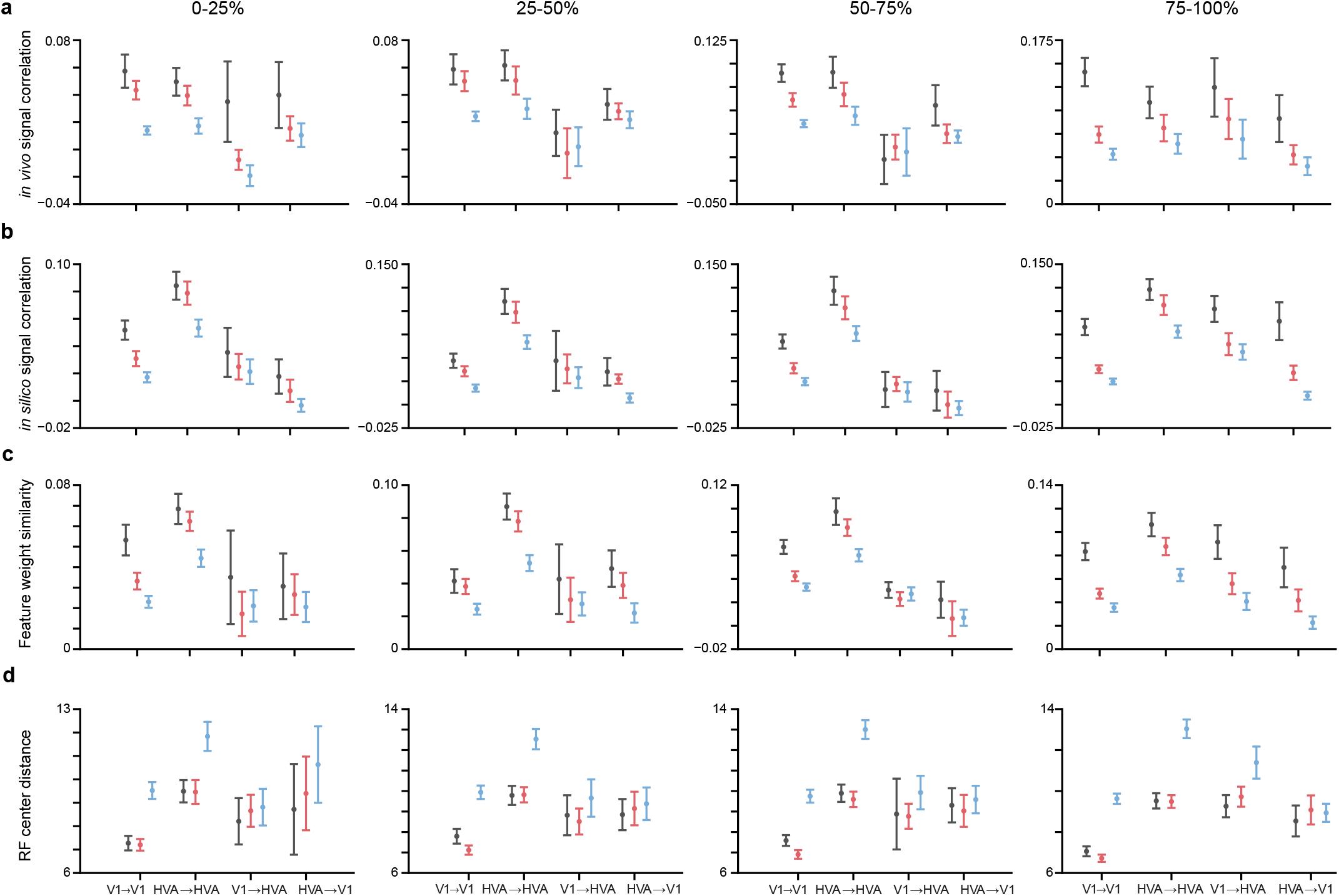
Pairwise functional measurements across varying levels of model predictive performance. Mean of *in vivo* signal correlations (**a**), *in silico* signal correlations (**b**), feature weight similarity (**c**), and RF center distance (**d**) for all projection types across 4 quantiles of model predictive performance (*CC*_*abs*_). All panels share a base filtering for visual responsiveness (*CC*_*max*_ > 0.4, 90% of neurons pass this threshold). Presynaptic neurons are filtered to *CC*_*abs*_ > 0.2 (4 did not pass this threshold).

**Supplemental Figure 12.**
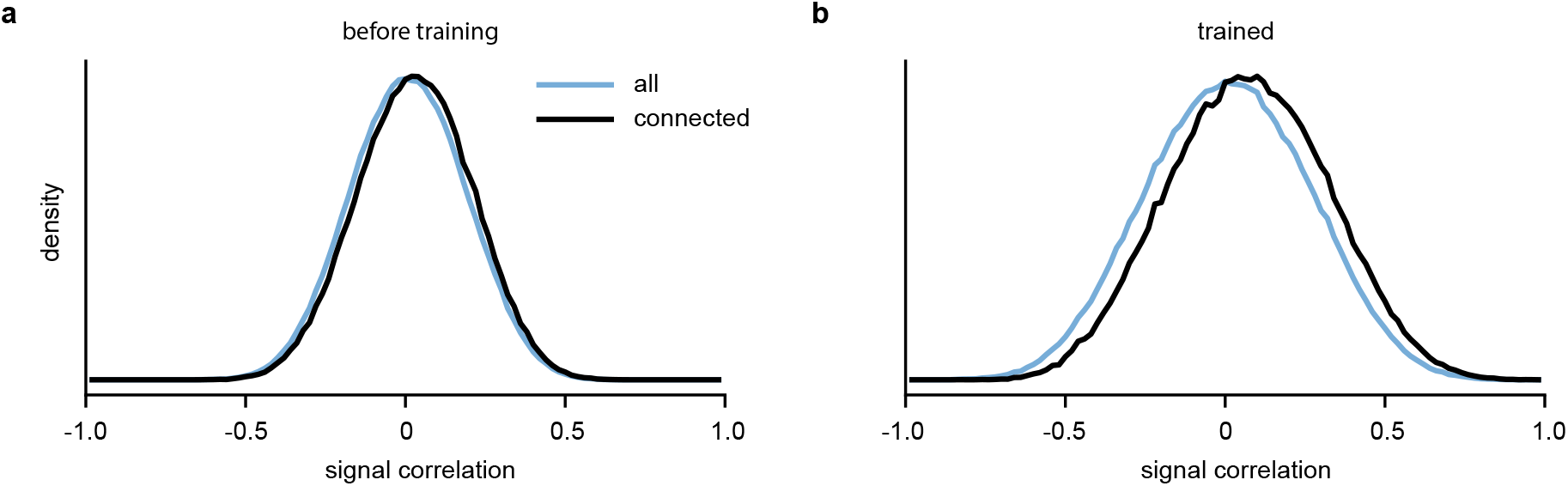
Signal correlation distributions for connected neurons vs all neurons in the RNN before and after training. **a**, Signal correlation distribution for connected neurons vs all neurons in the RNN before training. A neuron pair was classified as connected if the associated weight was in the top 35^th^ percentile of all weights. **b**, Same as **a** except after training.

**Supplemental Table 1.** Proofread presynaptic neuron nucleus ID’s, area, layer, and proofreading strategy. nucleus_id’s are from CAVE table nucleus_detection_v0.

| index | nucleus_id | area | layer | proofreading strategy |
| --- | --- | --- | --- | --- |
| 1 | 189149 | V1 | L2/3 | full cleaning and extension |
| 2 | 222998 | V1 | L2/3 | full cleaning and extension |
| 3 | 223037 | V1 | L2/3 | full cleaning and extension |
| 4 | 224565 | V1 | L2/3 | full cleaning and extension |
| 5 | 225498 | V1 | L4 | full cleaning and extension |
| 6 | 230236 | V1 | L5 | full cleaning and extension |
| 7 | 236197 | V1 | L6 | full cleaning and extension |
| 8 | 255217 | V1 | L2/3 | full cleaning and extension |
| 9 | 256443 | V1 | L2/3 | full cleaning and extension |
| 10 | 256576 | V1 | L2/3 | full cleaning and extension |
| 11 | 258307 | V1 | L2/3 | full cleaning and extension |
| 12 | 259167 | V1 | L2/3 | full cleaning and extension |
| 13 | 262773 | V1 | L4 | full cleaning and extension |
| 14 | 264870 | V1 | L4 | full cleaning and extension |
| 15 | 269247 | V1 | L6 | full cleaning and extension |
| 16 | 269380 | V1 | L6 | full cleaning and extension |
| 17 | 271518 | V1 | L6 | full cleaning and extension |
| 18 | 292676 | V1 | L2/3 | full cleaning and extension |
| 19 | 292685 | V1 | L2/3 | full cleaning and extension |
| 20 | 292713 | V1 | L2/3 | full cleaning and extension |
| 21 | 294484 | V1 | L2/3 | full cleaning and extension |
| 22 | 294545 | V1 | L2/3 | full cleaning and extension |
| 23 | 294657 | V1 | L2/3 | full cleaning and extension |
| 24 | 294776 | V1 | L2/3 | full cleaning and extension |
| 25 | 294858 | V1 | L2/3 | full cleaning and extension |
| 26 | 294897 | V1 | L2/3 | full cleaning and extension |
| 27 | 296726 | V1 | L2/3 | full cleaning and extension |
| 28 | 300763 | V1 | L5 | full cleaning and extension |
| 29 | 301095 | V1 | L5 | full cleaning and extension |
| 30 | 301189 | V1 | L5 | full cleaning and extension |
| 31 | 327859 | V1 | L2/3 | full cleaning and extension |
| 32 | 330079 | V1 | L4 | full cleaning and extension |
| 33 | 330326 | V1 | L4 | full cleaning and extension |
| 34 | 331945 | V1 | L5 | full cleaning and extension |
| 35 | 332199 | V1 | L4 | full cleaning and extension |
| 36 | 335175 | V1 | L5 | full cleaning and extension |
| 37 | 460053 | RL | L4 | full cleaning and extension |
| 38 | 460391 | RL | L5 | full cleaning and extension |
| 39 | 487512 | RL | L2/3 | full cleaning and extension |
| 40 | 489675 | RL | L2/3 | full cleaning and extension |
| 41 | 493419 | RL | L5 | full cleaning and extension |
| 42 | 493806 | RL | L5 | full cleaning and extension |
| 43 | 493885 | RL | L5 | full cleaning and extension |
| 44 | 493968 | RL | L4 | full cleaning and extension |
| 45 | 516758 | RL | L2/3 | full cleaning and extension |
| 46 | 517056 | RL | L2/3 | full cleaning and extension |
| 47 | 518848 | RL | L2/3 | full cleaning and extension |
| 48 | 518853 | AL | L2/3 | full cleaning and extension |
| 49 | 518898 | RL | L2/3 | full cleaning and extension |
| 50 | 520364 | RL | L4 | full cleaning and extension |
| 51 | 522656 | RL | L4 | full cleaning and extension |
| 52 | 524491 | RL | L5 | full cleaning and extension |
| 53 | 525405 | RL | L5 | full cleaning and extension |
| 54 | 525498 | RL | L5 | full cleaning and extension |
| 55 | 525758 | RL | L5 | full cleaning and extension |
| 56 | 553325 | RL | L2/3 | full cleaning and extension |
| 57 | 554200 | AL | L2/3 | full cleaning and extension |
| 58 | 554741 | RL | L2/3 | full cleaning and extension |
| 59 | 554833 | RL | L4 | full cleaning and extension |
| 60 | 554921 | RL | L2/3 | full cleaning and extension |
| 61 | 556823 | RL | L4 | full cleaning and extension |
| 62 | 557030 | RL | L4 | full cleaning and extension |
| 63 | 557121 | RL | L4 | full cleaning and extension |
| 64 | 558684 | RL | L5 | full cleaning and extension |
| 65 | 558709 | RL | L4 | full cleaning and extension |
| 66 | 559081 | RL | L5 | full cleaning and extension |

**Supplemental Table 1. Proofread presynaptic neuron nucleus ID's, area, layer, and proofreading strategy**
| index | nucleus_id | area | layer | proofreading strategy |
| --- | --- | --- | --- | --- |
| 67 | 559381 | RL | L5 | full cleaning and extension |
| 68 | 560109 | RL | L5 | full cleaning and extension |
| 69 | 560217 | RL | L5 | full cleaning and extension |
| 70 | 560530 | RL | L5 | full cleaning and extension |
| 71 | 560732 | RL | L5 | full cleaning and extension |
| 72 | 562808 | RL | L5 | full cleaning and extension |
| 73 | 581967 | AL | L2/3 | full cleaning and extension |
| 74 | 582056 | AL | L2/3 | full cleaning and extension |
| 75 | 582129 | AL | L2/3 | full cleaning and extension |
| 76 | 582210 | AL | L2/3 | full cleaning and extension |
| 77 | 583848 | AL | L2/3 | full cleaning and extension |
| 78 | 583961 | RL | L2/3 | full cleaning and extension |
| 79 | 585723 | RL | L4 | full cleaning and extension |
| 80 | 587426 | AL | L4 | full cleaning and extension |
| 81 | 588839 | RL | L5 | full cleaning and extension |
| 82 | 588983 | AL | L5 | full cleaning and extension |
| 83 | 610498 | AL | L2/3 | full cleaning and extension |
| 84 | 616159 | AL | L5 | full cleaning and extension |
| 85 | 516621 | RL | L2/3 | full cleaning and partial axonal extension |
| 86 | 516988 | RL | L2/3 | full cleaning and partial axonal extension |
| 87 | 517993 | RL | L2/3 | full cleaning and partial axonal extension |
| 88 | 518004 | RL | L2/3 | full cleaning and partial axonal extension |
| 89 | 518134 | RL | L2/3 | full cleaning and partial axonal extension |
| 90 | 518224 | RL | L2/3 | full cleaning and partial axonal extension |
| 91 | 518312 | RL | L2/3 | full cleaning and partial axonal extension |
| 92 | 518623 | RL | L2/3 | full cleaning and partial axonal extension |
| 93 | 518632 | RL | L2/3 | full cleaning and partial axonal extension |
| 94 | 519746 | RL | L2/3 | full cleaning and partial axonal extension |
| 95 | 520027 | RL | L4 | full cleaning and partial axonal extension |
| 96 | 520182 | RL | L2/3 | full cleaning and partial axonal extension |
| 97 | 551802 | RL | L2/3 | full cleaning and partial axonal extension |
| 98 | 553216 | RL | L2/3 | full cleaning and partial axonal extension |
| 99 | 553283 | RL | L2/3 | full cleaning and partial axonal extension |
| 100 | 553321 | RL | L2/3 | full cleaning and partial axonal extension |
| 101 | 553339 | RL | L2/3 | full cleaning and partial axonal extension |
| 102 | 553360 | RL | L2/3 | full cleaning and partial axonal extension |
| 103 | 553469 | RL | L2/3 | full cleaning and partial axonal extension |
| 104 | 553556 | RL | L2/3 | full cleaning and partial axonal extension |
| 105 | 553585 | RL | L2/3 | full cleaning and partial axonal extension |
| 106 | 553589 | RL | L2/3 | full cleaning and partial axonal extension |
| 107 | 554734 | RL | L2/3 | full cleaning and partial axonal extension |
| 108 | 554775 | RL | L2/3 | full cleaning and partial axonal extension |
| 109 | 554891 | RL | L2/3 | full cleaning and partial axonal extension |
| 110 | 554900 | RL | L2/3 | full cleaning and partial axonal extension |
| 111 | 555010 | RL | L2/3 | full cleaning and partial axonal extension |
| 112 | 580774 | AL | L2/3 | full cleaning and partial axonal extension |
| 113 | 580826 | AL | L2/3 | full cleaning and partial axonal extension |
| 114 | 580905 | AL | L2/3 | full cleaning and partial axonal extension |
| 115 | 580948 | RL | L2/3 | full cleaning and partial axonal extension |
| 116 | 580988 | AL | L2/3 | full cleaning and partial axonal extension |
| 117 | 581988 | AL | L2/3 | full cleaning and partial axonal extension |
| 118 | 581998 | AL | L2/3 | full cleaning and partial axonal extension |
| 119 | 582011 | AL | L2/3 | full cleaning and partial axonal extension |
| 120 | 582091 | AL | L2/3 | full cleaning and partial axonal extension |
| 121 | 582294 | AL | L2/3 | full cleaning and partial axonal extension |
| 122 | 582313 | RL | L2/3 | full cleaning and partial axonal extension |
| 123 | 582353 | AL | L2/3 | full cleaning and partial axonal extension |
| 124 | 582388 | RL | L2/3 | full cleaning and partial axonal extension |
| 125 | 582390 | RL | L2/3 | full cleaning and partial axonal extension |
| 126 | 582409 | RL | L2/3 | full cleaning and partial axonal extension |
| 127 | 582412 | RL | L2/3 | full cleaning and partial axonal extension |
| 128 | 582414 | RL | L2/3 | full cleaning and partial axonal extension |
| 129 | 582444 | RL | L2/3 | full cleaning and partial axonal extension |
| 130 | 582468 | RL | L2/3 | full cleaning and partial axonal extension |
| 131 | 582471 | RL | L2/3 | full cleaning and partial axonal extension |
| 132 | 583659 | AL | L2/3 | full cleaning and partial axonal extension |
| 133 | 583739 | AL | L2/3 | full cleaning and partial axonal extension |
| 134 | 583741 | AL | L2/3 | full cleaning and partial axonal extension |

**Supplemental Table 1. Proofread presynaptic neuron nucleus ID's, area, layer, and proofreading strategy**
| index | nucleus_id | area | layer | proofreading strategy |
| --- | --- | --- | --- | --- |
| 135 | 583792 | AL | L2/3 | full cleaning and partial axonal extension |
| 136 | 583891 | RL | L2/3 | full cleaning and partial axonal extension |
| 137 | 584004 | RL | L2/3 | full cleaning and partial axonal extension |
| 138 | 608166 | AL | L2/3 | full cleaning and partial axonal extension |
| 139 | 608213 | AL | L2/3 | full cleaning and partial axonal extension |
| 140 | 610396 | AL | L2/3 | full cleaning and partial axonal extension |
| 141 | 610403 | AL | L2/3 | full cleaning and partial axonal extension |
| 142 | 610434 | AL | L2/3 | full cleaning and partial axonal extension |
| 143 | 610535 | AL | L2/3 | full cleaning and partial axonal extension |
| 144 | 610607 | AL | L2/3 | full cleaning and partial axonal extension |
| 145 | 610615 | AL | L2/3 | full cleaning and partial axonal extension |
| 146 | 612143 | AL | L2/3 | full cleaning and partial axonal extension |
| 147 | 612266 | AL | L2/3 | full cleaning and partial axonal extension |
| 148 | 612352 | AL | L2/3 | full cleaning and partial axonal extension |

**Supplemental Table 2.** Pairwise comparison of the presynaptic mean in silico signal correlation between different neuron pair populations. For each comparison, a pairwise t-test was performed to test the null hypothesis that for each presynaptic neuron, the mean in silico signal correlation is the same between two postsynaptic populations. adjusted p-value is the adjusted p-value through the BH multicomparison correction procedure.

| Comparison | Projection type | Mean pairwise difference | p-value | adjusted p-value | t statistic | n |
| --- | --- | --- | --- | --- | --- | --- |
| ADP vs Same region | HVA->HVA | 0.015 | 5.30e-05 | 7.96e-05 | 4.405 | 53 |
| ADP vs Same region | HVA->V1 | 0.007 | 1.14e-02 | 1.14e-02 | 2.661 | 39 |
| ADP vs Same region | V1->HVA | 0.011 | 3.12e-03 | 3.41e-03 | 3.475 | 17 |
| ADP vs Same region | V1->V1 | 0.009 | 3.18e-05 | 5.45e-05 | 4.793 | 35 |
| Connected vs ADP | HVA->HVA | 0.026 | 3.58e-08 | 2.15e-07 | 6.460 | 53 |
| Connected vs ADP | HVA->V1 | 0.029 | 7.85e-06 | 1.57e-05 | 5.168 | 39 |
| Connected vs ADP | V1->HVA | 0.023 | 1.25e-03 | 1.50e-03 | 3.908 | 17 |
| Connected vs ADP | V1->V1 | 0.030 | 7.33e-06 | 1.57e-05 | 5.285 | 35 |
| Connected vs Same region | HVA->HVA | 0.042 | 3.37e-10 | 4.04e-09 | 7.733 | 53 |
| Connected vs Same region | HVA->V1 | 0.036 | 5.77e-06 | 1.57e-05 | 5.266 | 39 |
| Connected vs Same region | V1->HVA | 0.034 | 9.21e-05 | 1.23e-04 | 5.175 | 17 |
| Connected vs Same region | V1->V1 | 0.039 | 4.05e-07 | 1.62e-06 | 6.253 | 35 |

**Supplemental Table 3.**
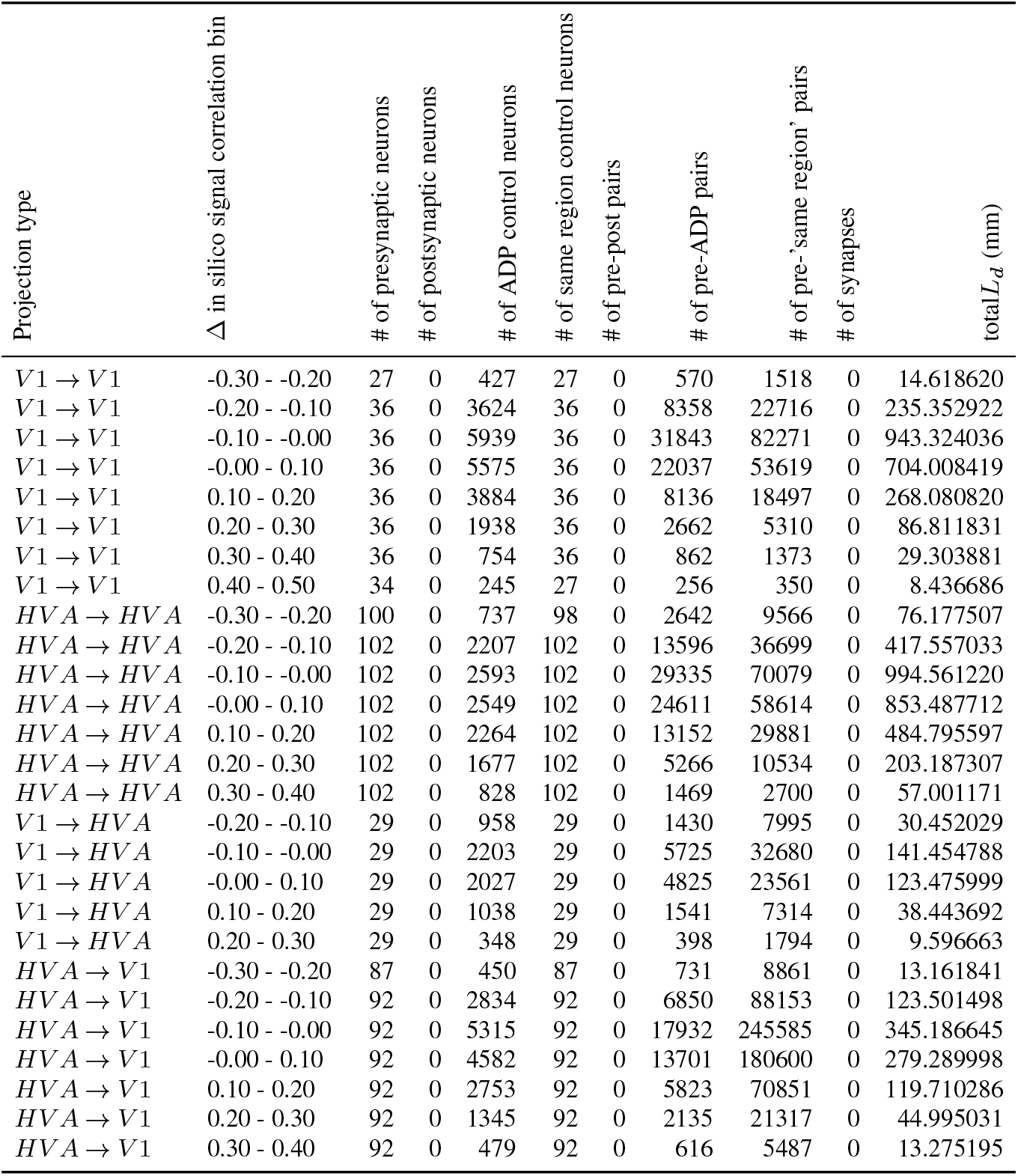
Number of neurons and neuron pairs invovled in the visualization of the correlation between in silico signal correlation and *L*_*d*_ / neuron pair (synapses excluded) in different projection types across brain areas.

**Supplemental Table 4.**
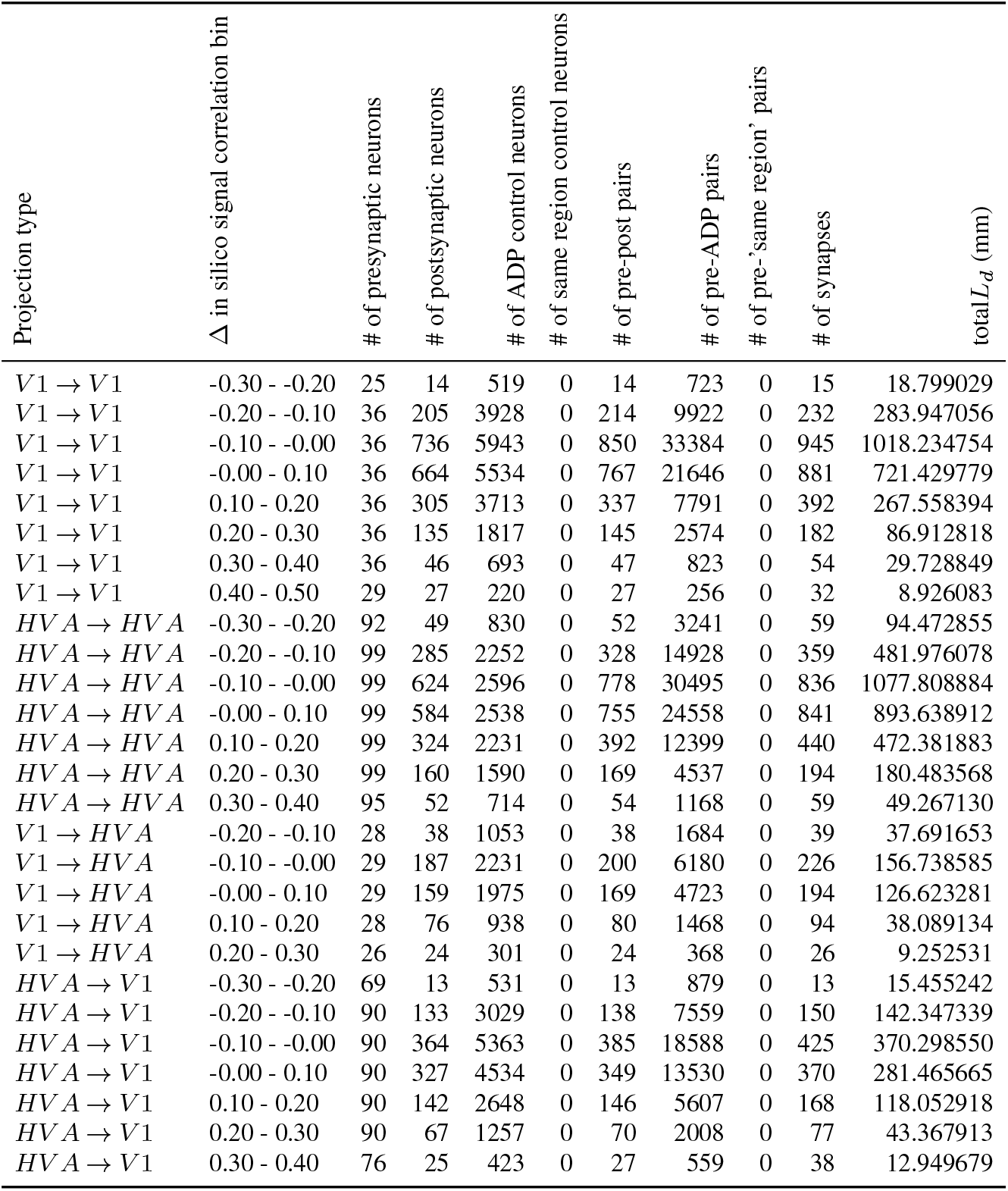
Number of neurons and neuron pairs invovled in the visualization of the correlation between in silico signal correlation and *N*_*syn*_*/mm L*_*d*_ in different projection types across brain areas.

**Supplemental Table 5.**
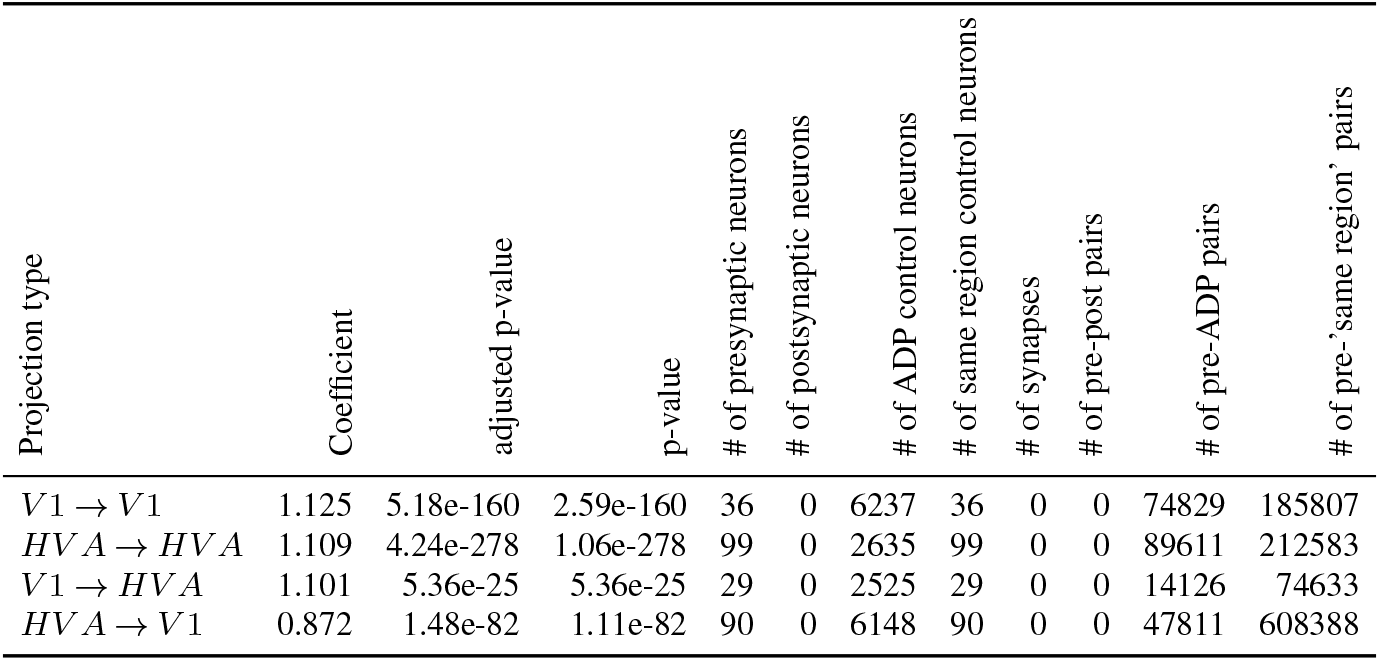
Estimated marginal means of linear trends for the effect of in silico signal correlation on *L*_*d*_ / neuron pair (synapses excluded) in different projection types across brain areas. z and p-value are the z statistics and p-value of the marginal mean linear trends estimated from the fitted GLMMs. adjusted p-value is the adjusted p value through the BH multicomparison correction procedure.

**Supplemental Table 6.** Estimated marginal means of linear trends for the effect of in silico signal correlation on *N*_*syn*_*/mm L*_*d*_ in different projection types across brain areas. z and p-value are the z statistics and p-value of the marginal mean linear trends estimated from the fitted GLMMs. adjusted p-value is the adjusted p value through the BH multicomparison correction procedure.

| Projection type | Coefficient | adjusted p-value | p-value | # of presynaptic neurons | # of postsynaptic neurons | # of ADP control neurons | # of same region control neurons | # of synapses | # of pre-post pairs | # of pre-ADP pairs | # of pre-'same region' pairs |
| --- | --- | --- | --- | --- | --- | --- | --- | --- | --- | --- | --- |
| $V1 \rightarrow V1$ | 2.297 | 9.17e-50 | 2.29e-50 | 36 | 1719 | 6237 | 0 | 2744 | 2411 | 77240 | 0 |
| $HVA \rightarrow HVA$ | 1.043 | 3.84e-12 | 2.88e-12 | 99 | 1396 | 2635 | 0 | 2803 | 2543 | 92154 | 0 |
| $V1 \rightarrow HVA$ | 1.985 | 8.57e-07 | 8.57e-07 | 29 | 448 | 2525 | 0 | 584 | 515 | 14641 | 0 |
| $HVA \rightarrow V1$ | 1.603 | 2.59e-12 | 1.29e-12 | 90 | 974 | 6148 | 0 | 1255 | 1139 | 48950 | 0 |

**Supplemental Table 7.**
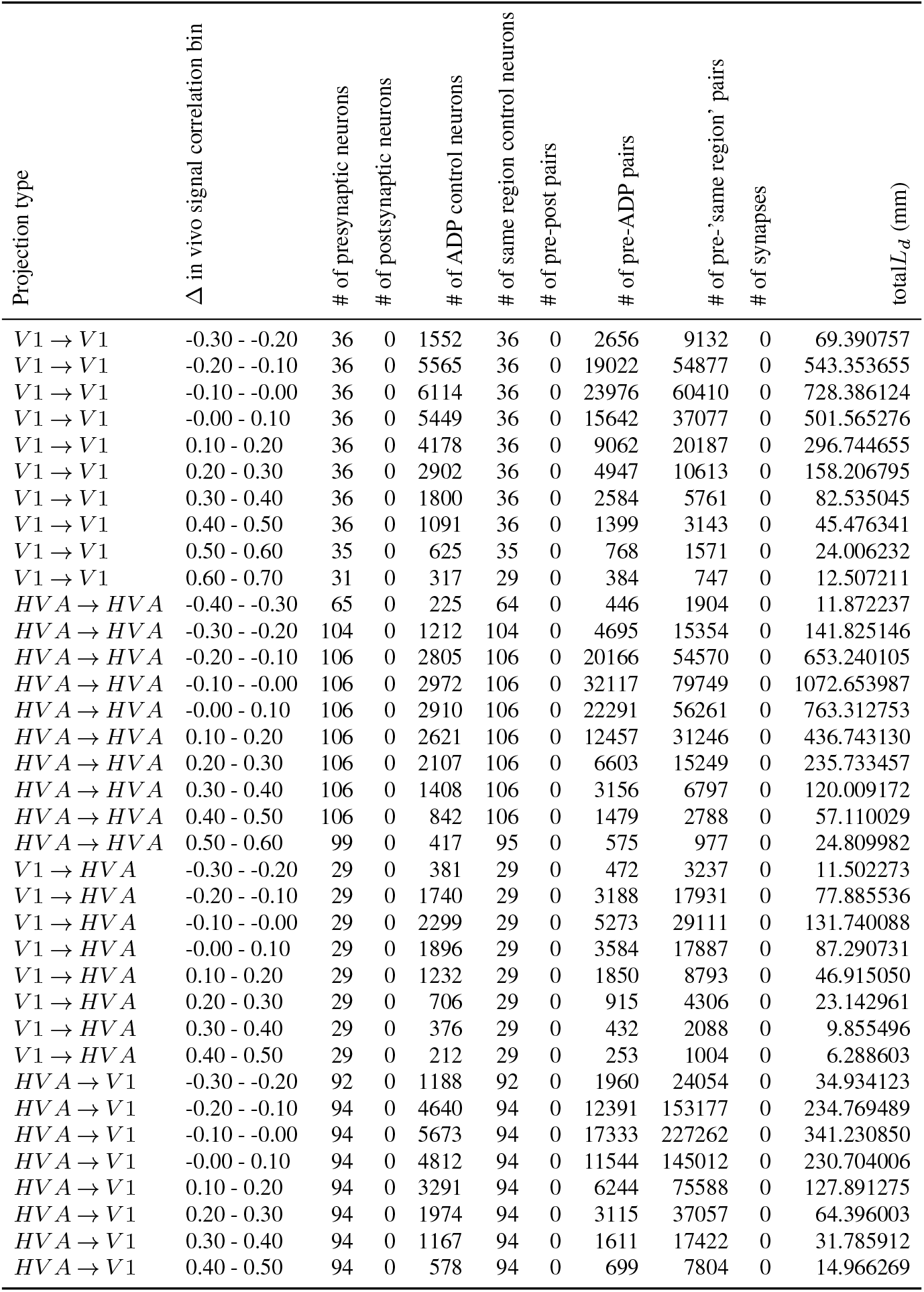
Number of neurons and neuron pairs invovled in the visualization of the correlation between in vivo signal correlation and *L*_*d*_ / neuron pair (synapses excluded) in different projection types across brain areas.

**Supplemental Table 8.**
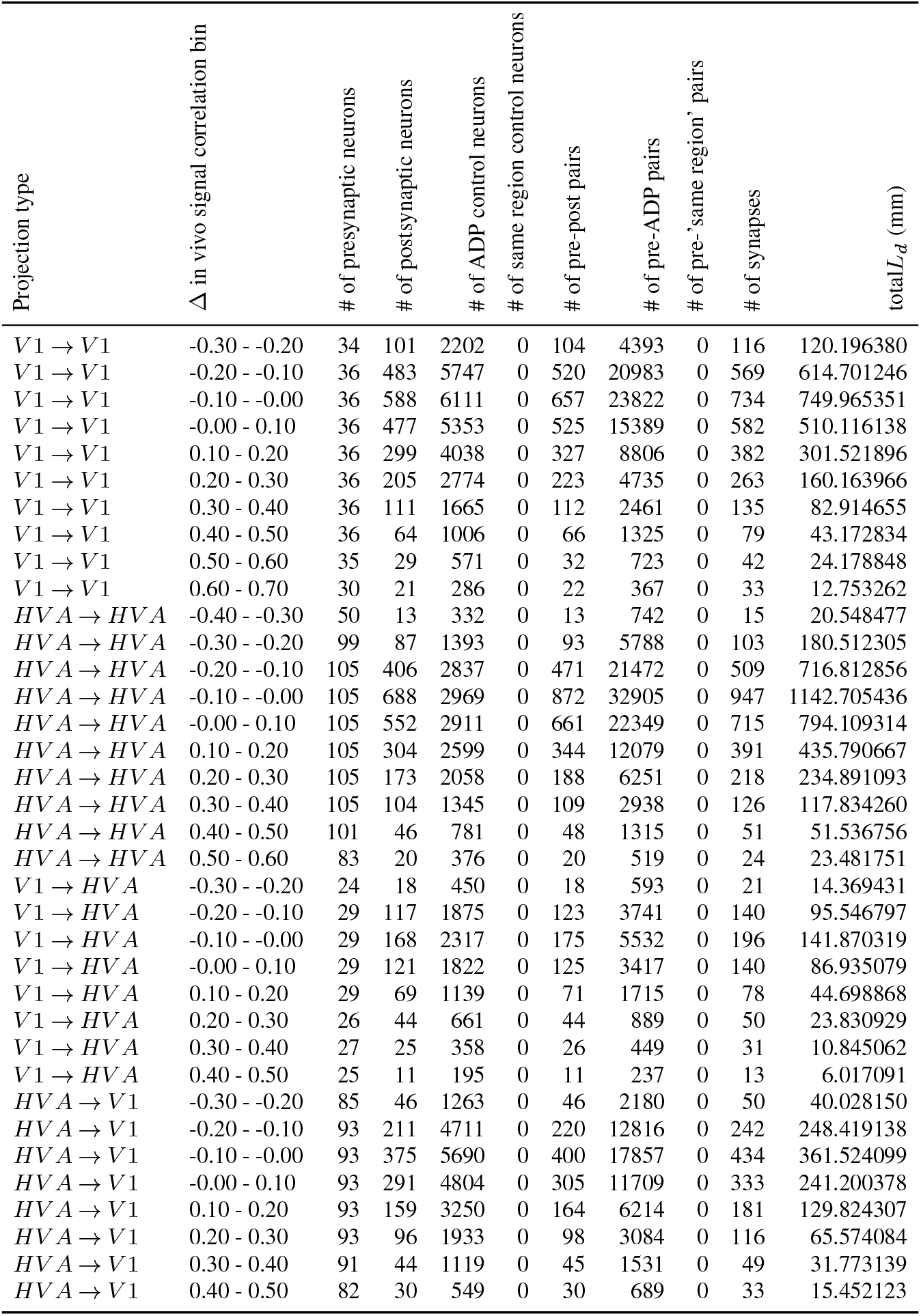
Number of neurons and neuron pairs invovled in the visualization of the correlation between in vivo signal correlation and *N*_*syn*_*/mm L*_*d*_ in different projection types across brain areas.

**Supplemental Table 9.** Estimated marginal means of linear trends for the effect of in vivo signal correlation on *L*_*d*_ / neuron pair (synapses excluded) in different projection types across brain areas. z and p-value are the z statistics and p-value of the marginal mean linear trends estimated from the fitted GLMMs. adjusted p-value is the adjusted p value through the BH multicomparison correction procedure.

| Projection type | Coefficient | adjusted p-value | p-value | # of presynaptic neurons | # of postsynaptic neurons | # of ADP control neurons | # of same region control neurons | # of synapses | # of pre-post pairs | # of pre-ADP pairs | # of pre-'same region' pairs |
| --- | --- | --- | --- | --- | --- | --- | --- | --- | --- | --- | --- |
| $V1 \rightarrow V1$ | 0.779 | 1.96e-210 | 4.89e-211 | 36 | 0 | 6807 | 36 | 0 | 0 | 80703 | 204150 |
| $HVA \rightarrow HVA$ | 0.698 | 3.51e-198 | 1.75e-198 | 105 | 0 | 3010 | 105 | 0 | 0 | 103839 | 262309 |
| $V1 \rightarrow HVA$ | 0.624 | 1.60e-23 | 1.60e-23 | 29 | 0 | 2887 | 29 | 0 | 0 | 16164 | 85137 |
| $HVA \rightarrow V1$ | 0.428 | 5.69e-43 | 4.27e-43 | 93 | 0 | 6720 | 93 | 0 | 0 | 55313 | 685610 |

**Supplemental Table 10.** Estimated marginal means of linear trends for the effect of in vivo signal correlation on *N*_*syn*_*/mm L*_*d*_ in different projection types across brain areas. z and p-value are the z statistics and p-value of the marginal mean linear trends estimated from the fitted GLMMs. adjusted p-value is the adjusted p value through the BH multicomparison correction procedure.

| Projection type | Coefficient | adjusted p-value | p-value | # of presynaptic neurons | # of postsynaptic neurons | # of ADP control neurons | # of same region control neurons | # of synapses | # of pre-post pairs | # of pre-ADP pairs | # of pre-'same region' pairs |
| --- | --- | --- | --- | --- | --- | --- | --- | --- | --- | --- | --- |
| $V1 \rightarrow V1$ | 1.126 | 3.66e-30 | 9.16e-31 | 36 | 1865 | 6807 | 0 | 2947 | 2600 | 83303 | 0 |
| $HVA \rightarrow HVA$ | 0.667 | 1.96e-09 | 9.79e-10 | 105 | 1566 | 3010 | 0 | 3114 | 2832 | 106671 | 0 |
| $V1 \rightarrow HVA$ | 0.857 | 1.63e-04 | 1.63e-04 | 29 | 522 | 2887 | 0 | 679 | 602 | 16766 | 0 |
| $HVA \rightarrow V1$ | 0.861 | 9.53e-08 | 7.15e-08 | 93 | 1116 | 6720 | 0 | 1445 | 1315 | 56628 | 0 |

**Supplemental Table 11.**
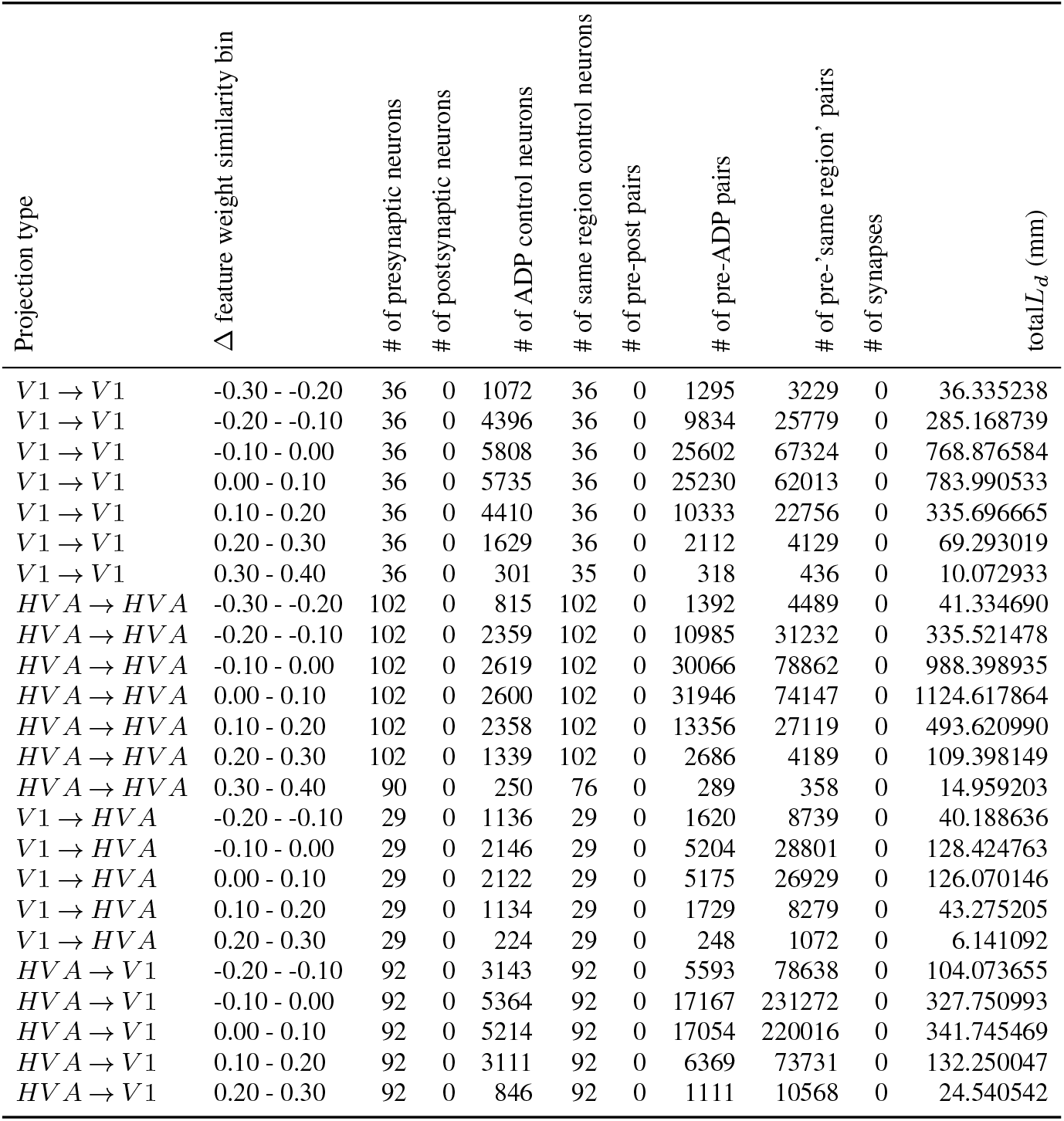
Number of neurons and neuron pairs invovled in the visualization of the correlation between feature weight similarity and *L*_*d*_ / neuron pair (synapses excluded) in different projection types across brain areas.

**Supplemental Table 12.** Number of neurons and neuron pairs invovled in the visualization of the correlation between feature weight similarity and *N*_*syn*_*/mm L*_*d*_ in different projection types across brain areas.

| Projection type | $\Delta$ feature weight similarity bin | # of presynaptic neurons | # of postsynaptic neurons | # of ADP control neurons | # of same region control neurons | # of pre-post pairs | # of pre-ADP pairs | # of pre-'same region' pairs | # of synapses | total $L_d$ (mm) |
| --- | --- | --- | --- | --- | --- | --- | --- | --- | --- | --- |
| V1 → V1 | -0.30 - -0.20 | 36 | 36 | 1182 | 0 | 36 | 1509 | 0 | 38 | 43.620452 |
| V1 → V1 | -0.20 - -0.10 | 36 | 230 | 4537 | 0 | 237 | 10895 | 0 | 258 | 322.934569 |
| V1 → V1 | -0.10 - 0.00 | 36 | 630 | 5812 | 0 | 708 | 27056 | 0 | 794 | 833.029064 |
| V1 → V1 | 0.00 - 0.10 | 36 | 729 | 5720 | 0 | 852 | 25425 | 0 | 969 | 819.256558 |
| V1 → V1 | 0.10 - 0.20 | 36 | 388 | 4292 | 0 | 438 | 9940 | 0 | 509 | 337.394190 |
| V1 → V1 | 0.20 - 0.30 | 36 | 102 | 1496 | 0 | 112 | 2007 | 0 | 139 | 69.123548 |
| V1 → V1 | 0.30 - 0.40 | 33 | 24 | 259 | 0 | 24 | 294 | 0 | 31 | 10.411849 |
| HVA → HVA | -0.30 - -0.20 | 99 | 33 | 968 | 0 | 35 | 1852 | 0 | 37 | 55.775849 |
| HVA → HVA | -0.20 - -0.10 | 99 | 254 | 2421 | 0 | 276 | 12617 | 0 | 301 | 398.118425 |
| HVA → HVA | -0.10 - 0.00 | 99 | 663 | 2619 | 0 | 807 | 32094 | 0 | 876 | 1097.132235 |
| HVA → HVA | 0.00 - 0.10 | 99 | 649 | 2594 | 0 | 847 | 31140 | 0 | 926 | 1149.553761 |
| HVA → HVA | 0.10 - 0.20 | 99 | 386 | 2318 | 0 | 450 | 11911 | 0 | 514 | 461.647156 |
| HVA → HVA | 0.20 - 0.30 | 98 | 104 | 1189 | 0 | 109 | 2216 | 0 | 128 | 98.577172 |
| HVA → HVA | 0.30 - 0.40 | 71 | 15 | 183 | 0 | 15 | 213 | 0 | 16 | 11.813672 |
| V1 → HVA | -0.20 - -0.10 | 28 | 45 | 1187 | 0 | 47 | 1799 | 0 | 50 | 45.872386 |
| V1 → HVA | -0.10 - 0.00 | 29 | 174 | 2162 | 0 | 179 | 5474 | 0 | 196 | 139.731979 |
| V1 → HVA | 0.00 - 0.10 | 29 | 196 | 2103 | 0 | 204 | 5296 | 0 | 242 | 132.939401 |
| V1 → HVA | 0.10 - 0.20 | 29 | 67 | 1102 | 0 | 70 | 1684 | 0 | 80 | 44.564126 |
| V1 → HVA | 0.20 - 0.30 | 27 | 13 | 196 | 0 | 13 | 227 | 0 | 14 | 5.948031 |
| HVA → V1 | -0.20 - -0.10 | 90 | 106 | 3354 | 0 | 109 | 6261 | 0 | 121 | 120.205383 |
| HVA → V1 | -0.10 - 0.00 | 90 | 370 | 5397 | 0 | 389 | 18035 | 0 | 429 | 355.383651 |
| HVA → V1 | 0.00 - 0.10 | 90 | 365 | 5192 | 0 | 396 | 16927 | 0 | 425 | 348.693858 |
| HVA → V1 | 0.10 - 0.20 | 90 | 187 | 2962 | 0 | 195 | 5992 | 0 | 218 | 128.408223 |
| HVA → V1 | 0.20 - 0.30 | 84 | 34 | 709 | 0 | 36 | 941 | 0 | 45 | 20.898991 |

**Supplemental Table 13.**
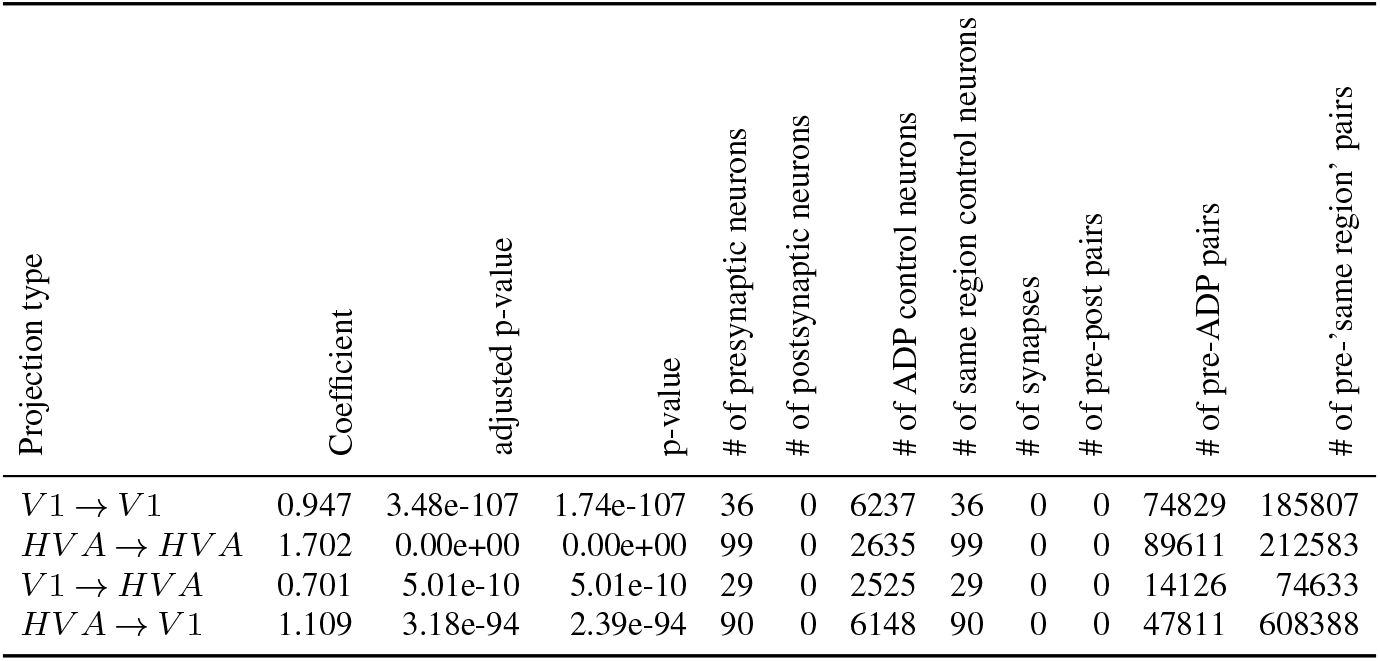
Estimated marginal means of linear trends for the effect of feature weight similarity on *L*_*d*_ / neuron pair (synapses excluded) in different projection types across brain areas. z and p-value are the z statistics and p-value of the marginal mean linear trends estimated from the fitted GLMMs. adjusted p-value is the adjusted p value through the BH multicomparison correction procedure.

**Supplemental Table 14.** Estimated marginal means of linear trends for the effect of feature weight similarity on *N*_*syn*_*/mm L*_*d*_ in different projection types across brain areas. z and p-value are the z statistics and p-value of the marginal mean linear trends estimated from the fitted GLMMs. adjusted p-value is the adjusted p value through the BH multicomparison correction procedure.

| Projection type | Coefficient | adjusted p-value | p-value | # of presynaptic neurons | # of postsynaptic neurons | # of ADP control neurons | # of same region control neurons | # of synapses | # of pre-post pairs | # of pre-ADP pairs | # of pre-'same region' pairs |
| --- | --- | --- | --- | --- | --- | --- | --- | --- | --- | --- | --- |
| V1 → V1 | 2.216 | 2.45e-35 | 6.12e-36 | 36 | 1719 | 6237 | 0 | 2744 | 2411 | 77240 | 0 |
| HVA → HVA | 1.398 | 1.13e-13 | 5.64e-14 | 99 | 1396 | 2635 | 0 | 2803 | 2543 | 92154 | 0 |
| V1 → HVA | 1.754 | 9.51e-05 | 9.51e-05 | 29 | 448 | 2525 | 0 | 584 | 515 | 14641 | 0 |
| HVA → V1 | 1.948 | 1.49e-11 | 1.11e-11 | 90 | 974 | 6148 | 0 | 1255 | 1139 | 48950 | 0 |

**Supplemental Table 15.**
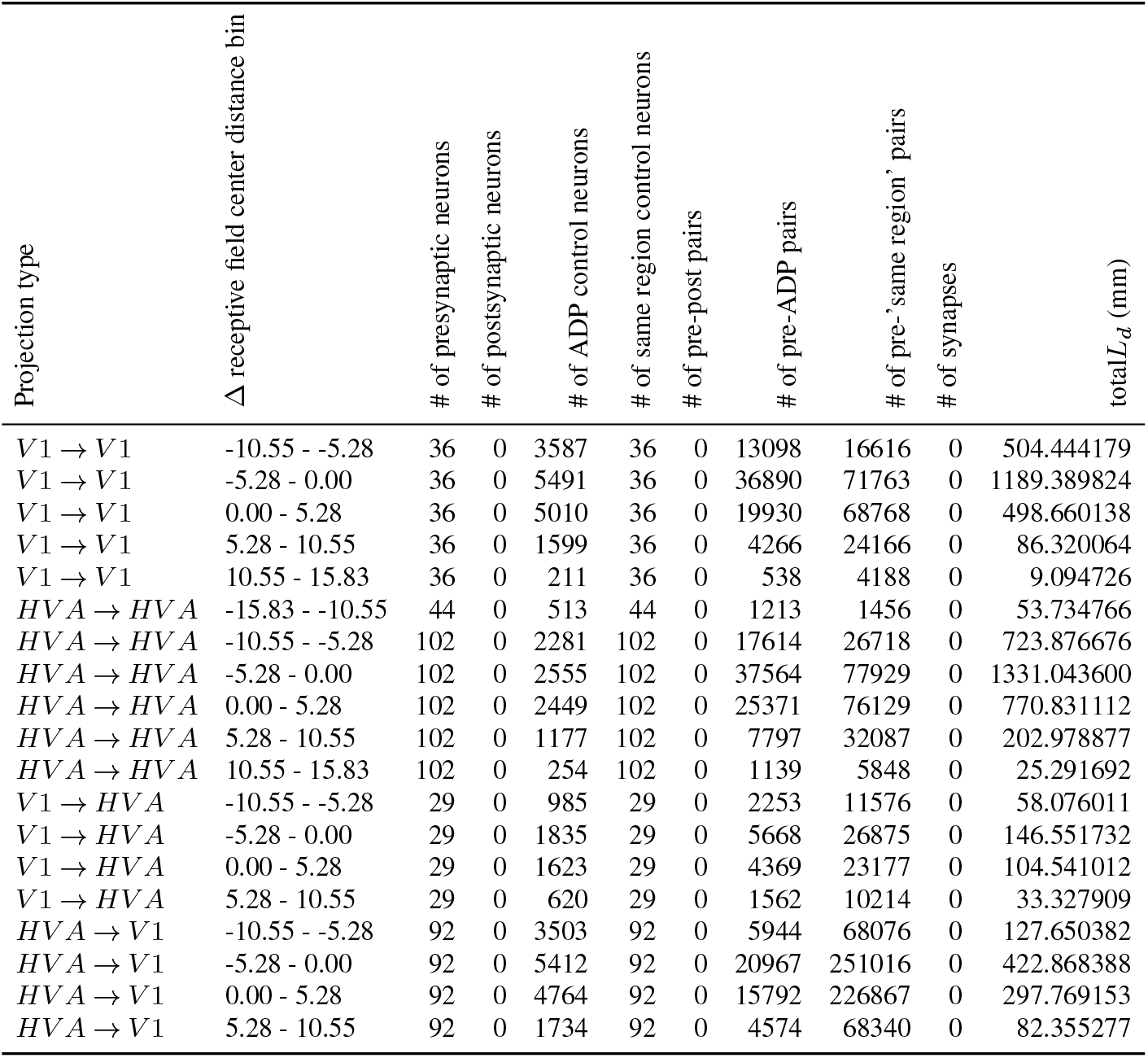
Number of neurons and neuron pairs invovled in the visualization of the correlation between receptive field center distance and *L*_*d*_ / neuron pair (synapses excluded) in different projection types across brain areas.

**Supplemental Table 16.**
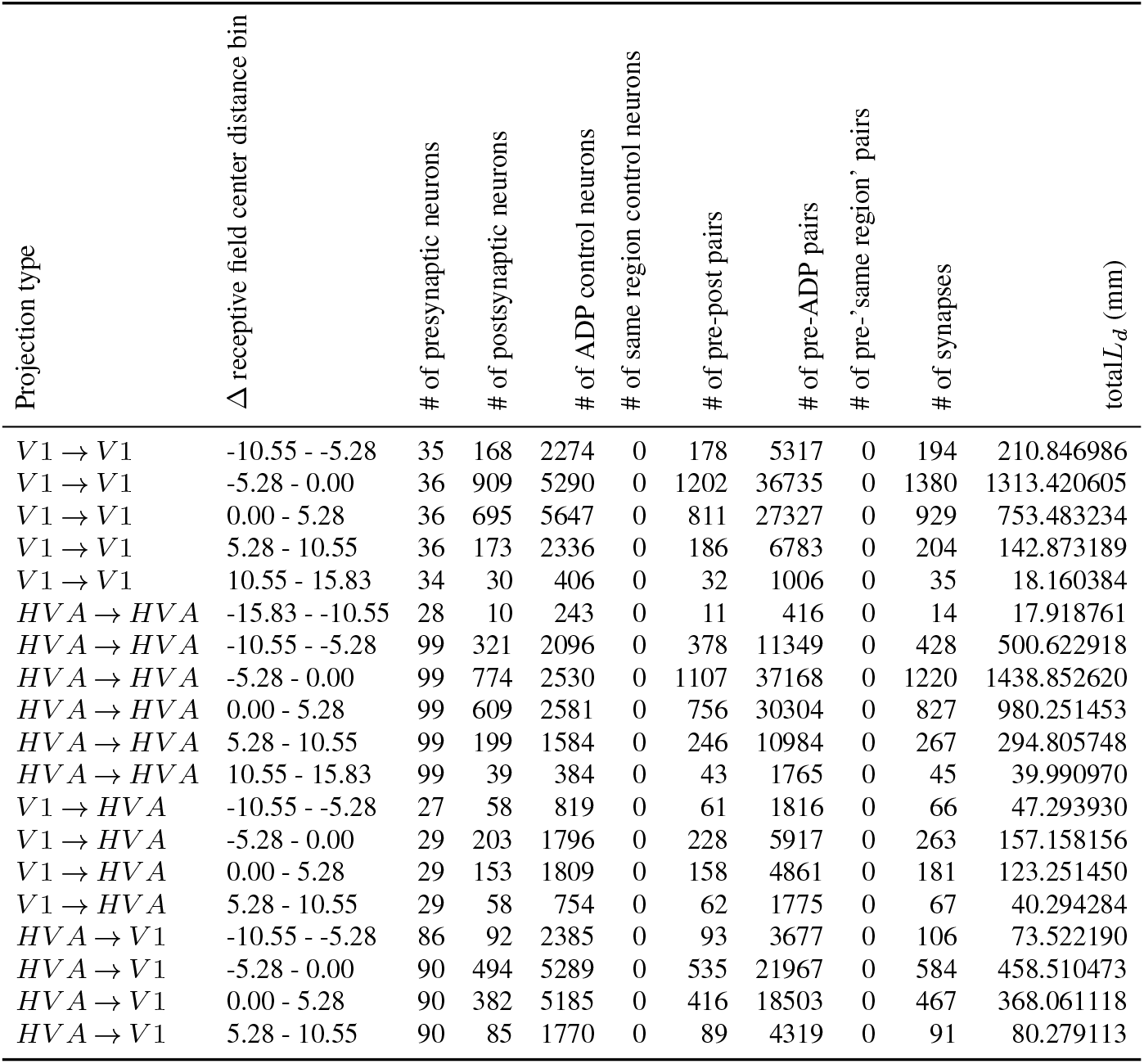
Number of neurons and neuron pairs invovled in the visualization of the correlation between receptive field center distance and *N*_*syn*_*/mm L*_*d*_ in different projection types across brain areas.

**Supplemental Table 17.** Estimated marginal means of linear trends for the effect of receptive field center distance on *L*_*d*_ / neuron pair (synapses excluded) in different projection types across brain areas. z and p-value are the z statistics and p-value of the marginal mean linear trends estimated from the fitted GLMMs. adjusted p-value is the adjusted p value through the BH multicomparison correction procedure.

| Projection type | Coefficient | adjusted p-value | p-value | # of presynaptic neurons | # of postsynaptic neurons | # of ADP control neurons | # of same region control neurons | # of synapses | # of pre-post pairs | # of pre-ADP pairs | # of pre-'same region' pairs |
| --- | --- | --- | --- | --- | --- | --- | --- | --- | --- | --- | --- |
| $V1 \rightarrow V1$ | -0.127 | 0.00e+00 | 0.00e+00 | 36 | 0 | 6237 | 36 | 0 | 0 | 74829 | 185807 |
| $HVA \rightarrow HVA$ | -0.080 | 0.00e+00 | 0.00e+00 | 99 | 0 | 2635 | 99 | 0 | 0 | 89611 | 212583 |
| $V1 \rightarrow HVA$ | -0.022 | 1.01e-27 | 1.01e-27 | 29 | 0 | 2525 | 29 | 0 | 0 | 14126 | 74633 |
| $HVA \rightarrow V1$ | -0.027 | 8.19e-119 | 6.14e-119 | 90 | 0 | 6148 | 90 | 0 | 0 | 47811 | 608388 |

**Supplemental Table 18.**
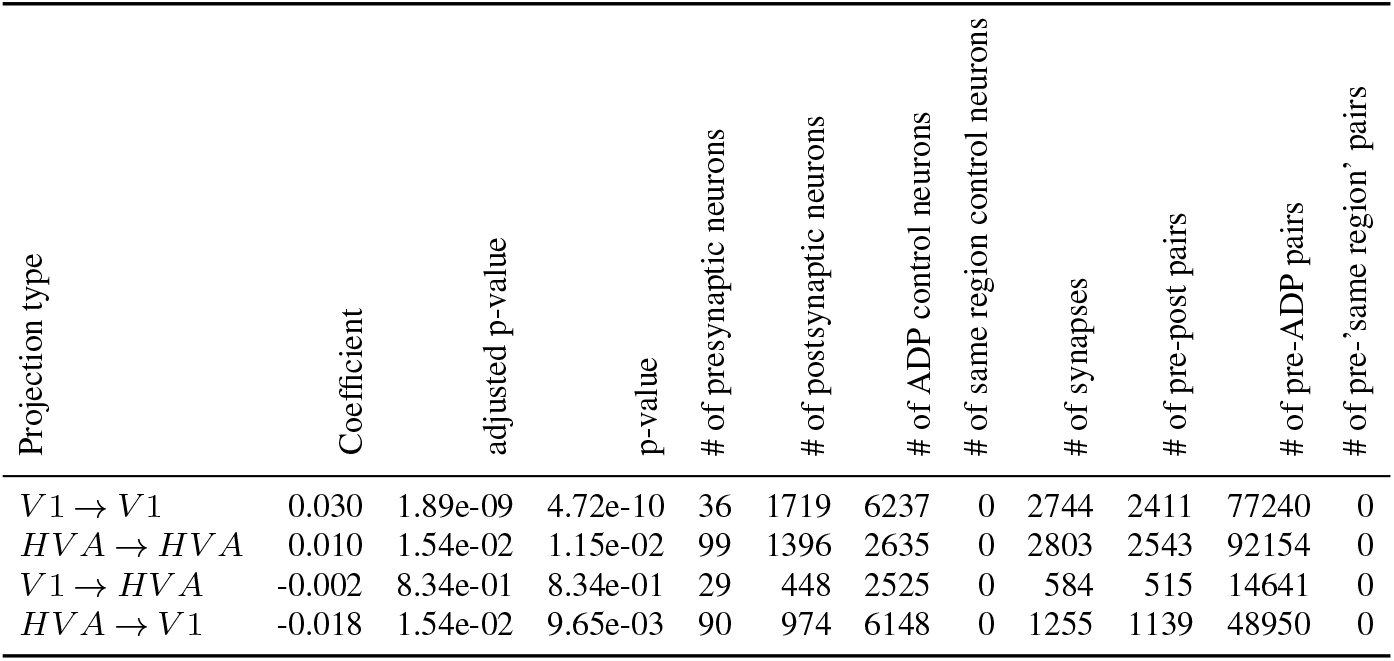
Estimated marginal means of linear trends for the effect of receptive field center distance on *N*_*syn*_*/mm L*_*d*_ in different projection types across brain areas. z and p-value are the z statistics and p-value of the marginal mean linear trends estimated from the fitted GLMMs. adjusted p-value is the adjusted p value through the BH multicomparison correction procedure.

**Supplemental Table 19.**
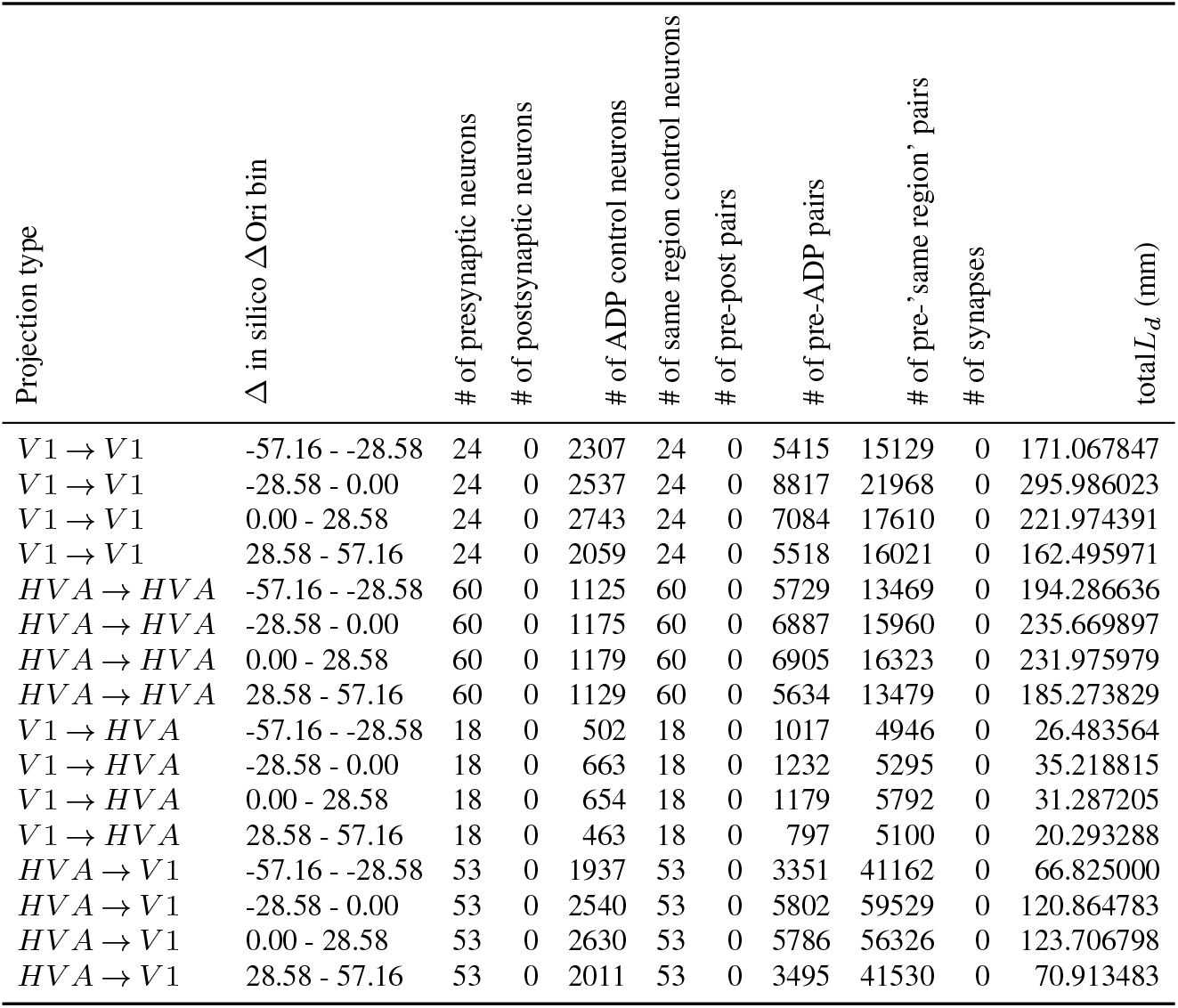
Number of neurons and neuron pairs invovled in the visualization of the correlation between in silico ΔOri and *L*_*d*_ / neuron pair (synapses excluded) in different projection types across brain areas.

**Supplemental Table 20.** Number of neurons and neuron pairs invovled in the visualization of the correlation between in silico ΔOri and *N*_*syn*_*/mm L*_*d*_ in different projection types across brain areas.

| Projection type | $\Delta$ in silico $\Delta Ori$ bin | # of presynaptic neurons | # of postsynaptic neurons | # of ADP control neurons | # of same region control neurons | # of pre-post pairs | # of pre-ADP pairs | # of pre-'same region' pairs | # of synapses | total $L_d$ (mm) |
| --- | --- | --- | --- | --- | --- | --- | --- | --- | --- | --- |
| V1 → V1 | -57.16 - -28.58 | 24 | 184 | 2256 | 0 | 203 | 5252 | 0 | 254 | 171.406452 |
| V1 → V1 | -28.58 - 0.00 | 24 | 302 | 2524 | 0 | 336 | 9435 | 0 | 394 | 329.222605 |
| V1 → V1 | 0.00 - 28.58 | 24 | 217 | 2816 | 0 | 241 | 7546 | 0 | 268 | 245.866967 |
| V1 → V1 | 28.58 - 57.16 | 24 | 144 | 1979 | 0 | 157 | 5538 | 0 | 180 | 166.321087 |
| HVA → HVA | -57.16 - -28.58 | 52 | 134 | 1113 | 0 | 155 | 5274 | 0 | 169 | 190.852637 |
| HVA → HVA | -28.58 - 0.00 | 52 | 169 | 1165 | 0 | 180 | 6698 | 0 | 193 | 241.124324 |
| HVA → HVA | 0.00 - 28.58 | 52 | 157 | 1179 | 0 | 173 | 6701 | 0 | 181 | 240.263087 |
| HVA → HVA | 28.58 - 57.16 | 52 | 129 | 1122 | 0 | 150 | 5299 | 0 | 165 | 182.145706 |
| V1 → HVA | -57.16 - -28.58 | 16 | 39 | 479 | 0 | 40 | 854 | 0 | 48 | 23.297975 |
| V1 → HVA | -28.58 - 0.00 | 16 | 35 | 655 | 0 | 36 | 1373 | 0 | 45 | 40.063947 |
| V1 → HVA | 0.00 - 28.58 | 16 | 50 | 678 | 0 | 51 | 1216 | 0 | 54 | 34.018144 |
| V1 → HVA | 28.58 - 57.16 | 16 | 28 | 481 | 0 | 30 | 925 | 0 | 33 | 24.334127 |
| HVA → V1 | -57.16 - -28.58 | 46 | 73 | 1831 | 0 | 75 | 3198 | 0 | 81 | 66.090308 |
| HVA → V1 | -28.58 - 0.00 | 47 | 138 | 2502 | 0 | 140 | 5974 | 0 | 159 | 128.419254 |
| HVA → V1 | 0.00 - 28.58 | 47 | 140 | 2641 | 0 | 149 | 5964 | 0 | 158 | 129.941363 |
| HVA → V1 | 28.58 - 57.16 | 47 | 63 | 1867 | 0 | 64 | 3187 | 0 | 69 | 67.486736 |

**Supplemental Table 21.** Estimated marginal means of linear trends for the effect of in silico ΔOri on *L*_*d*_ / neuron pair (synapses excluded) in different projection types across brain areas. z and p-value are the z statistics and p-value of the marginal mean linear trends estimated from the fitted GLMMs. adjusted p-value is the adjusted p value through the BH multicomparison correction procedure.

| Projection type | Coefficient | adjusted p-value | p-value | # of presynaptic neurons | # of postsynaptic neurons | # of ADP control neurons | # of same region control neurons | # of synapses | # of pre-post pairs | # of pre-ADP pairs | # of pre-'same region' pairs |
| --- | --- | --- | --- | --- | --- | --- | --- | --- | --- | --- | --- |
| $V1 \rightarrow V1$ | -0.001 | 1.22e-05 | 6.11e-06 | 24 | 0 | 3456 | 24 | 0 | 0 | 26834 | 70728 |
| $HVA \rightarrow HVA$ | -0.001 | 4.70e-02 | 4.70e-02 | 52 | 0 | 1222 | 52 | 0 | 0 | 23314 | 49735 |
| $V1 \rightarrow HVA$ | -0.004 | 4.65e-08 | 1.16e-08 | 16 | 0 | 1123 | 16 | 0 | 0 | 4211 | 18313 |
| $HVA \rightarrow V1$ | 0.001 | 3.02e-02 | 2.27e-02 | 47 | 0 | 3392 | 47 | 0 | 0 | 17907 | 174465 |

**Supplemental Table 22.** Estimated marginal means of linear trends for the effect of in silico ΔOri on *N*_*syn*_*/mm L*_*d*_ in different projection types across brain areas. z and p-value are the z statistics and p-value of the marginal mean linear trends estimated from the fitted GLMMs. adjusted p-value is the adjusted p value through the BH multicomparison correction procedure.

| Projection type | Coefficient | adjusted p-value | p-value | # of presynaptic neurons | # of postsynaptic neurons | # of ADP control neurons | # of same region control neurons | # of synapses | # of pre-post pairs | # of pre-ADP pairs | # of pre-'same region' pairs |
| --- | --- | --- | --- | --- | --- | --- | --- | --- | --- | --- | --- |
| V1 → V1 | -0.004 | 1.12e-03 | 2.79e-04 | 24 | 739 | 3456 | 0 | 1096 | 937 | 27771 | 0 |
| HVA → HVA | 0.000 | 9.36e-01 | 9.36e-01 | 52 | 452 | 1222 | 0 | 708 | 658 | 23972 | 0 |
| V1 → HVA | -0.004 | 2.33e-01 | 1.75e-01 | 16 | 147 | 1123 | 0 | 180 | 157 | 4368 | 0 |
| HVA → V1 | -0.002 | 2.33e-01 | 1.68e-01 | 47 | 392 | 3392 | 0 | 467 | 428 | 18335 | 0 |

**Supplemental Table 23.** Estimated marginal means of linear trends for the effect of in vivo signal correlation on *L*_*d*_ / neuron pair (synapses excluded) in different projection types across brain areas and layers. z and p-value are the z statistics and p-value of the marginal mean linear trends estimated from the fitted GLMMs. adjusted p-value is the adjusted p value through the BH multicomparison correction procedure.

| Projection type | Coefficient | adjusted p-value | p-value | # of presynaptic neurons | # of postsynaptic neurons | # of ADP control neurons | # of same region control neurons | # of synapses | # of pre-post pairs | # of pre-ADP pairs | # of pre-'same region' pairs |
| --- | --- | --- | --- | --- | --- | --- | --- | --- | --- | --- | --- |
| V1L2/3 → V1L2/3 | 1.074 | 2.49e-109 | 2.08e-110 | 20 | 0 | 2670 | 20 | 0 | 0 | 20511 | 44349 |
| V1L2/3 → V1L4 | 0.341 | 1.52e-07 | 7.59e-08 | 19 | 0 | 2090 | 19 | 0 | 0 | 13791 | 32623 |
| V1L2/3 → V1L5 | 0.580 | 6.10e-18 | 1.27e-18 | 20 | 0 | 1185 | 20 | 0 | 0 | 11845 | 14652 |
| V1L2/3 → HV AL2/3 | 1.210 | 1.21e-25 | 2.02e-26 | 15 | 0 | 1169 | 15 | 0 | 0 | 4610 | 18075 |
| V1L2/3 → HV AL4 | 0.798 | 1.26e-07 | 5.78e-08 | 14 | 0 | 856 | 14 | 0 | 0 | 3202 | 10900 |
| V1L2/3 → HV AL5 | 1.136 | 1.84e-11 | 4.61e-12 | 13 | 0 | 429 | 13 | 0 | 0 | 2444 | 4028 |
| V1L4 → V1L2/3 | 0.335 | 5.41e-03 | 4.51e-03 | 6 | 0 | 1784 | 6 | 0 | 0 | 3107 | 16451 |
| V1L4 → V1L4 | 0.422 | 1.22e-04 | 8.12e-05 | 6 | 0 | 1865 | 6 | 0 | 0 | 4503 | 10073 |
| V1L4 → V1L5 | 0.435 | 1.04e-04 | 6.51e-05 | 6 | 0 | 1138 | 6 | 0 | 0 | 3365 | 4636 |
| V1L5 → V1L4 | 0.407 | 1.85e-04 | 1.31e-04 | 6 | 0 | 1769 | 6 | 0 | 0 | 3980 | 10686 |
| V1L5 → V1L5 | 0.523 | 6.94e-09 | 2.60e-09 | 6 | 0 | 1145 | 6 | 0 | 0 | 3721 | 4280 |
| HV AL2/3 → V1L2/3 | 0.067 | 3.37e-01 | 3.09e-01 | 36 | 0 | 2626 | 36 | 0 | 0 | 12670 | 104893 |
| HV AL2/3 → V1L4 | -0.024 | 8.66e-01 | 8.30e-01 | 28 | 0 | 1882 | 28 | 0 | 0 | 5122 | 63511 |
| HV AL2/3 → V1L5 | 0.361 | 4.28e-08 | 1.78e-08 | 59 | 0 | 1172 | 59 | 0 | 0 | 12966 | 66476 |
| HV AL2/3 → HV AL2/3 | 1.089 | 3.19e-121 | 1.33e-122 | 45 | 0 | 1264 | 45 | 0 | 0 | 19194 | 48691 |
| HV AL2/3 → HV AL4 | 0.831 | 3.91e-42 | 4.89e-43 | 38 | 0 | 893 | 38 | 0 | 0 | 13326 | 24937 |
| HV AL2/3 → HV AL5 | 0.280 | 1.10e-05 | 6.07e-06 | 62 | 0 | 439 | 62 | 0 | 0 | 13451 | 17560 |
| HV AL4 → HV AL2/3 | 0.633 | 5.28e-09 | 1.76e-09 | 12 | 0 | 1233 | 12 | 0 | 0 | 5899 | 12092 |
| HV AL4 → HV AL4 | 0.679 | 1.93e-09 | 5.62e-10 | 12 | 0 | 893 | 12 | 0 | 0 | 5266 | 6729 |
| HV AL4 → HV AL5 | 0.355 | 7.93e-03 | 6.94e-03 | 11 | 0 | 434 | 11 | 0 | 0 | 2992 | 2477 |
| HV AL5 → V1L5 | -0.013 | 9.11e-01 | 9.11e-01 | 14 | 0 | 1093 | 14 | 0 | 0 | 3539 | 15315 |
| HV AL5 → HV AL2/3 | 0.332 | 1.38e-03 | 1.04e-03 | 17 | 0 | 1236 | 17 | 0 | 0 | 7564 | 18063 |
| HV AL5 → HV AL4 | 0.326 | 4.43e-03 | 3.51e-03 | 17 | 0 | 896 | 17 | 0 | 0 | 6110 | 11017 |
| HV AL5 → HV AL5 | 0.458 | 1.10e-05 | 6.39e-06 | 19 | 0 | 439 | 19 | 0 | 0 | 5390 | 4009 |

**Supplemental Table 24.** Estimated marginal means of linear trends for the effect of in vivo signal correlation on *N*_*syn*_*/mm L*_*d*_ in different projection types across brain areas and layers. z and p-value are the z statistics and p-value of the marginal mean linear trends estimated from the fitted GLMMs. adjusted p-value is the adjusted p value through the BH multicomparison correction procedure.

| Projection type | Coefficient | adjusted p-value | p-value | # of presynaptic neurons | # of postsynaptic neurons | # of ADP control neurons | # of same region control neurons | # of synapses | # of pre-post pairs | # of pre-ADP pairs | # of pre-'same region' pairs |
| --- | --- | --- | --- | --- | --- | --- | --- | --- | --- | --- | --- |
| *V1L2/3* → *V1L2/3* | 1.110 | 2.08e-06 | 1.73e-07 | 20 | 604 | 2670 | 0 | 792 | 736 | 21247 | 0 |
| *V1L2/3* → *V1L4* | 0.617 | 1.33e-01 | 8.35e-02 | 19 | 197 | 2090 | 0 | 235 | 219 | 14010 | 0 |
| *V1L2/3* → *V1L5* | 1.820 | 3.11e-19 | 1.29e-20 | 20 | 311 | 1185 | 0 | 687 | 539 | 12384 | 0 |
| *V1L2/3* → *HVAL2/3* | 0.618 | 2.18e-01 | 1.54e-01 | 15 | 176 | 1169 | 0 | 208 | 196 | 4806 | 0 |
| *V1L2/3* → *HVAL4* | 1.080 | 1.33e-01 | 8.90e-02 | 14 | 80 | 856 | 0 | 89 | 82 | 3284 | 0 |
| *V1L2/3* → *HVAL5* | 1.225 | 3.34e-02 | 1.81e-02 | 13 | 91 | 429 | 0 | 131 | 106 | 2550 | 0 |
| *V1L4* → *V1L2/3* | 0.674 | 2.20e-01 | 1.65e-01 | 6 | 108 | 1784 | 0 | 120 | 110 | 3217 | 0 |
| *V1L4* → *V1L4* | 1.162 | 1.10e-02 | 4.57e-03 | 6 | 141 | 1865 | 0 | 155 | 146 | 4649 | 0 |
| *V1L4* → *V1L5* | 1.759 | 4.71e-05 | 7.85e-06 | 6 | 101 | 1138 | 0 | 130 | 110 | 3475 | 0 |
| *V1L5* → *V1L4* | -1.058 | 2.24e-01 | 1.78e-01 | 6 | 64 | 1769 | 0 | 65 | 64 | 4044 | 0 |
| *V1L5* → *V1L5* | 0.916 | 2.64e-02 | 1.21e-02 | 6 | 103 | 1145 | 0 | 121 | 104 | 3825 | 0 |
| *HVAL2/3* → *V1L2/3* | 0.880 | 8.21e-03 | 3.08e-03 | 36 | 381 | 2626 | 0 | 436 | 411 | 13081 | 0 |
| *HVAL2/3* → *V1L4* | 1.346 | 6.96e-02 | 4.06e-02 | 28 | 79 | 1882 | 0 | 88 | 81 | 5203 | 0 |
| *HVAL2/3* → *V1L5* | 1.378 | 4.12e-05 | 5.16e-06 | 59 | 213 | 1172 | 0 | 324 | 278 | 13244 | 0 |
| *HVAL2/3* → *HVAL2/3* | -0.083 | 7.29e-01 | 6.98e-01 | 45 | 519 | 1264 | 0 | 801 | 732 | 19926 | 0 |
| *HVAL2/3* → *HVAL4* | 1.223 | 4.13e-04 | 1.03e-04 | 38 | 204 | 893 | 0 | 301 | 258 | 13584 | 0 |
| *HVAL2/3* → *HVAL5* | 1.188 | 5.20e-05 | 1.08e-05 | 62 | 216 | 439 | 0 | 410 | 361 | 13812 | 0 |
| *HVAL4* → *HVAL2/3* | 0.843 | 3.34e-02 | 1.70e-02 | 12 | 259 | 1233 | 0 | 334 | 316 | 6215 | 0 |
| *HVAL4* → *HVAL4* | 1.349 | 8.21e-03 | 2.89e-03 | 12 | 138 | 893 | 0 | 174 | 155 | 5421 | 0 |
| *HVAL4* → *HVAL5* | 1.836 | 4.71e-03 | 1.37e-03 | 11 | 89 | 434 | 0 | 108 | 97 | 3089 | 0 |
| *HVAL5* → *V1L5* | -0.416 | 6.32e-01 | 5.79e-01 | 14 | 59 | 1093 | 0 | 67 | 62 | 3601 | 0 |
| *HVAL5* → *HVAL2/3* | 0.094 | 8.17e-01 | 8.17e-01 | 17 | 260 | 1236 | 0 | 331 | 308 | 7872 | 0 |
| *HVAL5* → *HVAL4* | 0.672 | 3.11e-01 | 2.62e-01 | 17 | 92 | 896 | 0 | 110 | 102 | 6212 | 0 |
| *HVAL5* → *HVAL5* | 0.460 | 3.11e-01 | 2.72e-01 | 19 | 148 | 439 | 0 | 214 | 196 | 5586 | 0 |

**Supplemental Table 25.** Estimated marginal means of linear trends for the effect of in silico signal correlation on *L*_*d*_ / neuron pair (synapses excluded) in different projection types across brain areas and layers. z and p-value are the z statistics and p-value of the marginal mean linear trends estimated from the fitted GLMMs. adjusted p-value is the adjusted p value through the BH multicomparison correction procedure.

| Projection type | Coefficient | adjusted p-value | p-value | # of presynaptic neurons | # of postsynaptic neurons | # of ADP control neurons | # of same region control neurons | # of synapses | # of pre-post pairs | # of pre-ADP pairs | # of pre-'same region' pairs |
| --- | --- | --- | --- | --- | --- | --- | --- | --- | --- | --- | --- |
| V1L2/3 → V1L2/3 | 1.692 | 6.20e-109 | 5.17e-110 | 20 | 0 | 2670 | 20 | 0 | 0 | 20511 | 44349 |
| V1L2/3 → V1L4 | 0.632 | 9.58e-10 | 5.19e-10 | 19 | 0 | 2090 | 19 | 0 | 0 | 13791 | 32623 |
| V1L2/3 → V1L5 | 0.929 | 1.39e-17 | 5.23e-18 | 20 | 0 | 1185 | 20 | 0 | 0 | 11845 | 14652 |
| V1L2/3 → HV AL2/3 | 2.276 | 5.44e-33 | 9.06e-34 | 15 | 0 | 1169 | 15 | 0 | 0 | 4610 | 18075 |
| V1L2/3 → HV AL4 | 1.520 | 4.26e-10 | 2.13e-10 | 14 | 0 | 856 | 14 | 0 | 0 | 3202 | 10900 |
| V1L2/3 → HV AL5 | 0.738 | 5.58e-03 | 4.65e-03 | 13 | 0 | 429 | 13 | 0 | 0 | 2444 | 4028 |
| V1L4 → V1L2/3 | 0.551 | 7.48e-03 | 6.54e-03 | 6 | 0 | 1784 | 6 | 0 | 0 | 3107 | 16451 |
| V1L4 → V1L4 | 0.525 | 1.65e-03 | 1.25e-03 | 6 | 0 | 1865 | 6 | 0 | 0 | 4503 | 10073 |
| V1L4 → V1L5 | 0.538 | 1.65e-03 | 1.31e-03 | 6 | 0 | 1138 | 6 | 0 | 0 | 3365 | 4636 |
| V1L5 → V1L4 | 0.491 | 7.49e-03 | 6.87e-03 | 6 | 0 | 1769 | 6 | 0 | 0 | 3980 | 10686 |
| V1L5 → V1L5 | 0.546 | 8.97e-04 | 5.98e-04 | 6 | 0 | 1145 | 6 | 0 | 0 | 3721 | 4280 |
| HV AL2/3 → V1L2/3 | -0.020 | 8.30e-01 | 8.30e-01 | 36 | 0 | 2626 | 36 | 0 | 0 | 12670 | 104893 |
| HV AL2/3 → V1L4 | 0.473 | 1.45e-03 | 1.03e-03 | 28 | 0 | 1882 | 28 | 0 | 0 | 5122 | 63511 |
| HV AL2/3 → V1L5 | 0.973 | 7.73e-29 | 1.61e-29 | 59 | 0 | 1172 | 59 | 0 | 0 | 12966 | 66476 |
| HV AL2/3 → HV AL2/3 | 2.087 | 3.32e-246 | 1.38e-247 | 45 | 0 | 1264 | 45 | 0 | 0 | 19194 | 48691 |
| HV AL2/3 → HV AL4 | 1.410 | 2.16e-69 | 2.70e-70 | 38 | 0 | 893 | 38 | 0 | 0 | 13326 | 24937 |
| HV AL2/3 → HV AL5 | 0.116 | 1.45e-01 | 1.39e-01 | 62 | 0 | 439 | 62 | 0 | 0 | 13451 | 17560 |
| HV AL4 → HV AL2/3 | 1.264 | 2.34e-22 | 6.83e-23 | 12 | 0 | 1233 | 12 | 0 | 0 | 5899 | 12092 |
| HV AL4 → HV AL4 | 0.975 | 1.87e-13 | 8.58e-14 | 12 | 0 | 893 | 12 | 0 | 0 | 5266 | 6729 |
| HV AL4 → HV AL5 | 0.631 | 1.69e-04 | 1.05e-04 | 11 | 0 | 434 | 11 | 0 | 0 | 2992 | 2477 |
| HV AL5 → V1L5 | 0.873 | 5.35e-07 | 3.12e-07 | 14 | 0 | 1093 | 14 | 0 | 0 | 3539 | 15315 |
| HV AL5 → HV AL2/3 | 1.266 | 3.38e-24 | 8.45e-25 | 17 | 0 | 1236 | 17 | 0 | 0 | 7564 | 18063 |
| HV AL5 → HV AL4 | 1.146 | 1.40e-16 | 5.83e-17 | 17 | 0 | 896 | 17 | 0 | 0 | 6110 | 11017 |
| HV AL5 → HV AL5 | 1.215 | 9.66e-22 | 3.22e-22 | 19 | 0 | 439 | 19 | 0 | 0 | 5390 | 4009 |

**Supplemental Table 26.** Estimated marginal means of linear trends for the effect of in silico signal correlation on *N*_*syn*_*/mm L*_*d*_ in different projection types across brain areas and layers. z and p-value are the z statistics and p-value of the marginal mean linear trends estimated from the fitted GLMMs. adjusted p-value is the adjusted p value through the BH multicomparison correction procedure.

| Projection type | Coefficient | adjusted p-value | p-value | # of presynaptic neurons | # of postsynaptic neurons | # of ADP control neurons | # of same region control neurons | # of synapses | # of pre-post pairs | # of pre-ADP pairs | # of pre-'same region' pairs |
| --- | --- | --- | --- | --- | --- | --- | --- | --- | --- | --- | --- |
| V1L2/3 → V1L2/3 | 2.026 | 2.87e-09 | 3.58e-10 | 20 | 604 | 2670 | 0 | 792 | 736 | 21247 | 0 |
| V1L2/3 → V1L4 | 2.973 | 1.67e-07 | 3.47e-08 | 19 | 197 | 2090 | 0 | 235 | 219 | 14010 | 0 |
| V1L2/3 → V1L5 | 4.022 | 1.75e-39 | 7.29e-41 | 20 | 311 | 1185 | 0 | 687 | 539 | 12384 | 0 |
| V1L2/3 → HVAL2/3 | 0.808 | 3.52e-01 | 2.93e-01 | 15 | 176 | 1169 | 0 | 208 | 196 | 4806 | 0 |
| V1L2/3 → HVAL4 | 2.167 | 9.18e-02 | 6.12e-02 | 14 | 80 | 856 | 0 | 89 | 82 | 3284 | 0 |
| V1L2/3 → HVAL5 | 3.941 | 3.48e-06 | 1.16e-06 | 13 | 91 | 429 | 0 | 131 | 106 | 2550 | 0 |
| V1L4 → V1L2/3 | 1.097 | 2.48e-01 | 1.86e-01 | 6 | 108 | 1784 | 0 | 120 | 110 | 3217 | 0 |
| V1L4 → V1L4 | 2.180 | 5.08e-04 | 1.90e-04 | 6 | 141 | 1865 | 0 | 155 | 146 | 4649 | 0 |
| V1L4 → V1L5 | 3.194 | 7.21e-09 | 1.20e-09 | 6 | 101 | 1138 | 0 | 130 | 110 | 3475 | 0 |
| V1L5 → V1L4 | -1.271 | 3.52e-01 | 2.87e-01 | 6 | 64 | 1769 | 0 | 65 | 64 | 4044 | 0 |
| V1L5 → V1L5 | 1.508 | 5.43e-02 | 2.94e-02 | 6 | 103 | 1145 | 0 | 121 | 104 | 3825 | 0 |
| HVAL2/3 → V1L2/3 | 0.837 | 9.18e-02 | 5.90e-02 | 36 | 381 | 2626 | 0 | 436 | 411 | 13081 | 0 |
| HVAL2/3 → V1L4 | 1.304 | 2.15e-01 | 1.52e-01 | 28 | 79 | 1882 | 0 | 88 | 81 | 5203 | 0 |
| HVAL2/3 → V1L5 | 2.730 | 1.70e-10 | 1.42e-11 | 59 | 213 | 1172 | 0 | 324 | 278 | 13244 | 0 |
| HVAL2/3 → HVAL2/3 | -0.108 | 7.41e-01 | 7.19e-01 | 45 | 519 | 1264 | 0 | 801 | 732 | 19926 | 0 |
| HVAL2/3 → HVAL4 | 1.585 | 9.99e-04 | 4.16e-04 | 38 | 204 | 893 | 0 | 301 | 258 | 13584 | 0 |
| HVAL2/3 → HVAL5 | 1.834 | 2.24e-06 | 5.61e-07 | 62 | 216 | 439 | 0 | 410 | 361 | 13812 | 0 |
| HVAL4 → HVAL2/3 | 1.203 | 2.36e-02 | 1.18e-02 | 12 | 259 | 1233 | 0 | 334 | 316 | 6215 | 0 |
| HVAL4 → HVAL4 | 2.515 | 2.28e-06 | 6.64e-07 | 12 | 138 | 893 | 0 | 174 | 155 | 5421 | 0 |
| HVAL4 → HVAL5 | 2.154 | 5.01e-03 | 2.30e-03 | 11 | 89 | 434 | 0 | 108 | 97 | 3089 | 0 |
| HVAL5 → V1L5 | 1.878 | 8.64e-02 | 5.04e-02 | 14 | 59 | 1093 | 0 | 67 | 62 | 3601 | 0 |
| HVAL5 → HVAL2/3 | 0.308 | 5.91e-01 | 5.17e-01 | 17 | 260 | 1236 | 0 | 331 | 308 | 7872 | 0 |
| HVAL5 → HVAL4 | 0.248 | 7.41e-01 | 7.41e-01 | 17 | 92 | 896 | 0 | 110 | 102 | 6212 | 0 |
| HVAL5 → HVAL5 | 0.313 | 6.00e-01 | 5.50e-01 | 19 | 148 | 439 | 0 | 214 | 196 | 5586 | 0 |

**Supplemental Table 27.** Estimated marginal means of linear trends for the effect of feature weight similarity on *L*_*d*_ / neuron pair (synapses excluded) in different projection types across brain areas and layers. z and p-value are the z statistics and p-value of the marginal mean linear trends estimated from the fitted GLMMs. adjusted p-value is the adjusted p value through the BH multicomparison correction procedure.

| Projection type | Coefficient | adjusted p-value | p-value | # of presynaptic neurons | # of postsynaptic neurons | # of ADP control neurons | # of same region control neurons | # of synapses | # of pre-post pairs | # of pre-ADP pairs | # of pre-'same region' pairs |
| --- | --- | --- | --- | --- | --- | --- | --- | --- | --- | --- | --- |
| V1L2/3 → V1L2/3 | 1.005 | 8.41e-38 | 1.05e-38 | 20 | 0 | 2670 | 20 | 0 | 0 | 20511 | 44349 |
| V1L2/3 → V1L4 | 0.512 | 9.17e-07 | 4.59e-07 | 19 | 0 | 2090 | 19 | 0 | 0 | 13791 | 32623 |
| V1L2/3 → V1L5 | 0.602 | 8.40e-08 | 3.85e-08 | 20 | 0 | 1185 | 20 | 0 | 0 | 11845 | 14652 |
| V1L2/3 → HVAL2/3 | 0.849 | 1.12e-05 | 6.97e-06 | 15 | 0 | 1169 | 15 | 0 | 0 | 4610 | 18075 |
| V1L2/3 → HVAL4 | 0.804 | 1.27e-03 | 9.01e-04 | 14 | 0 | 856 | 14 | 0 | 0 | 3202 | 10900 |
| V1L2/3 → HVAL5 | 0.410 | 1.34e-01 | 1.28e-01 | 13 | 0 | 429 | 13 | 0 | 0 | 2444 | 4028 |
| V1L4 → V1L2/3 | 0.257 | 2.09e-01 | 2.09e-01 | 6 | 0 | 1784 | 6 | 0 | 0 | 3107 | 16451 |
| V1L4 → V1L4 | 0.769 | 1.79e-06 | 9.70e-07 | 6 | 0 | 1865 | 6 | 0 | 0 | 4503 | 10073 |
| V1L4 → V1L5 | 0.663 | 2.60e-04 | 1.73e-04 | 6 | 0 | 1138 | 6 | 0 | 0 | 3365 | 4636 |
| V1L5 → V1L4 | 0.465 | 1.66e-02 | 1.25e-02 | 6 | 0 | 1769 | 6 | 0 | 0 | 3980 | 10686 |
| V1L5 → V1L5 | 0.361 | 3.27e-02 | 2.87e-02 | 6 | 0 | 1145 | 6 | 0 | 0 | 3721 | 4280 |
| HVAL2/3 → V1L2/3 | 0.251 | 2.75e-02 | 2.17e-02 | 36 | 0 | 2626 | 36 | 0 | 0 | 12670 | 104893 |
| HVAL2/3 → V1L4 | 0.378 | 3.17e-02 | 2.64e-02 | 28 | 0 | 1882 | 28 | 0 | 0 | 5122 | 63511 |
| HVAL2/3 → V1L5 | 1.078 | 5.07e-22 | 1.06e-22 | 59 | 0 | 1172 | 59 | 0 | 0 | 12966 | 66476 |
| HVAL2/3 → HVAL2/3 | 2.692 | 3.67e-248 | 1.53e-249 | 45 | 0 | 1264 | 45 | 0 | 0 | 19194 | 48691 |
| HVAL2/3 → HVAL4 | 2.013 | 5.69e-83 | 4.74e-84 | 38 | 0 | 893 | 38 | 0 | 0 | 13326 | 24937 |
| HVAL2/3 → HVAL5 | 0.885 | 1.56e-17 | 5.20e-18 | 62 | 0 | 439 | 62 | 0 | 0 | 13451 | 17560 |
| HVAL4 → HVAL2/3 | 1.560 | 2.27e-24 | 3.78e-25 | 12 | 0 | 1233 | 12 | 0 | 0 | 5899 | 12092 |
| HVAL4 → HVAL4 | 1.436 | 1.03e-21 | 2.57e-22 | 12 | 0 | 893 | 12 | 0 | 0 | 5266 | 6729 |
| HVAL4 → HVAL5 | 0.884 | 2.15e-06 | 1.25e-06 | 11 | 0 | 434 | 11 | 0 | 0 | 2992 | 2477 |
| HVAL5 → V1L5 | 1.169 | 1.05e-09 | 4.38e-10 | 14 | 0 | 1093 | 14 | 0 | 0 | 3539 | 15315 |
| HVAL5 → HVAL2/3 | 0.273 | 6.10e-02 | 5.59e-02 | 17 | 0 | 1236 | 17 | 0 | 0 | 7564 | 18063 |
| HVAL5 → HVAL4 | 1.195 | 3.89e-14 | 1.46e-14 | 17 | 0 | 896 | 17 | 0 | 0 | 6110 | 11017 |
| HVAL5 → HVAL5 | 1.306 | 1.37e-19 | 3.99e-20 | 19 | 0 | 439 | 19 | 0 | 0 | 5390 | 4009 |

**Supplemental Table 28.** Estimated marginal means of linear trends for the effect of feature weight similarity on *N*_*syn*_*/mm L*_*d*_ in different projection types across brain areas and layers. z and p-value are the z statistics and p-value of the marginal mean linear trends estimated from the fitted GLMMs. adjusted p-value is the adjusted p value through the BH multicomparison correction procedure.

| Projection type | Coefficient | adjusted p-value | p-value | # of presynaptic neurons | # of postsynaptic neurons | # of ADP control neurons | # of same region control neurons | # of synapses | # of pre-post pairs | # of pre-ADP pairs | # of pre-'same region' pairs |
| --- | --- | --- | --- | --- | --- | --- | --- | --- | --- | --- | --- |
| V1L2/3 → V1L2/3 | 1.840 | 6.63e-07 | 1.11e-07 | 20 | 604 | 2670 | 0 | 792 | 736 | 21247 | 0 |
| V1L2/3 → V1L4 | 2.015 | 3.78e-03 | 1.57e-03 | 19 | 197 | 2090 | 0 | 235 | 219 | 14010 | 0 |
| V1L2/3 → V1L5 | 3.855 | 1.05e-25 | 4.39e-27 | 20 | 311 | 1185 | 0 | 687 | 539 | 12384 | 0 |
| V1L2/3 → HVAL2/3 | 0.715 | 3.77e-01 | 3.45e-01 | 15 | 176 | 1169 | 0 | 208 | 196 | 4806 | 0 |
| V1L2/3 → HVAL4 | -0.143 | 9.04e-01 | 9.04e-01 | 14 | 80 | 856 | 0 | 89 | 82 | 3284 | 0 |
| V1L2/3 → HVAL5 | 3.330 | 4.87e-04 | 1.83e-04 | 13 | 91 | 429 | 0 | 131 | 106 | 2550 | 0 |
| V1L4 → V1L2/3 | 1.005 | 3.28e-01 | 2.59e-01 | 6 | 108 | 1784 | 0 | 120 | 110 | 3217 | 0 |
| V1L4 → V1L4 | 2.416 | 4.52e-04 | 1.51e-04 | 6 | 141 | 1865 | 0 | 155 | 146 | 4649 | 0 |
| V1L4 → V1L5 | 4.052 | 8.61e-08 | 7.18e-09 | 6 | 101 | 1138 | 0 | 130 | 110 | 3475 | 0 |
| V1L5 → V1L4 | -1.410 | 3.11e-01 | 2.33e-01 | 6 | 64 | 1769 | 0 | 65 | 64 | 4044 | 0 |
| V1L5 → V1L5 | 0.304 | 7.36e-01 | 7.05e-01 | 6 | 103 | 1145 | 0 | 121 | 104 | 3825 | 0 |
| HVAL2/3 → V1L2/3 | 1.248 | 3.23e-02 | 1.89e-02 | 36 | 381 | 2626 | 0 | 436 | 411 | 13081 | 0 |
| HVAL2/3 → V1L4 | 3.178 | 8.70e-03 | 4.35e-03 | 28 | 79 | 1882 | 0 | 88 | 81 | 5203 | 0 |
| HVAL2/3 → V1L5 | 3.210 | 1.28e-07 | 1.59e-08 | 59 | 213 | 1172 | 0 | 324 | 278 | 13244 | 0 |
| HVAL2/3 → HVAL2/3 | 0.351 | 3.77e-01 | 3.44e-01 | 45 | 519 | 1264 | 0 | 801 | 732 | 19926 | 0 |
| HVAL2/3 → HVAL4 | 2.861 | 7.44e-06 | 1.86e-06 | 38 | 204 | 893 | 0 | 301 | 258 | 13584 | 0 |
| HVAL2/3 → HVAL5 | 2.452 | 3.18e-06 | 6.63e-07 | 62 | 216 | 439 | 0 | 410 | 361 | 13812 | 0 |
| HVAL4 → HVAL2/3 | 1.416 | 2.23e-02 | 1.21e-02 | 12 | 259 | 1233 | 0 | 334 | 316 | 6215 | 0 |
| HVAL4 → HVAL4 | 2.614 | 1.54e-04 | 4.51e-05 | 12 | 138 | 893 | 0 | 174 | 155 | 5421 | 0 |
| HVAL4 → HVAL5 | 2.486 | 8.70e-03 | 4.12e-03 | 11 | 89 | 434 | 0 | 108 | 97 | 3089 | 0 |
| HVAL5 → V1L5 | 1.423 | 2.89e-01 | 2.05e-01 | 14 | 59 | 1093 | 0 | 67 | 62 | 3601 | 0 |
| HVAL5 → HVAL2/3 | 0.794 | 2.51e-01 | 1.67e-01 | 17 | 260 | 1236 | 0 | 331 | 308 | 7872 | 0 |
| HVAL5 → HVAL4 | -0.926 | 3.54e-01 | 2.95e-01 | 17 | 92 | 896 | 0 | 110 | 102 | 6212 | 0 |
| HVAL5 → HVAL5 | 1.090 | 1.09e-01 | 6.81e-02 | 19 | 148 | 439 | 0 | 214 | 196 | 5586 | 0 |

**Supplemental Table 29.** Estimated marginal means of linear trends for the effect of receptive field center distance on *L*_*d*_ / neuron pair (synapses excluded) in different projection types across brain areas and layers. z and p-value are the z statistics and p-value of the marginal mean linear trends estimated from the fitted GLMMs. adjusted p-value is the adjusted p value through the BH multicomparison correction procedure.

| Projection type | Coefficient | adjusted p-value | p-value | # of presynaptic neurons | # of postsynaptic neurons | # of ADP control neurons | # of same region control neurons | # of synapses | # of pre-post pairs | # of pre-ADP pairs | # of pre-'same region' pairs |
| --- | --- | --- | --- | --- | --- | --- | --- | --- | --- | --- | --- |
| *V1L2/3* → *V1L2/3* | -0.167 | 0.00e+00 | 0.00e+00 | 20 | 0 | 2670 | 20 | 0 | 0 | 20511 | 44349 |
| *V1L2/3* → *V1L4* | -0.141 | 0.00e+00 | 0.00e+00 | 19 | 0 | 2090 | 19 | 0 | 0 | 13791 | 32623 |
| *V1L2/3* → *V1L5* | -0.111 | 0.00e+00 | 0.00e+00 | 20 | 0 | 1185 | 20 | 0 | 0 | 11845 | 14652 |
| *V1L2/3* → *HVAL2/3* | -0.032 | 7.39e-21 | 6.16e-21 | 15 | 0 | 1169 | 15 | 0 | 0 | 4610 | 18075 |
| *V1L2/3* → *HVAL4* | -0.030 | 5.27e-14 | 4.83e-14 | 14 | 0 | 856 | 14 | 0 | 0 | 3202 | 10900 |
| *V1L2/3* → *HVAL5* | -0.014 | 4.94e-03 | 4.94e-03 | 13 | 0 | 429 | 13 | 0 | 0 | 2444 | 4028 |
| *V1L4* → *V1L2/3* | -0.097 | 1.11e-68 | 7.42e-69 | 6 | 0 | 1784 | 6 | 0 | 0 | 3107 | 16451 |
| *V1L4* → *V1L4* | -0.136 | 1.76e-189 | 5.14e-190 | 6 | 0 | 1865 | 6 | 0 | 0 | 4503 | 10073 |
| *V1L4* → *V1L5* | -0.134 | 8.03e-134 | 4.01e-134 | 6 | 0 | 1138 | 6 | 0 | 0 | 3365 | 4636 |
| *V1L5* → *V1L4* | -0.053 | 2.71e-27 | 2.14e-27 | 6 | 0 | 1769 | 6 | 0 | 0 | 3980 | 10686 |
| *V1L5* → *V1L5* | -0.092 | 1.19e-80 | 7.44e-81 | 6 | 0 | 1145 | 6 | 0 | 0 | 3721 | 4280 |
| *HVAL2/3* → *V1L2/3* | -0.015 | 7.37e-11 | 7.06e-11 | 36 | 0 | 2626 | 36 | 0 | 0 | 12670 | 104893 |
| *HVAL2/3* → *V1L4* | -0.031 | 1.82e-16 | 1.60e-16 | 28 | 0 | 1882 | 28 | 0 | 0 | 5122 | 63511 |
| *HVAL2/3* → *V1L5* | -0.035 | 1.25e-46 | 9.39e-47 | 59 | 0 | 1172 | 59 | 0 | 0 | 12966 | 66476 |
| *HVAL2/3* → *HVAL2/3* | -0.099 | 0.00e+00 | 0.00e+00 | 45 | 0 | 1264 | 45 | 0 | 0 | 19194 | 48691 |
| *HVAL2/3* → *HVAL4* | -0.083 | 0.00e+00 | 0.00e+00 | 38 | 0 | 893 | 38 | 0 | 0 | 13326 | 24937 |
| *HVAL2/3* → *HVAL5* | -0.047 | 1.41e-112 | 7.63e-113 | 62 | 0 | 439 | 62 | 0 | 0 | 13451 | 17560 |
| *HVAL4* → *HVAL2/3* | -0.084 | 4.83e-169 | 1.81e-169 | 12 | 0 | 1233 | 12 | 0 | 0 | 5899 | 12092 |
| *HVAL4* → *HVAL4* | -0.093 | 2.33e-215 | 5.82e-216 | 12 | 0 | 893 | 12 | 0 | 0 | 5266 | 6729 |
| *HVAL4* → *HVAL5* | -0.109 | 5.56e-172 | 1.85e-172 | 11 | 0 | 434 | 11 | 0 | 0 | 2992 | 2477 |
| *HVAL5* → *V1L5* | -0.067 | 3.07e-53 | 2.18e-53 | 14 | 0 | 1093 | 14 | 0 | 0 | 3539 | 15315 |
| *HVAL5* → *HVAL2/3* | -0.052 | 3.63e-86 | 2.12e-86 | 17 | 0 | 1236 | 17 | 0 | 0 | 7564 | 18063 |
| *HVAL5* → *HVAL4* | -0.071 | 8.03e-134 | 3.74e-134 | 17 | 0 | 896 | 17 | 0 | 0 | 6110 | 11017 |
| *HVAL5* → *HVAL5* | -0.082 | 4.99e-159 | 2.08e-159 | 19 | 0 | 439 | 19 | 0 | 0 | 5390 | 4009 |

**Supplemental Table 30.** Estimated marginal means of linear trends for the effect of receptive field center distance on *N*_*syn*_*/mm L*_*d*_ in different projection types across brain areas and layers. z and p-value are the z statistics and p-value of the marginal mean linear trends estimated from the fitted GLMMs. adjusted p-value is the adjusted p value through the BH multicomparison correction procedure.

| Projection type | Coefficient | adjusted p-value | p-value | # of presynaptic neurons | # of postsynaptic neurons | # of ADP control neurons | # of same region control neurons | # of synapses | # of pre-post pairs | # of pre-ADP pairs | # of pre-'same region' pairs |
| --- | --- | --- | --- | --- | --- | --- | --- | --- | --- | --- | --- |
| V1L2/3 → V1L2/3 | 0.020 | 2.56e-01 | 4.26e-02 | 20 | 604 | 2670 | 0 | 792 | 736 | 21247 | 0 |
| V1L2/3 → V1L4 | 0.050 | 5.19e-02 | 2.16e-03 | 19 | 197 | 2090 | 0 | 235 | 219 | 14010 | 0 |
| V1L2/3 → V1L5 | 0.012 | 5.26e-01 | 2.27e-01 | 20 | 311 | 1185 | 0 | 687 | 539 | 12384 | 0 |
| V1L2/3 → HVAL2/3 | 0.008 | 8.34e-01 | 6.26e-01 | 15 | 176 | 1169 | 0 | 208 | 196 | 4806 | 0 |
| V1L2/3 → HVAL4 | 0.014 | 8.18e-01 | 5.11e-01 | 14 | 80 | 856 | 0 | 89 | 82 | 3284 | 0 |
| V1L2/3 → HVAL5 | 0.001 | 9.50e-01 | 9.50e-01 | 13 | 91 | 429 | 0 | 131 | 106 | 2550 | 0 |
| V1L4 → V1L2/3 | 0.020 | 8.10e-01 | 4.59e-01 | 6 | 108 | 1784 | 0 | 120 | 110 | 3217 | 0 |
| V1L4 → V1L4 | 0.060 | 6.82e-02 | 5.69e-03 | 6 | 141 | 1865 | 0 | 155 | 146 | 4649 | 0 |
| V1L4 → V1L5 | 0.042 | 3.98e-01 | 8.30e-02 | 6 | 101 | 1138 | 0 | 130 | 110 | 3475 | 0 |
| V1L5 → V1L4 | -0.016 | 8.34e-01 | 6.22e-01 | 6 | 64 | 1769 | 0 | 65 | 64 | 4044 | 0 |
| V1L5 → V1L5 | 0.034 | 5.26e-01 | 2.02e-01 | 6 | 103 | 1145 | 0 | 121 | 104 | 3825 | 0 |
| HVAL2/3 → V1L2/3 | -0.015 | 5.26e-01 | 2.19e-01 | 36 | 381 | 2626 | 0 | 436 | 411 | 13081 | 0 |
| HVAL2/3 → V1L4 | 0.004 | 9.39e-01 | 8.88e-01 | 28 | 79 | 1882 | 0 | 88 | 81 | 5203 | 0 |
| HVAL2/3 → V1L5 | -0.020 | 5.26e-01 | 1.37e-01 | 59 | 213 | 1172 | 0 | 324 | 278 | 13244 | 0 |
| HVAL2/3 → HVAL2/3 | 0.004 | 8.49e-01 | 6.74e-01 | 45 | 519 | 1264 | 0 | 801 | 732 | 19926 | 0 |
| HVAL2/3 → HVAL4 | -0.002 | 9.39e-01 | 8.99e-01 | 38 | 204 | 893 | 0 | 301 | 258 | 13584 | 0 |
| HVAL2/3 → HVAL5 | -0.004 | 8.49e-01 | 7.08e-01 | 62 | 216 | 439 | 0 | 410 | 361 | 13812 | 0 |
| HVAL4 → HVAL2/3 | 0.013 | 5.26e-01 | 2.41e-01 | 12 | 259 | 1233 | 0 | 334 | 316 | 6215 | 0 |
| HVAL4 → HVAL4 | 0.014 | 6.51e-01 | 3.26e-01 | 12 | 138 | 893 | 0 | 174 | 155 | 5421 | 0 |
| HVAL4 → HVAL5 | 0.048 | 1.25e-01 | 1.57e-02 | 11 | 89 | 434 | 0 | 108 | 97 | 3089 | 0 |
| HVAL5 → V1L5 | 0.016 | 8.34e-01 | 5.74e-01 | 14 | 59 | 1093 | 0 | 67 | 62 | 3601 | 0 |
| HVAL5 → HVAL2/3 | 0.008 | 8.10e-01 | 4.72e-01 | 17 | 260 | 1236 | 0 | 331 | 308 | 7872 | 0 |
| HVAL5 → HVAL4 | 0.021 | 5.26e-01 | 2.11e-01 | 17 | 92 | 896 | 0 | 110 | 102 | 6212 | 0 |
| HVAL5 → HVAL5 | -0.004 | 8.78e-01 | 7.69e-01 | 19 | 148 | 439 | 0 | 214 | 196 | 5586 | 0 |

**Supplemental Table 31.** Estimated marginal means of linear trends for the effect of in silico ΔOri on *L*_*d*_ / neuron pair (synapses excluded) in different projection types across brain areas and layers. z and p-value are the z statistics and p-value of the marginal mean linear trends estimated from the fitted GLMMs. adjusted p-value is the adjusted p value through the BH multicomparison correction procedure.

| Projection type | Coefficient | adjusted p-value | p-value | # of presynaptic neurons | # of postsynaptic neurons | # of ADP control neurons | # of same region control neurons | # of synapses | # of pre-post pairs | # of pre-ADP pairs | # of pre-'same region' pairs |
| --- | --- | --- | --- | --- | --- | --- | --- | --- | --- | --- | --- |
| *V1L2/3* → *V1L2/3* | -0.003 | 1.22e-10 | 7.61e-12 | 12 | 0 | 1409 | 12 | 0 | 0 | 6927 | 14702 |
| *V1L2/3* → *V1L4* | -0.004 | 3.52e-07 | 4.40e-08 | 9 | 0 | 1050 | 9 | 0 | 0 | 4190 | 7845 |
| *V1L2/3* → *V1L5* | -0.001 | 5.15e-01 | 3.22e-01 | 12 | 0 | 701 | 12 | 0 | 0 | 4527 | 4791 |
| *V1L2/3* → *HVAL2/3* | -0.006 | 3.69e-06 | 6.93e-07 | 7 | 0 | 487 | 7 | 0 | 0 | 1354 | 3258 |
| *V1L2/3* → *HVAL5* | -0.003 | 9.32e-02 | 4.21e-02 | 8 | 0 | 224 | 8 | 0 | 0 | 786 | 1249 |
| *HVAL2/3* → *V1L2/3* | 0.003 | 4.97e-04 | 1.24e-04 | 20 | 0 | 1299 | 20 | 0 | 0 | 4401 | 31930 |
| *HVAL2/3* → *V1L4* | -0.000 | 9.42e-01 | 9.42e-01 | 16 | 0 | 885 | 16 | 0 | 0 | 1706 | 19798 |
| *HVAL2/3* → *V1L5* | 0.001 | 9.32e-02 | 4.66e-02 | 30 | 0 | 689 | 30 | 0 | 0 | 4433 | 19345 |
| *HVAL2/3* → *HVAL2/3* | 0.000 | 8.54e-01 | 7.47e-01 | 17 | 0 | 557 | 17 | 0 | 0 | 4103 | 7038 |
| *HVAL2/3* → *HVAL4* | 0.000 | 8.41e-01 | 6.83e-01 | 16 | 0 | 397 | 16 | 0 | 0 | 2812 | 4472 |
| *HVAL2/3* → *HVAL5* | -0.001 | 4.10e-01 | 2.31e-01 | 27 | 0 | 229 | 27 | 0 | 0 | 3420 | 3482 |
| *HVAL4* → *HVAL2/3* | -0.001 | 6.16e-01 | 4.24e-01 | 7 | 0 | 499 | 7 | 0 | 0 | 1262 | 3344 |
| *HVAL4* → *HVAL5* | -0.000 | 8.41e-01 | 6.74e-01 | 7 | 0 | 225 | 7 | 0 | 0 | 937 | 848 |
| *HVAL5* → *V1L5* | 0.000 | 9.42e-01 | 9.22e-01 | 9 | 0 | 603 | 9 | 0 | 0 | 1414 | 5725 |
| *HVAL5* → *HVAL2/3* | 0.002 | 2.94e-02 | 9.20e-03 | 12 | 0 | 541 | 12 | 0 | 0 | 2307 | 5594 |
| *HVAL5* → *HVAL5* | -0.002 | 9.32e-02 | 4.49e-02 | 14 | 0 | 228 | 14 | 0 | 0 | 2125 | 1430 |

**Supplemental Table 32.** Estimated marginal means of linear trends for the effect of in silico ΔOri on *N*_*syn*_*/mm L*_*d*_ in different projection types across brain areas and layers. z and p-value are the z statistics and p-value of the marginal mean linear trends estimated from the fitted GLMMs. adjusted p-value is the adjusted p value through the BH multicomparison correction procedure.

| Projection type | Coefficient | adjusted p-value | p-value | # of presynaptic neurons | # of postsynaptic neurons | # of ADP control neurons | # of same region control neurons | # of synapses | # of pre-post pairs | # of pre-ADP pairs | # of pre-'same region' pairs |
| --- | --- | --- | --- | --- | --- | --- | --- | --- | --- | --- | --- |
| V1L2/3 → V1L2/3 | -0.003 | 4.73e-01 | 1.80e-01 | 12 | 246 | 1409 | 0 | 296 | 272 | 7199 | 0 |
| V1L2/3 → V1L4 | 0.005 | 4.73e-01 | 1.82e-01 | 9 | 82 | 1050 | 0 | 91 | 83 | 4273 | 0 |
| V1L2/3 → V1L5 | -0.009 | 1.66e-03 | 1.04e-04 | 12 | 165 | 701 | 0 | 323 | 245 | 4772 | 0 |
| V1L2/3 → HVAL2/3 | 0.001 | 8.89e-01 | 8.89e-01 | 7 | 52 | 487 | 0 | 54 | 53 | 1407 | 0 |
| V1L2/3 → HVAL5 | -0.014 | 8.07e-02 | 1.01e-02 | 8 | 38 | 224 | 0 | 51 | 40 | 826 | 0 |
| HVAL2/3 → V1L2/3 | -0.004 | 4.73e-01 | 2.37e-01 | 20 | 123 | 1299 | 0 | 134 | 129 | 4530 | 0 |
| HVAL2/3 → V1L4 | -0.012 | 4.04e-01 | 7.57e-02 | 16 | 33 | 885 | 0 | 36 | 33 | 1739 | 0 |
| HVAL2/3 → V1L5 | -0.002 | 7.20e-01 | 6.06e-01 | 30 | 97 | 689 | 0 | 123 | 109 | 4542 | 0 |
| HVAL2/3 → HVAL2/3 | -0.001 | 7.20e-01 | 6.07e-01 | 17 | 145 | 557 | 0 | 192 | 177 | 4280 | 0 |
| HVAL2/3 → HVAL4 | 0.002 | 7.31e-01 | 6.85e-01 | 16 | 59 | 397 | 0 | 70 | 67 | 2879 | 0 |
| HVAL2/3 → HVAL5 | 0.004 | 4.73e-01 | 2.07e-01 | 27 | 80 | 229 | 0 | 122 | 106 | 3526 | 0 |
| HVAL4 → HVAL2/3 | 0.005 | 5.95e-01 | 3.35e-01 | 7 | 55 | 499 | 0 | 59 | 59 | 1321 | 0 |
| HVAL4 → HVAL5 | -0.009 | 4.73e-01 | 1.49e-01 | 7 | 29 | 225 | 0 | 32 | 30 | 967 | 0 |
| HVAL5 → V1L5 | -0.003 | 7.20e-01 | 6.30e-01 | 9 | 25 | 603 | 0 | 31 | 26 | 1440 | 0 |
| HVAL5 → HVAL2/3 | -0.002 | 7.20e-01 | 5.38e-01 | 12 | 87 | 541 | 0 | 106 | 98 | 2405 | 0 |
| HVAL5 → HVAL5 | -0.004 | 6.13e-01 | 3.83e-01 | 14 | 56 | 228 | 0 | 71 | 67 | 2192 | 0 |

**Supplemental Table 33.** Paired t-tests for comparing the mean presyn-postsyn functional similarity between observation in the MICrONS dataset and values expected by GLMMs fit on the MICrONS dataset.

| Projection type | Comparison | t-statistic | p-value | adjusted p-value |
| --- | --- | --- | --- | --- |
| $HVA \rightarrow HVA$ | observed vs expected | 660.0 | 7.92e-01 | 9.81e-01 |
| $HVA \rightarrow V1$ | observed vs expected | 362.0 | 9.09e-01 | 9.81e-01 |
| $V1 \rightarrow HVA$ | observed vs expected | 62.0 | 5.17e-01 | 9.81e-01 |
| $V1 \rightarrow V1$ | observed vs expected | 313.0 | 9.81e-01 | 9.81e-01 |
| $HVA \rightarrow HVA$ | observed vs expected (synaptic scale) | 675.0 | 8.99e-01 | 9.81e-01 |
| $HVA \rightarrow V1$ | observed vs expected (synaptic scale) | 349.0 | 7.63e-01 | 9.81e-01 |
| $V1 \rightarrow HVA$ | observed vs expected (synaptic scale) | 71.0 | 8.18e-01 | 9.81e-01 |
| $V1 \rightarrow V1$ | observed vs expected (synaptic scale) | 280.0 | 5.76e-01 | 9.81e-01 |
| $HVA \rightarrow HVA$ | observed vs expected (axonal scale) | 629.0 | 5.85e-01 | 9.81e-01 |
| $HVA \rightarrow V1$ | observed vs expected (axonal scale) | 349.0 | 7.63e-01 | 9.81e-01 |
| $V1 \rightarrow HVA$ | observed vs expected (axonal scale) | 63.0 | 5.48e-01 | 9.81e-01 |
| $V1 \rightarrow V1$ | observed vs expected (axonal scale) | 284.0 | 6.22e-01 | 9.81e-01 |

**Supplemental Table 34.** Paired t-tests for comparing the mean postsyn-postsyn functional similarity between observation in the MICrONS dataset and values expected by GLMMs fit on the MICrONS dataset.

| Projection type | Comparison | t-statistic | p-value | adjusted p-value |
| --- | --- | --- | --- | --- |
| *HVA* → *HVA* | observed vs expected | 254.0 | 7.45e-05 | 1.79e-04 |
| *HVA* → *V1* | observed vs expected | 46.0 | 1.39e-07 | 5.56e-07 |
| *V1* → *HVA* | observed vs expected | 53.0 | 2.84e-01 | 2.84e-01 |
| *V1* → *V1* | observed vs expected | 45.0 | 9.74e-07 | 2.92e-06 |
| *HVA* → *HVA* | observed vs expected (synaptic scale) | 344.0 | 1.68e-03 | 2.52e-03 |
| *HVA* → *V1* | observed vs expected (synaptic scale) | 125.0 | 2.00e-04 | 3.99e-04 |
| *V1* → *HVA* | observed vs expected (synaptic scale) | 32.0 | 3.48e-02 | 4.64e-02 |
| *V1* → *V1* | observed vs expected (synaptic scale) | 197.0 | 5.35e-02 | 6.42e-02 |
| *HVA* → *HVA* | observed vs expected (axonal scale) | 344.0 | 1.68e-03 | 2.52e-03 |
| *HVA* → *V1* | observed vs expected (axonal scale) | 19.0 | 2.23e-09 | 1.34e-08 |
| *V1* → *HVA* | observed vs expected (axonal scale) | 53.0 | 2.84e-01 | 2.84e-01 |
| *V1* → *V1* | observed vs expected (axonal scale) | 5.0 | 5.82e-10 | 6.98e-09 |

## Bibliography

D. D. Bock, W.-C. A. Lee, A. M. Kerlin, M. L. Andermann, G. Hood, A. W. Wetzel, S. Yurgenson, E. R. Soucy, H. S. Kim, and R. C. Reid. Network anatomy and in vivo physiology of visual cortical neurons. Nature, 471(7337):177–182, Mar. 2011.

V. Braitenberg and A. Schüz. Cortex: Statistics and geometry of neuronal connectivity. Springer Science & Business Media, Mar. 2013.

S. A. Cadena, G. H. Denfield, E. Y. Walker, L. A. Gatys, A. S. Tolias, M. Bethge, and A. S. Ecker. Deep convolutional models improve predictions of macaque v1 responses to natural images. PLoS computational biology, 15(4):e1006897, 2019.

B. Celii, S. Papadopoulos, Z. Ding, P. G. Fahey, E. Wang, C. Papadopoulos, A. B. Kunin, S. Patel, J. Alexander Bae, A. L. Bodor, D. Brittain, J. Buchanan, D. J. Bumbarger, M. A. Castro, E. Cobos, S. Dorkenwald, L. Elabbady, A. Halageri, Z. Jia, C. Jordan, D. Kapner, N. Kemnitz, S. Kinn, K. Lee, K. Li, R. Lu, T. Macrina, G. Mahalingam, E. Mitchell, S. S. Mondal, S. Mu, B. Nehoran, S. Popovych, C. M. Schneider-Mizell, W. Silversmith, M. Takeno, R. Torres, N. L. Turner, W. Wong, J. Wu, S.-C. Yu, W. Yin, D. Xenes, L. M. Kitchell, P. K. Rivlin, V. A. Rose, C. A. Bishop, B. Wester, E. Froudarakis, E. Y. Walker, F. Sinz, H. Sebastian Seung, F. Collman, N. M. da Costa, R. Clay Reid, X. Pitkow, A. S. Tolias, and J. Reimer. NEURD: automated proofreading and feature extraction for connectomics. bioRxiv, Apr. 2024. URL https://www.biorxiv.org/content/10.1101/2023.03.14.532674v4.

L. Cossell, M. F. Iacaruso, D. R. Muir, R. Houlton, E. N. Sader, H. Ko, S. B. Hofer, and T. D. Mrsic-Flogel. Functional organization of excitatory synaptic strength in primary visual cortex. Nature, 518(7539):399–403, 2015.

S. Dorkenwald, C. E. McKellar, T. Macrina, N. Kemnitz, K. Lee, R. Lu, J. Wu, S. Popovych, E. Mitchell, B. Nehoran, Z. Jia, J. A. Bae, S. Mu, D. Ih, M. Castro, O. Ogedengbe, A. Halageri, K. Kuehner, A. R. Sterling, Z. Ashwood, J. Zung, D. Brittain, F. Collman, C. Schneider-Mizell, C. Jordan, W. Silversmith, C. Baker, D. Deutsch, L. Encarnacion-Rivera, S. Kumar, A. Burke, D. Bland, J. Gager, J. Hebditch, S. Koolman, M. Moore, S. Morejohn, B. Silverman, K. Willie, R. Willie, S.-C. Yu, M. Murthy, and H. S. Seung. FlyWire: online community for whole-brain connectomics. Nat. Methods, 19(1):119–128, Jan. 2022a.

S. Dorkenwald, N. L. Turner, T. Macrina, K. Lee, R. Lu, J. Wu, A. L. Bodor, A. A. Bleckert, D. Brittain, N. Kemnitz, W. M. Silversmith, D. Ih, J. Zung, A. Zlateski, I. Tartavull, S.-C. Yu, S. Popovych, W. Wong, M. Castro, C. S. Jordan, A. M. Wilson, E. Froudarakis, J. Buchanan, M. M. Takeno, R. Torres, G. Mahalingam, F. Collman, C. M. Schneider-Mizell, D. J. Bumbarger, Y. Li, L. Becker, S. Suckow, J. Reimer, A. S. Tolias, N. Macarico da Costa, R. C. Reid, and H. S. Seung. Binary and analog variation of synapses between cortical pyramidal neurons. Elife, 11, Nov. 2022b.

L. Elabbady, S. Seshamani, S. Mu, G. Mahalingam, C. Schneider-Mizell, A. L. Bodor, J. Alexander Bae, D. Brittain, J. Buchanan, D. J. Bumbarger, M. A. Castro, S. Dorkenwald, A. Halageri, Z. Jia, C. Jordan, D. Kapner, N. Kemnitz, S. Kinn, K. Lee, K. Li, R. Lu, T. Macrina, E. Mitchell, S. S. Mondal, B. Nehoran, S. Popovych, W. Silversmith, M. Takeno, R. Torres, N. L. Turner, W. Wong, J. Wu, W. Yin, S.-C. Yu, The MICrONS Consortium, H. Sebastian Seung, R. Clay Reid, N. M. Da Costa, and F. Collman. Perisomatic features enable efficient and dataset wide Cell-Type classifications across Large-Scale electron microscopy volumes. bioRxiv, Jan. 2024. URL https://www.biorxiv.org/content/10.1101/2022.07.20.499976v2.

P. G. Fahey, T. Muhammad, C. Smith, E. Froudarakis, E. Cobos, J. Fu, E. Y. Walker, D. Yatsenko, F. H. Sinz, J. Reimer, and A. S. Tolias. A global map of orientation tuning in mouse visual cortex. bioRxiv, Aug. 2019. URL https://www.biorxiv.org/content/10.1101/745323v1.

E. Froudarakis, P. Berens, A. S. Ecker, R. J. Cotton, F. H. Sinz, D. Yatsenko, P. Saggau, M. Bethge, and A. S. Tolias. Population code in mouse V1 facilitates readout of natural scenes through increased sparseness. Nat. Neurosci., 17(6):851–857, June 2014.

M. E. Garrett, I. Nauhaus, J. H. Marshel, and E. M. Callaway. Topography and areal organization of mouse visual cortex. J. Neurosci., 34(37):12587–12600, Sept. 2014.

D. O. Hebb. The organization of behavior; a neuropsychological theory. A Wiley Book in Clinical Psychology, 62:78, 1949.

S. Holler, G. Köstinger, K. A. C. Martin, G. F. P. Schuhknecht, and K. J. Stratford. Structure and function of a neocortical synapse. Nature, 591(7848):111–116, Mar. 2021.

C. Holmgren, T. Harkany, B. Svennenfors, and Y. Zilberter. Pyramidal cell communication within local networks in layer 2/3 of rat neocortex. The Journal of physiology, 551(Pt 1):139–153, Aug. 2003. ISSN 0022-3751. doi: 10.1113/jphysiol.2003.044784. URL http://dx.doi.org/10.1113/jphysiol.2003.044784.

J. Hopfield. Neural networks and physical systems with emergent collective computational abilities. Proc. Natl. Acad. Sci. USA, 79(8):2554–2558, 1982.

D. H. Hubel and T. N. Wiesel. Receptive fields, binocular interaction and functional architecture in the cat’s visual cortex. J. Physiol., 160:106–154, Jan. 1962.

M. F. Iacaruso, I. T. Gasler, and S. B. Hofer. Synaptic organization of visual space in primary visual cortex. Nature, 547(7664):449–452, July 2017.

M. Khona and I. Fiete. Attractor and integrator networks in the brain. Nat. Rev. Neurosci., 23(12): 744–766, 2022a.

M. Khona and I. R. Fiete. Attractor and integrator networks in the brain. Nat. Rev. Neurosci., 23 (12):744–766, Dec. 2022b.

H. Ko, S. B. Hofer, B. Pichler, K. A. Buchanan, P.J. Sjöström, and T. D. Mrsic-Flogel. Functional specificity of local synaptic connections in neocortical networks. Nature, 473(7345):87–91, 2011.

H. Ko, L. Cossell, C. Baragli, J. Antolik, C. Clopath, S. B. Hofer, and T. D. Mrsic-Flogel. The emergence of functional microcircuits in visual cortex. Nature, 496(7443):96–100, Apr. 2013.

S. Kondo and K. Ohki. Laminar differences in the orientation selectivity of geniculate afferents in mouse primary visual cortex. Nat. Neurosci., 19(2):316–319, Feb. 2016.

A. T. Kuan, G. Bondanelli, L. N. Driscoll, J. Han, M. Kim, D. G. C. Hildebrand, B. J. Graham, D. E. Wilson, L. A. Thomas, S. Panzeri, C. D. Harvey, and W.-C. A. Lee. Synaptic wiring motifs in posterior parietal cortex support decision-making. Nature, Feb. 2024.

K. Lee, J. Zung, P. Li, V. Jain, and H. Sebastian Seung. Superhuman accuracy on the SNEMI3D connectomics challenge. arXiv, May 2017.

W.-C. A. Lee, V. Bonin, M. Reed, B. J. Graham, G. Hood, K. Glattfelder, and R. C. Reid. Anatomy and function of an excitatory network in the visual cortex. Nature, 532(7599):370–374, Apr. 2016.

R. Lu, A. Zlateski, and H. Sebastian Seung. Large-scale image segmentation based on distributed clustering algorithms. arXiv, June 2021.

T. Marques, J. Nguyen, G. Fioreze, and L. Petreanu. The functional organization of cortical feedback inputs to primary visual cortex. Nat. Neurosci., 21(5):757–764, May 2018.

A. Mathis, P. Mamidanna, K. M. Cury, T. Abe, V. N. Murthy, M. W. Mathis, and M. Bethge. DeepLabCut: markerless pose estimation of user-defined body parts with deep learning. Nat. Neurosci., 21(9):1281–1289, Sept. 2018.

M. Mazurek, M. Kager, and S. D. Van Hooser. Robust quantification of orientation selectivity and direction selectivity. Front. Neural Circuits, 8:92, Aug. 2014.

MICrONS Consortium, J. Alexander Bae, M. Baptiste, A. L. Bodor, D. Brittain, J. Buchanan, D. J. Bumbarger, M. A. Castro, B. Celii, E. Cobos, F. Collman, N. M. da Costa, S. Dorkenwald, L. Elabbady, P. G. Fahey, T. Fliss, E. Froudakis, J. Gager, C. Gamlin, A. Halageri, J. Hebditch, Z. Jia, C. Jordan, D. Kapner, N. Kemnitz, S. Kinn, S. Koolman, K. Kuehner, K. Lee, K. Li, R. Lu, T. Macrina, G. Mahalingam, S. McReynolds, E. Miranda, E. Mitchell, S. S. Mondal, M. Moore, S. Mu, T. Muhammad, B. Nehoran, O. Ogedengbe, C. Papadopoulos, S. Papadopoulos, S. Patel, X. Pitkow, S. Popovych, A. Ramos, R. Clay Reid, J. Reimer, C. M. Schneider-Mizell, H. Sebastian Seung, B. Silverman, W. Silversmith, A. Sterling, F. H. Sinz, C. L. Smith, S. Suckow, Z. H. Tan, A. S. Tolias, R. Torres, N. L. Turner, E. Y. Walker, T. Wang, G. Williams, S. Williams, K. Willie, R. Willie, W. Wong, J. Wu, C. Xu, R. Yang, D. Yatsenko, F. Ye, W. Yin, and S.-C. Yu. Functional connectomics spanning multiple areas of mouse visual cortex. bioRxiv, July 2021. URL https://www.biorxiv.org/content/10.1101/2021.07.28.454025v3.

E. Mitchell, S. Keselj, S. Popovych, D. Buniatyan, and H. Sebastian Seung. Siamese encoding and alignment by multiscale learning with Self-Supervision. arXiv, Apr. 2019.

S. Musall, M. T. Kaufman, A. L. Juavinett, S. Gluf, and A. K. Churchland. Single-trial neural dynamics are dominated by richly varied movements. Nat. Neurosci., 22(10):1677–1686, Oct. 2019.

I. A. Oldenburg, W. D. Hendricks, G. Handy, K. Shamardani, H. A. Bounds, B. Doiron, and H. Adesnik. The logic of recurrent circuits in the primary visual cortex. Nature neuroscience, 27(1):137–147, Jan. 2024. ISSN 1097-6256. doi: 10.1038/s41593-023-01510-5. URL https://www.nature.com/articles/s41593-023-01510-5.

A. Peters and M. L. Feldman. The projection of the lateral geniculate nucleus to area 17 of the rat cerebral cortex. I. general description. J. Neurocytol., 5(1):63–84, Feb. 1976.

J. S. Phelps, D. G. C. Hildebrand, B. J. Graham, A. T. Kuan, L. A. Thomas, T. M. Nguyen, J. Buhmann, A. W. Azevedo, A. Sustar, S. Agrawal, M. Liu, B. L. Shanny, J. Funke, J. C. Tuthill, and W.-C. A. Lee. Reconstruction of motor control circuits in adult drosophila using automated transmission electron microscopy. Cell, 184(3):759–774.e18, Feb. 2021.

S. Ramón y Cajal. Histologie du système nerveux de l’homme et des vertébrés. 1911.

C. L. Rees, K. Moradi, and G. A. Ascoli. Weighing the evidence in peters’ rule: Does neuronal morphology predict connectivity?, Feb. 2017.

R. C. Reid. From functional architecture to functional connectomics. Neuron, 75(2):209–217, July 2012.

J. Reimer, E. Froudarakis, C. R. Cadwell, D. Yatsenko, G. H. Denfield, and A. S. Tolias. Pupil fluctuations track fast switching of cortical states during quiet wakefulness. Neuron, 84(2): 355–362, Oct. 2014.

D. L. Ringach, P. J. Mineault, E. Tring, N. D. Olivas, P. Garcia-Junco-Clemente, and J. T. Trachtenberg. Spatial clustering of tuning in mouse primary visual cortex. Nat. Commun., 7:12270, Aug. 2016.

L. F. Rossi, K. D. Harris, and M. Carandini. Spatial connectivity matches direction selectivity in visual cortex. Nature, 588(7839):648–652, Dec. 2020.

C. Schneider-Mizell and F. Collman. Alleninstitute/pcg_skel: v0.3.2, Mar. 2023. URL 10.5281/zenodo.7703278.

C. M. Schneider-Mizell, A. L. Bodor, D. Brittain, J. Buchanan, D. J. Bumbarger, L. Elabbady, C. Gamlin, D. Kapner, S. Kinn, G. Mahalingam, S. Seshamani, S. Suckow, M. Takeno, R. Torres, W. Yin, S. Dorkenwald, J. A. Bae, M. A. Castro, A. Halageri, Z. Jia, C. Jordan, N. Kemnitz, K. Lee, K. Li, R. Lu, T. Macrina, E. Mitchell, S. S. Mondal, S. Mu, B. Nehoran, S. Popovych, W. Silversmith, N. L. Turner, W. Wong, J. Wu, MICrONS Consortium, J. Reimer, A. S. Tolias, H. S. Seung, R. C. Reid, F. Collman, and N. Maçarico da Costa. Cell-type-specific inhibitory circuitry from a connectomic census of mouse visual cortex. bioRxiv, Jan. 2024. URL https://www.biorxiv.org/content/10.1101/2023.01.23.525290v3.

B. Scholl, C. I. Thomas, M. A. Ryan, N. Kamasawa, and D. Fitzpatrick. Cortical response selectivity derives from strength in numbers of synapses. Nature, 590(7844):111–114, Feb. 2021.

O. Schoppe, N. S. Harper, B. D. B. Willmore, A. J. King, and J. W. H. Schnupp. Measuring the performance of neural models. Frontiers in Computational Neuroscience, 10, Feb. 2016. doi: 10.3389/fncom.2016.00010.

N. J. Sofroniew, D. Flickinger, J. King, and K. Svoboda. A large field of view two-photon mesoscope with subcellular resolution for in vivo imaging. Elife, 5, June 2016.

C. Stringer, M. Pachitariu, N. Steinmetz, C. B. Reddy, M. Carandini, and K. D. Harris. Spontaneous behaviors drive multidimensional, brainwide activity. Science, 364(6437):255, Apr. 2019.

R. Tong, R. da Silva, D. Lin, A. Ghosh, J. Wilsenach, E. Cianfarano, P. Bashivan, B. Richards, and S. Trenholm. The feature landscape of visual cortex. bioRxiv, 2023. doi: 10.1101/2023.11.03.565500. URL https://www.biorxiv.org/content/early/2023/11/05/2023.11.03.565500.

N. L. Turner, K. Lee, R. Lu, J. Wu, D. Ih, and H. S. Seung. Synaptic partner assignment using attentional voxel association networks. 2020 IEEE 17th International Symposium on Biomedical Imaging (ISBI), pages 1–5, Apr. 2020.

P. Virtanen, R. Gommers, T. E. Oliphant, M. Haberland, T. Reddy, D. Cournapeau, E. Burovski, P. Peterson, W. Weckesser, J. Bright, S. J. van der Walt, M. Brett, J. Wilson, K. J. Millman, N. Mayorov, A. R. J. Nelson, E. Jones, R. Kern, E. Larson, C. J. Carey Polat,, Y. Feng, E. W. Moore, J. VanderPlas, D. Laxalde, J. Perktold, R. Cimrman, I. Henriksen, E. A. Quintero, C. R. Harris, A. M. Archibald, A. H. Ribeiro, F. Pedregosa, P. van Mulbregt, and SciPy 1.0 Contributors. SciPy 1.0: fundamental algorithms for scientific computing in python. Nat. Methods, 17(3):261–272, Mar. 2020.

E. Y. Walker, F. H. Sinz, E. Cobos, T. Muhammad, E. Froudarakis, P. G. Fahey, A. S. Ecker, J. Reimer, X. Pitkow, and A. S. Tolias. Inception loops discover what excites neurons most using deep predictive models. Nat. Neurosci., 22(12):2060–2065, Dec. 2019.

E. Y. Wang, P. G. Fahey, Z. Ding, S. Papadopoulos, K. Ponder, M. A. Weis, A. Chang, T. Muhammad, S. Patel, Z. Ding, D. Tran, J. Fu, C. M. Schneider-Mizell, R. C. Reid, F. Collman, N. M. da Costa, K. Franke, A. S. Ecker, J. Reimer, X. Pitkow, F. H. Sinz, and A. S. Tolias. Foundation model of neural activity predicts response to new stimulus types and anatomy. bioRxiv, Aug. 2024. URL https://www.biorxiv.org/content/10.1101/2023.03.21.533548v4.

Q. Wang and A. Burkhalter. Area map of mouse visual cortex. J. Comp. Neurol., 502(3):339–357, May 2007.

A. A. Wanner, C. Genoud, T. Masudi, L. Siksou, and R. W. Friedrich. Dense EM-based reconstruction of the interglomerular projectome in the zebrafish olfactory bulb. Nat. Neurosci., 19 (6):816–825, June 2016.

M. A. Weis, S. Papadopoulos, L. Hansel, T. Lüddecke, B. Celii, P. G. Fahey, E. Y. Wang, J. A. Bae, A. L. Bodor, D. Brittain, J. Buchanan, D. J. Bumbarger, M. A. Castro, F. Collman, N. M. da Costa, S. Dorkenwald, L. Elabbady, A. Halageri, Z. Jia, C. Jordan, D. Kapner, N. Kemnitz, S. Kinn, K. Lee, K. Li, R. Lu, T. Macrina, G. Mahalingam, E. Mitchell, S. S. Mondal, S. Mu, B. Nehoran, S. Popovych, R. C. Reid, C. M. Schneider-Mizell, H. S. Seung, W. Silversmith, M. Takeno, R. Torres, N. L. Turner, W. Wong, J. Wu, W. Yin, S.-C. Yu, J. Reimer, P. Berens, A. S. Tolias, and A. S. Ecker. An unsupervised map of excitatory neurons’ dendritic morphology in the mouse visual cortex. bioRxiv, Apr. 2024. URL https://www.biorxiv.org/content/10.1101/2022.12.22.521541v3.

A. Wertz, S. Trenholm, K. Yonehara, D. Hillier, Z. Raics, M. Leinweber, G. Szalay, A. Ghanem, G. Keller, B. Rózsa, K.-K. Conzelmann, and B. Roska. PRESYNAPTIC NETWORKS. singlecell-initiated monosynaptic tracing reveals layer-specific cortical network modules. Science, 349(6243):70–74, July 2015.

J. Wu, W. M. Silversmith, K. Lee, and H. S. Seung. Chunkflow: hybrid cloud processing of large 3D images by convolutional nets. Nat. Methods, 18(4):328–330, Apr. 2021.

D. Yamins, H. Hong, C. Cadieu, E. A. Solomon, D. Seibert, and J. DiCarlo. Performance-optimized hierarchical models predict neural responses in higher visual cortex. Proc. Natl. Acad. Sci. USA, 111(23):8619–8624, 2014.

W. Yin, D. Brittain, J. Borseth, M. E. Scott, D. Williams, J. Perkins, C. S. Own, M. Murfitt, R. M. Torres, D. Kapner, G. Mahalingam, A. Bleckert, D. Castelli, D. Reid, W.-C. A. Lee, B. J. Graham, M. Takeno, D. J. Bumbarger, C. Farrell, R. C. Reid, and N. M. da Costa. A petascale automated imaging pipeline for mapping neuronal circuits with high-throughput transmission electron microscopy. Nat. Commun., 11(1):4949, Oct. 2020.

Z. Yu, M. Guindani, S. F. Grieco, L. Chen, T. C. Holmes, and X. Xu. Beyond t test and ANOVA: applications of mixed-effects models for more rigorous statistical analysis in neuroscience research. Neuron, 110(1):21–35, Jan. 2022.

P. Znamenskiy, M.-H. Kim, D. R. Muir, M. F. Iacaruso, S. B. Hofer, and T. D. Mrsic-Flogel. Functional specificity of recurrent inhibition in visual cortex. Neuron, 112(6):991–1000.e8, Mar. 2024.

